# Graph Embedding Method Based Genetical Trajectory Reveals Migration History Among East Asians

**DOI:** 10.1101/870253

**Authors:** Zhuang Wei, Ching-Wen Chang, Van Luo, Beilei Bian, Xuewei Ding

## Abstract

An important issue in human population genetics is the ancestry. By extracting the ancestral information retained in the single nucleotide polymorphism (SNP) of genomic DNA, the history of migration and reproduction of the population can be reconstructed. Since the SNP data of population are multidimensional, their dimensionality reduction can demonstrate their potential internal connections. In this study, the graph and structure learning based Graph Embedding method commonly used in single cell mRNA sequencing was applied to human population genetics research to decrease the data dimension. As a result, the human population trajectory of East Asia based on 1000 Genomes Project was reconstructed to discover the inseparable relationship between the Chinese population and other East Asian populations. These results are visualized from various ancestry calculators such as E11 and K12B. Finally, the unique SNPs along the psudotime of trajectory were found by differential analysis. Bioprocess enrichment analysis was also used to reveal that the genes of these SNPs may be related to neurological diseases. These results will lay the data foundation for precision medicine.

## Introduction

As an organism, DNA sequences store human genetic information. The genetic information of DNA is mainly inherited from parents, following the population genetic laws of diploid organisms, and random mutations of certain probability occur at any time (Tishkoff and Verrelli, 2003). In the long history of human evolution, certain types of DNA mutation (single nucleotide polymorphism) have created the diversity of human ancestry (Garrigan and Hammer, 2006). Due to the relatively stable alteration rate, those mutations can reflect the history of human evolution (Cavalli-Sforza and Feldman, 2003). Therefore, trajectory the of human ancestry can be restored by studying the modern human populations.

After the human genomic DNA sequencing data was removed from the unstable mutations and mutations do not reflect human evolution, the remaining mutation table is still a high-dimensional data containing the genetic information of the research population (Shendure, 2011; Tian et al., 2008). Through PCA, tSNE and UMAP, high-dimensional data can be reduced to two-dimensional table with efficient information, which can reflect the far-near relationships between genetic groups (Diaz-Papkovich et al., 2019; Gaspar and Breen, 2019; Lorenzo et al., 2019). However, the above dimension reduction methods do not give a very effective way for revealing the connection between groups. Traditional methods for identifying the internal structure of high-dimensional data include principal curve, elastic map and principal graph, etc., but these methods only give the characteristics of the data itself and do not have a high level learning process (Mao et al., 2015b; Qi et al., 2017; Zhang et al., 2011). Gaspar et al. studied the population structure of 1000 Genomes Project data by using GTM-based methods (Gaspar and Breen, 2019). The GTM method has certain machine learning properties and can self-organize the topology hidden inside the data, but still cannot reflect the complex connections within the data (Gaspar and Breen, 2019). Therefore, there is an urgent task to find a method to reduce the dimension of genetic data, which is more able to reflect the detailed internal connections between groups.

East Asia is one of the birthplaces of human culture (Su et al., 1999). Especially the ancient Yellow River Basin civilization and the large-scale agricultural society form fostered by the Yangtze River Basin have laid an irreplaceable material and cultural foundation for the development of civilization in East Asia. The early Chinese cultural model has greatly influenced the cultural patterns of the Japanese islands, the Korean peninsula and the Vietnamese peninsula, as well as parts of the Southeast Asia mainland. The vocabulary derived from classical Chinese has profoundly changed the language systems of these regions. However, the genetic landscape of this area, as well as the early migration of humans in East Asia, are still unclear. Based on historical records and information provided by archaeological literature, we can learn many clues that can be used for limited understanding, but due to historical records and non-scientific, almost “mythological” descriptions, many recorded “legends” seem to account for bigger. Some Asian human paleontology papers published in recent years have given very important information in this field (Su et al., 1999), but the dynamic migration process of the overall population structure is still very vague. Therefore, it is necessary to explore population migration patterns based on internal connections of population.

Our research used the combination of the principle graph and structure learning based Graph Embedding method and tSNE to reduce data dimension. Based on the R package “Monocle” we first filtered the mutations, then reduced the data dimension, and calculated trajectory with psudotime and states; we also adopted all the equilibrium mutations to tSNE method firstly to reduce the dimension, and then uses DDRTree to reduce the dimension twice. The E11 calculator and the K12B calculator verify the sampling results of the 1000 Genomes Project. At the same time, we found the mutations drifting trend along psudotime in the East Asian population from south to north, and found the genes corresponding to these mutations are significantly enriched in certain signaling pathways.

## Result

Previous studies have shown that the simple tSNE, PCA or GTM method do not distinguish the dataset from the 1000 Genomes Project Phase III well within the Asian group (Gaspar and Breen, 2019). Therefore, we use the principal graph and structure learning-based Graph Embedding method, which currently used in the single cell field to process the data preprocessed in the paper (Gaspar and Breen, 2019). Technically, we adopt two strategies, Fig 1A. One is to process the data using the R package “Monocle” that based on Graph Embedding method (Qiu et al., 2017a; Qiu et al., 2017b; Trapnell et al., 2014). Another one is the initial dimension reduction using tSNE and the second dimension reduction using DDRTree method, we call it tSNE + DDRTree.

**Fig 1.**
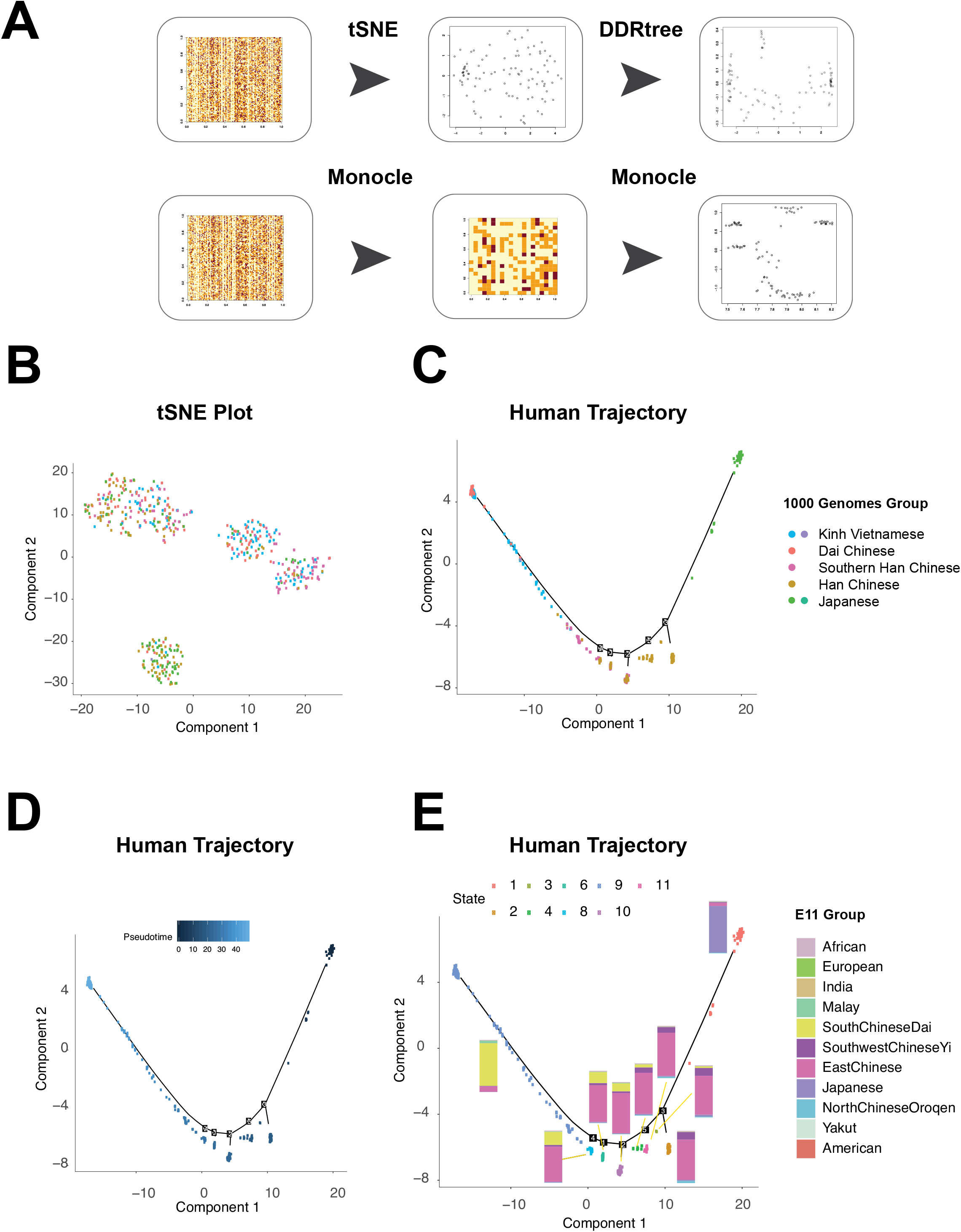
A. Principles of data analysis. B. tSNE dimension reduction of East Asian population of Genome 1000 program data. Each point indicated one individual, plots color indicated the area group from Genome 1000 program. C. Reversed graph embedding method learned Human trajectory are divided into two main groups by using Monocle package. D. Pseudotime, E. Lineage states, the bar chart indicated the population components predicted by E11 calculator. Bar color indicated the E11 components.

Using the “Monocle” package, we obtained SNP loci with large coefficient of variation. Using these SNP loci, the integrated tSNE in “Monocle” for dimensionality reduction was performed. The results in Fig 1B show that East Asia data can be roughly divided into three groups mainly correspond to Japan, the Chinese Han and the Chinese Yi groups plus Vietnamese.

Further, we use the methods of DDRTree in the package, which can generate the trajectory based on Graph Embedding method, Fig 1C, D and E. In addition, we also get the psudotime and states information from the data, Fig 1D and E. By mapping the 1000 Genomes Project Phase III population information on this result, Fig 1C, we can roughly see that along the two ends of the branches of the trajectory are the Japanese and the Chinese Yi plus Vietnamese groups. Groups Between those two ends are the Han Chinese in the north and the Han Chinese in the south.

From the calculated psudotime we can also see the relative origin time that two groups were separated. From our result, Fig 1D, psudotime between the Japanese to the Chinese Han and the Chinese Yi plus Vietnamese the Chinese Han are similar, which indicated the same origin time.

To further confirm the results, data was analyzed one by one using the E11 calculator for the East Asian population, which is popular in the community, and it was averaged according to the states, Fig 1E. The EastChinease component is the main proportion of the nodes of the entire trajectory. The results of E11 calculations correspond to the end points of Japanese to the Chinese Han and the Chinese Yi plus Vietnamese groups, indicating the correctness of their branches. Dbscan can use density to classify two-dimensional points clusters. In this study, we also performed E11 analysis on dbscan classfied fine groups. As Fig 2 and 3 showed, there are rare E11 components, such as NorthChineseQroqen or Malay groups enriched subgroups.

**Fig 2.**
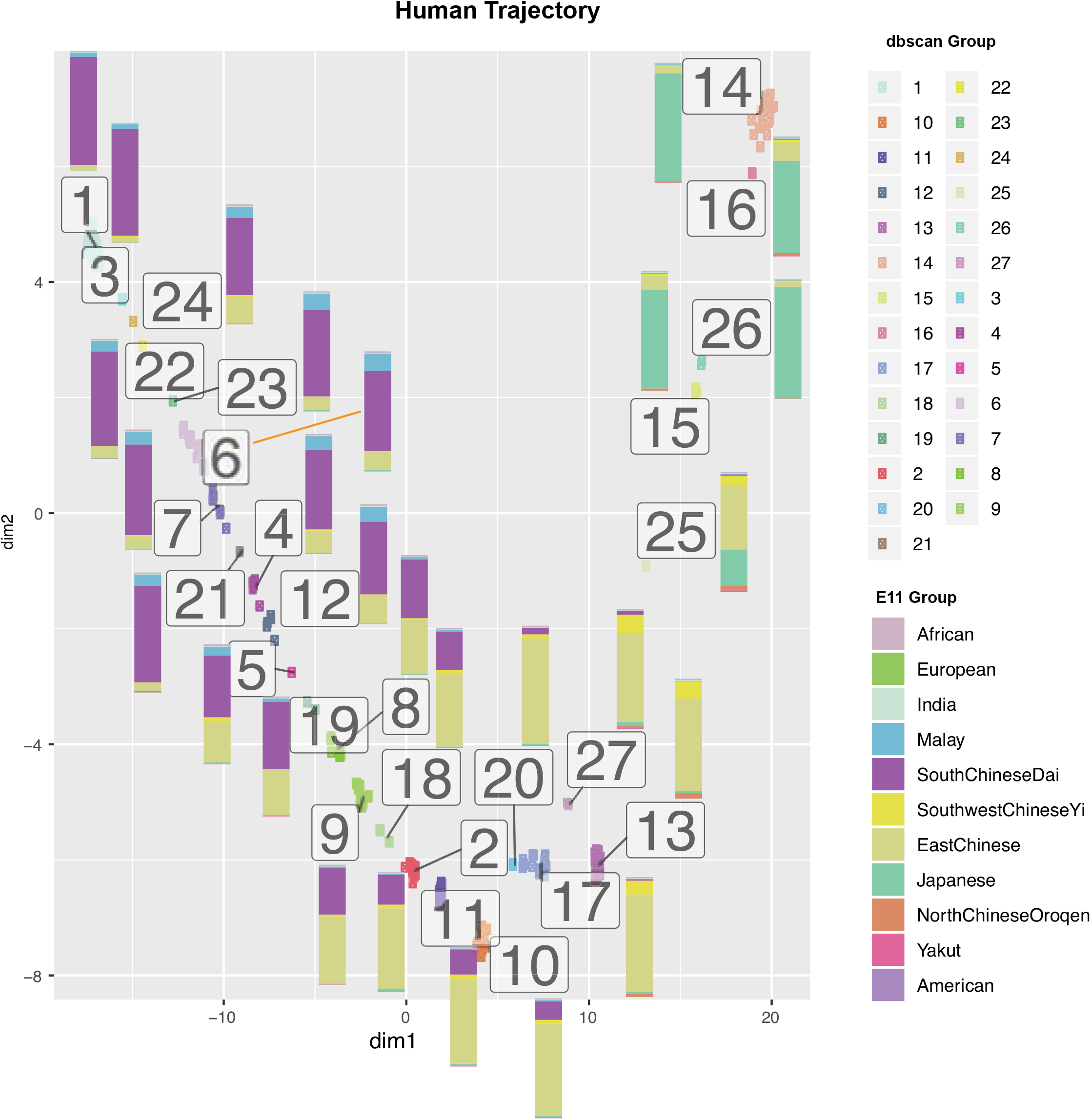
Groups were detected by dbscan method on Human trajectory. Plots color indicated the dbscan group. The bar chart indicated the population components predicted by E11 calculator. Bar color indicated the E11 components.

**Fig 3.**
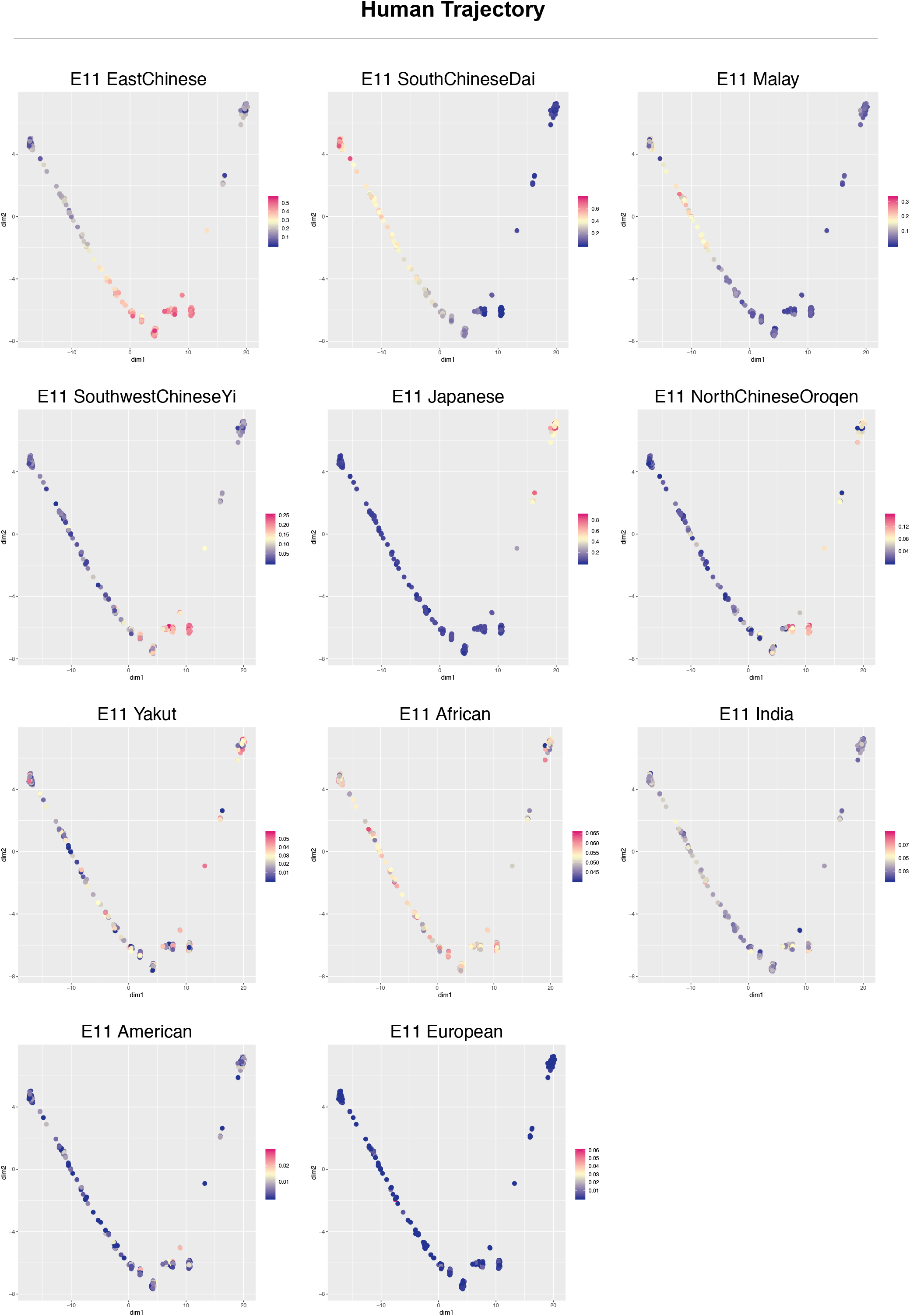
Reversed graph embedding method learned Human trajectory by using Monocle package. Each of E11 component was plotted on individual point by continuous color. Values were normalized by each E11 component.

In order to verify the results of E11 analysis, we used K12B calculator to confirm each individual in the dataset, and plot them on the trajectory, as shown in Fig 5. The results are geographically similar to the results displayed by the E11 calculator. It is worth mentioning that the groups of Malay enriched in E11 are basically coincident with the group enriched by K12B of South Asian, indicating their connection.

Using the tSNE+DDRTree method, we firstly performed tSNE dimensionality reduction on all the SNPs in the dataset, and further reduce the dimension using the DDRTree method. The results showed more branched trajectory map, as shown in Fig 4 and Fig 6. We can see that the E11 and K12B components in the branched are of higher purity than the groups “Monocle” package generated.

**Fig 4.**
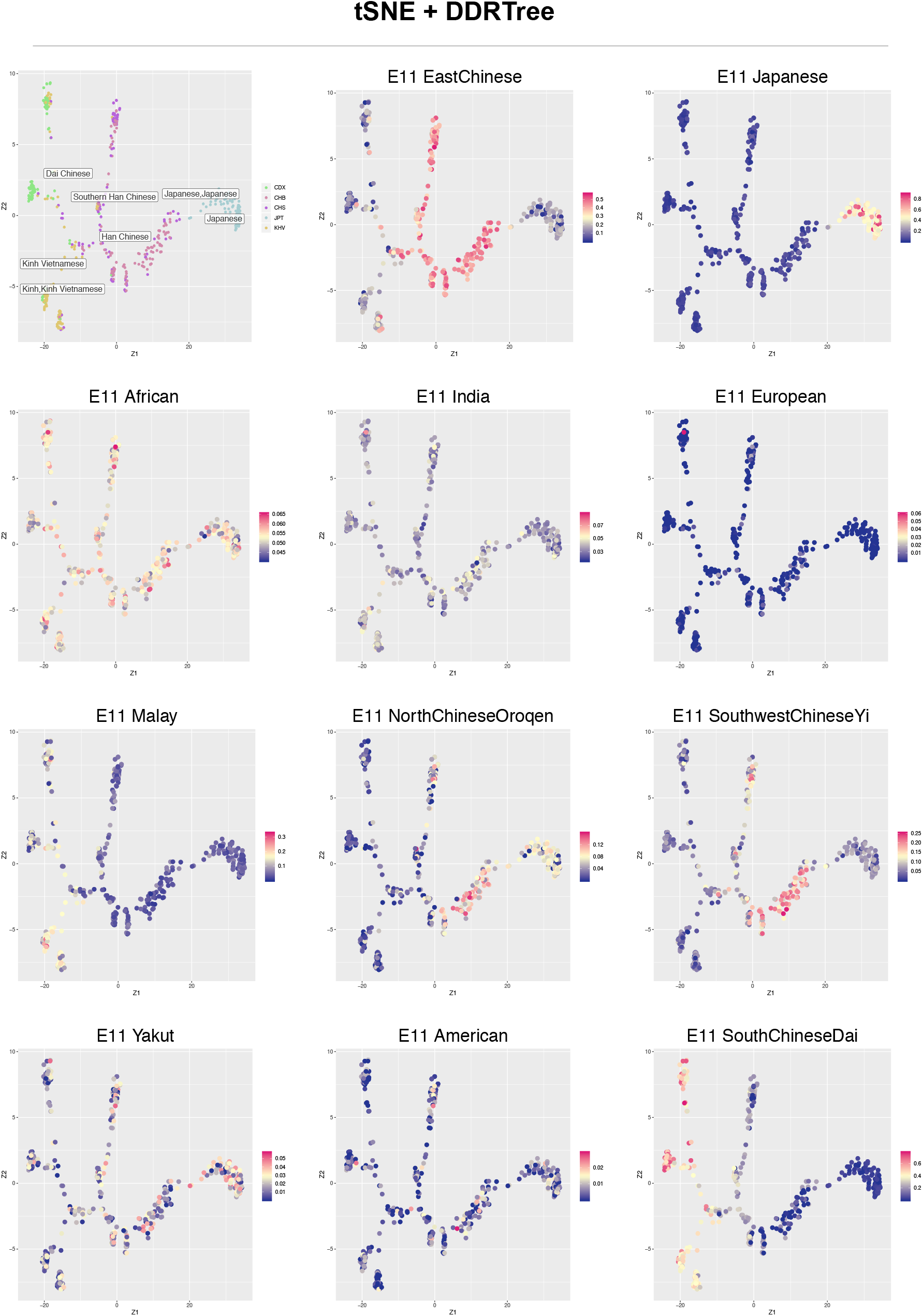
Reversed graph embedding method learned dimension reduction plot by using tSNE and DDRTree combined method. Each of E11 component was plotted on individual point by continuous color. Values were normalized by each E11 component.

Along the two ends of psudotime in trajectory, we can see that the difference in SNPs showed a gradual changing, Fig 7A, and the corresponding genes of the top 300 SNPs are analyzed for GO signal pathway function enrichment, and the results showed that most of the differential genes are related to membrane potentials or nervous system related pathways, such as morphine addiction, Fig7 B and C.

## Discussion

### Genetic interaction between Japanese and Chinese population

According to the theory put forward by archaeologists, the modern Japanese are new as a group who mixed Jomon hunter-gatherers during the 16kya period and the Yayoi people crossed the sea from the Korean peninsula during 400BC-1,200CE (Lee and Hasegawa, 2011; NANTA, 2008). Population DNA analysis also confirmed the conclusions of archaeological studies of two different cultural groups (Watanobe et al., 2004). Archaeologists believe that in the late Jomon period, temperatures and sea levels fell simultaneously, which had a negative impact on the hunter-gatherers of the archipelago, and the population of Jomon declined sharply (Crema et al., 2016). During this period, the Yayoi people migrated from the Korean peninsula or from northern China to the Japanese archipelago during the 2,400ya-1,700ya period (Iizuka and Nakahashi, 2002). These rice farmers are not only mixed with local hunter-gatherer groups, but also bring their rice-growing techniques and language systems to the early coastal farmers groups (Lee and Hasegawa, 2011). The influence of Yayoi on the genetics of people across Japan may vary. The Ainu people in the northernmost Hokkaido, as well as the Ryukyu people who live in the southernmost Ryukyu Islands, are less affected by the Yayoi people (Adachi et al., 2018). They retain more from the Jomons than the Japanese in other regions (Adachi et al., 2018; Crema et al., 2016).

Previous PCA analysis of DNA data of the Jomon people and the Japanese islanders, as well as the residents of various parts of China showed the Japanese and Chinese seem to be genetically far apart (Jinam et al., 2012). However, if we consider the island effect of the Japanese island, we must doubt the bias in these genetic analyses. Because the island effect may increase the rate of DNA change. These analyses may overestimate the distance between the Chinese and the Japanese.

Our results clearly show the close ties between Japanese and the Han Chinese in the north and the Han Chinese in the south and show the transitional population between these two groups, Fig 1, 2 and 3. It can be seen from our research that the E11 component shared by the Japanese and the Chinese are Oroqen in Northeast population of China. The Oroqen people in China are generally considered to be descendants of the indigenous people of Northeast Asia (Janhunen, 2005), and the K12B results also support this conclusion. In Fig 4, 5 and 6, the excessive group connected between the Japanese and the Chinese is also the higher component of Oroqen, which supported that the main group in Japan maybe the Northeast Asian immigrant from China and Korea peninsula.

**Fig 5.**
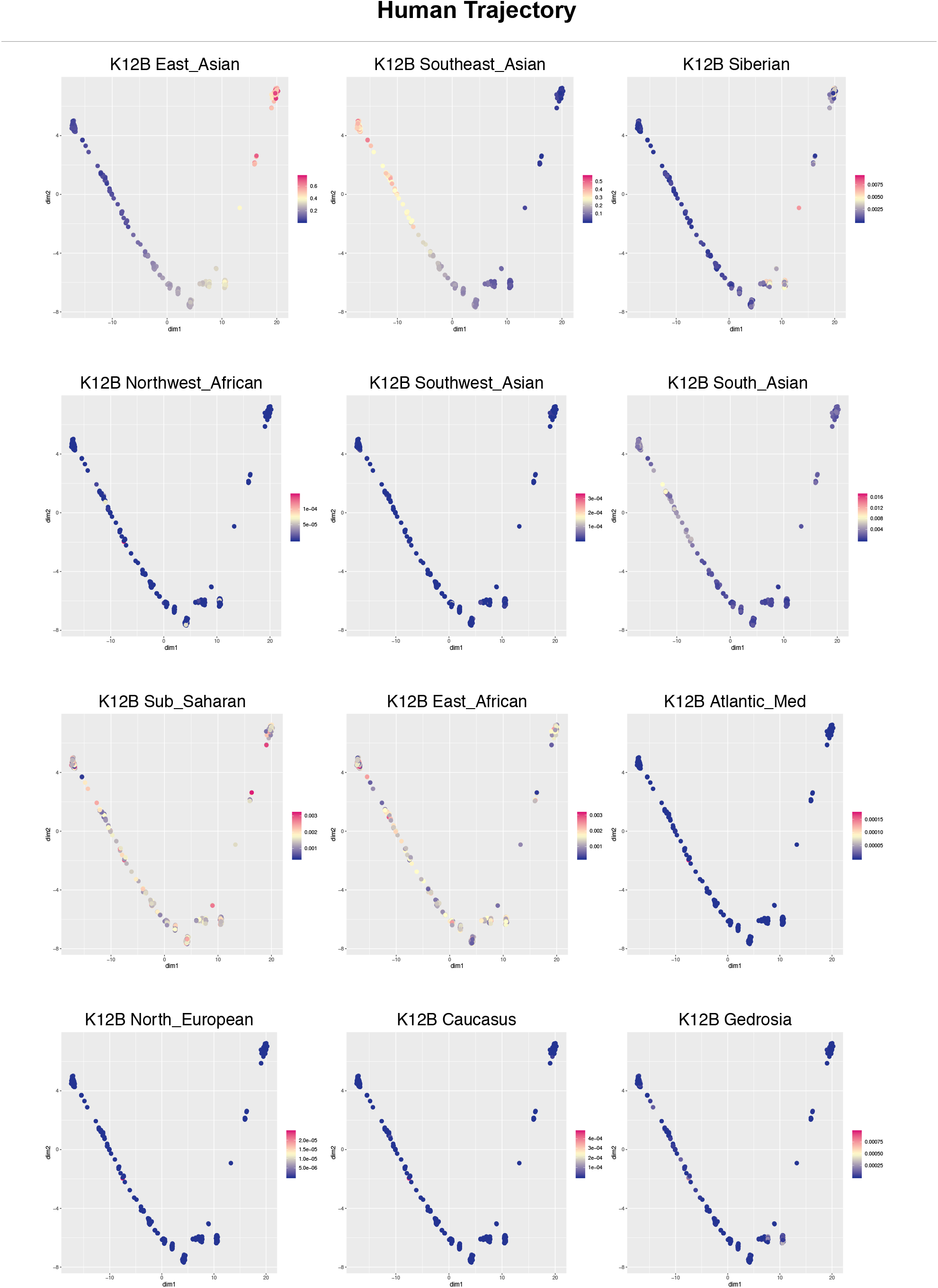
Reversed graph embedding method learned Human trajectory by using Monocle package. Each of K12B component was plotted on individual point by continuous color. Values were normalized by each K12B component.

**Fig 6.**
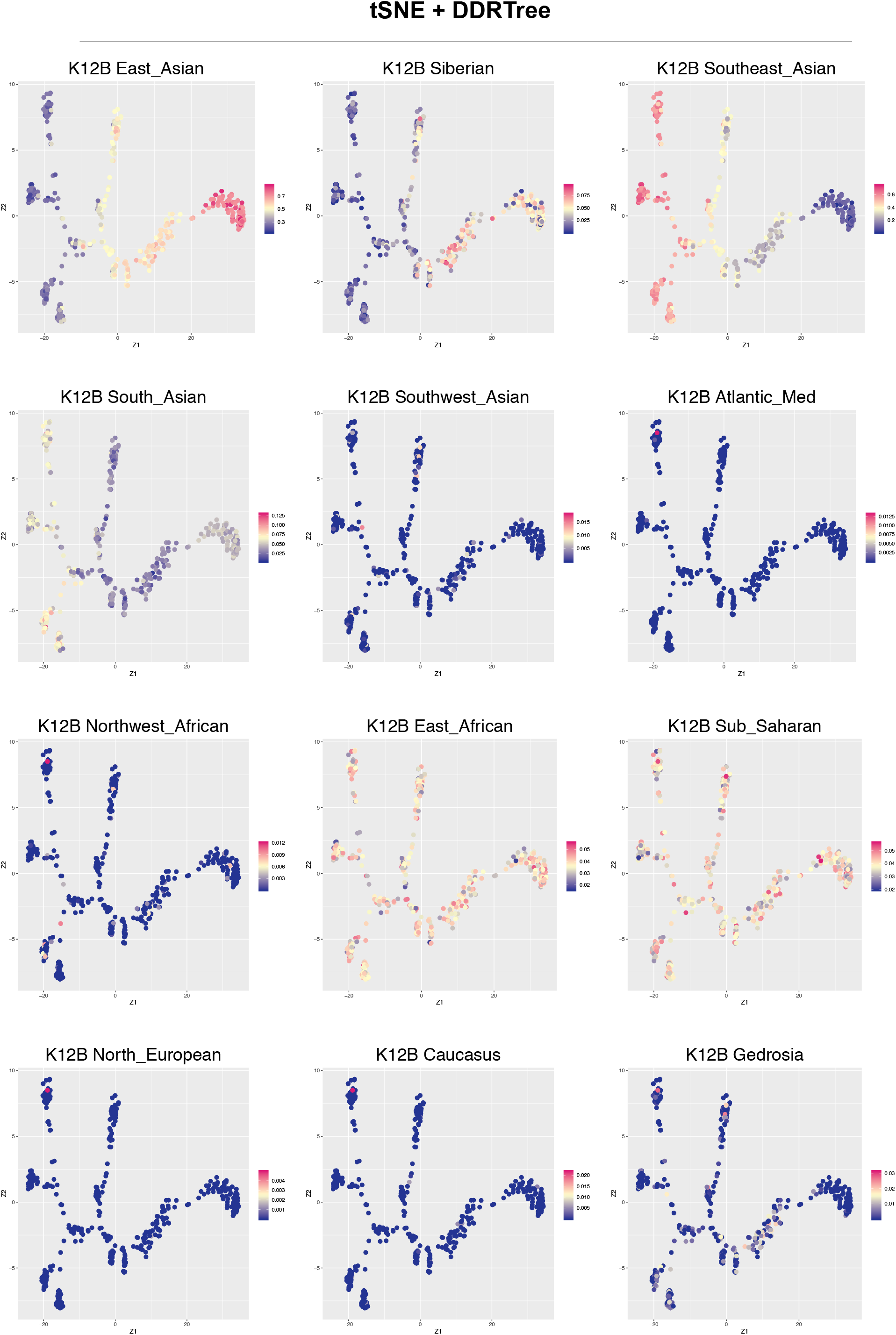
Reversed graph embedding method learned dimension reduction plot by using tSNE and DDRTree combined method. Each of K12B component was plotted on individual point by continuous color. Values were normalized by each K12B component.

### Genetic interaction between Vietnamese and Chinese population

Previous analysis of DNA samples from Hanoi, Vietnam, revealed that the Vietnamese in northern Vietnam are more likely to cluster with East Asians such as Chinese, Japanese and Korean peninsula (Kim et al., 2000). Compared with the Japanese, the Vietnamese and the Chinese share more genetic relationship (Le et al., 2019; Xia et al., 2019), especially with the South China population (He et al., 2012). These can be learned from historical literature. Vietnam has a millennial county or northern period. During this period, a large number of immigrant groups from North China occupied the dominance of Vietnam and originated from the Yangtze River valley in the late Neolithic period (Hanihara et al., 2012; Sun et al., 2013).

It is also worth noting that the genetic difference between the Taiyi group that entered the mainland of Southeast Asia in the 10th century and the Chinese Zhuang group in Guangxi was small. They clustered in the PCA analysis (Sun et al., 2013), and the Kinh and Thai people in the northern part of Vietnam connected to Guangxi in China. Groups have similar genomic structures and close evolutionary relationships (Le et al., 2019).

In our results, there is a close relationship between the Han Chinese in southern China and the Vietnamese, Fig 1, 2 and 3. The South Asian population of the E11-marked Malays also contributed to Vietnamese groups, which shows a great accordance with the previous studies, Fig 4, 5 and 6.

The genetic differentiation of the population is caused by genomic diversity (SNPs). In theory, these SNPs will gradually change in the associated groups. Using difference analysis, we found the differences of SNPs in different states along the trajectory can enriched in certain biological functions. The results show that the differences in SNPs in different states are mainly concentrated in biological processes related to nervous system functions such as morphine addiction, Fig 7.

**Fig 7.**
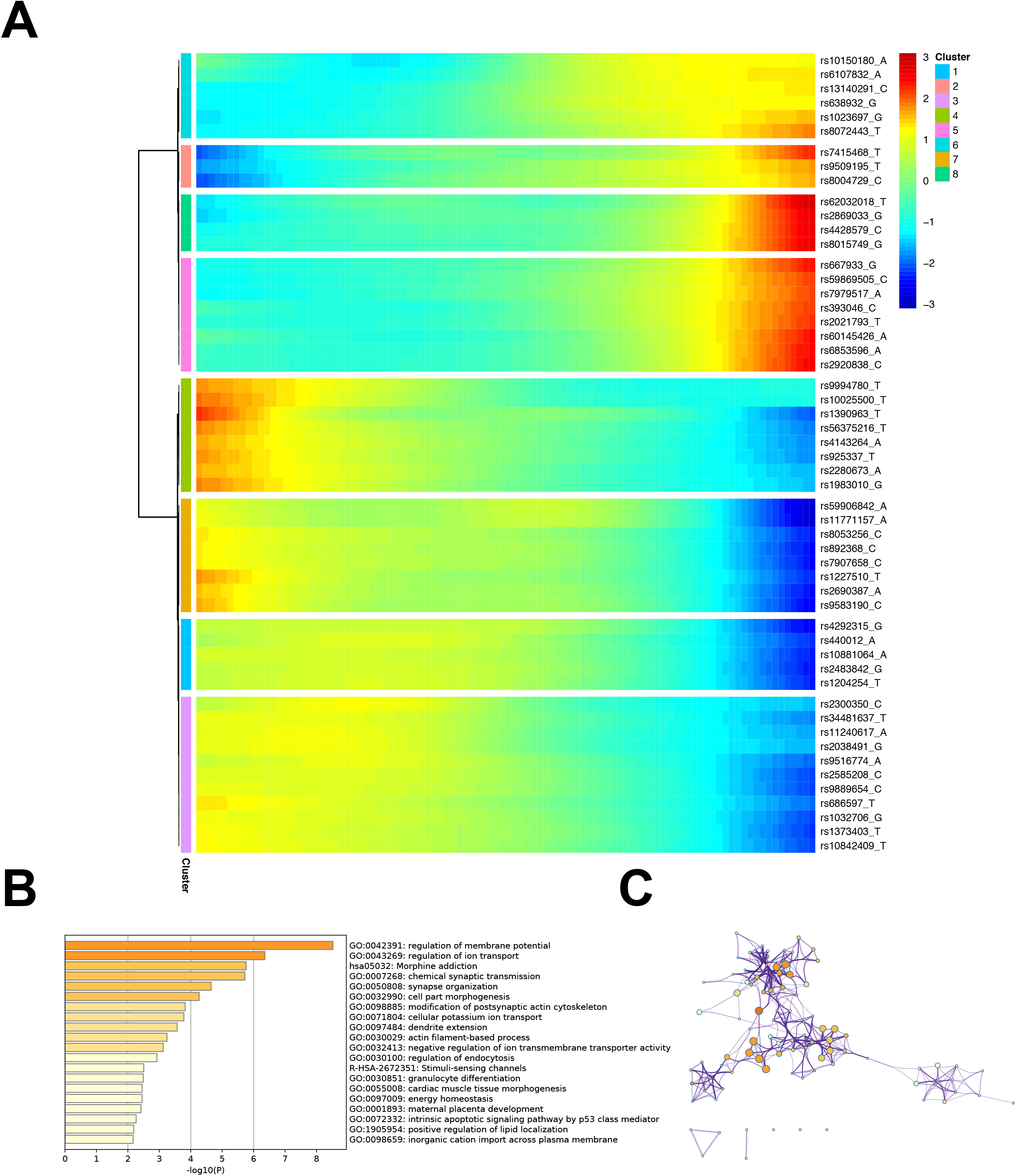
A. Heatmap of the first 50 differential SNPs following Human trajectory trends along 1 to 6 branches identified by Monocle. Color of heatmap indicated the normalized SNPs genotype index. B. GO signal pathway enrichment analysis of the first 300 differential SNPs in Matescope. C. Signal pathways correlation analysis, connect thinness indicated the p-value.

## Resources and Methods

### Date sets

Collated Human 1000 Genomes Project Phase III data were from website https://lovingscience.com/ancestries/human/. The recoded_1000G.raw file was used (Gaspar and Breen, 2019). E11 Data is from R package radmixture of wegene company. K12b data is a part of Dodecad project http://dodecad.blogspot.com/. Data of two calculators E11 and K12b are from the community, and there is no clear author. Real author of those please contact the corresponding author on Twitter after seeing this paper.

### Softwares

R, RStudio were downloaded from the official websites. Important R packages used in this paper include Monocle (Qiu et al., 2017a; Qiu et al., 2017b; Trapnell et al., 2014), radmixture, https://CRAN.R-project.org/package=radmixture, dbscan, Rtsne (Maaten, 2014; van der Maaten and Hinton, 2009), DDRTree (Mao et al., 2015a).

### Human Trajectories

We used the R Monocle package to perform trajectory analysis of population genetic data. The Monocle package mainly learns the gradual process of population genetic data and uses the built-in DDRTree algorithm to reduce the population genetic data and actively return to the tree to display the trajectory. Monocle’s construction of the population trajectory is mainly through two steps. The first step is to filter out the key SNP loci. This step mainly screens out a group of SNP loci with a large coefficient of variation under a certain expression level. In our analysis, we screened 276 SNP loci. The second step is to use the obtained SNP loci for dimensionality reduction and trajectory construction, which utilizes the DDRTree algorithm. Using the differentGeneTest function in the Monocle package, we calculated the SNP sites with the largest difference between states and plotted the smoothed heat map.

### TSNE + DDRTree

The tSNE dimension reduction is performed on the dataset. The idea here is that tSNE retains more details of the original data while reducing the dimension. The original data shows a linear change. After the tSNE reduces the dimension, there is still a certain relationships are preserved in the results. The advantage of using all of the SNPs sites is that more information is retained. The code is publicly available on github https://github.com/weizhuang128/TSNE_DDRTree_Human_Trajectories.

After the tSNE dimension reduction, the two-dimensional result is learned again by the DDRTree algorithm, and finally the Z component of DDRTree is taken out as a graph, which will greatly improve the detail retention of the cluster and relationship. This method can also greatly save computing resources and increase the speed. At the same time, we tested the initial dimensionality reduction by using UMAP or PCA, but did not get the ideal result, which shows the usability of tSNE in this method, Fig S3 to 5. But it does not rule out that UMAP, a manifold-based analysis method, has a higher degree of reduction in the real world.

### E11, K12B and K13 analysis

E11 is a calculator popular in the community of individual genetic testing enthusiasts, and it is rumored in the community that it has a very high resolving power for people in East Asia. E11 has its implementation in the R package radmixture. K12B and K13 are another implementation of the ancestor calculator in the R package radmixture. The calculation results of E11, K12B and K13 are visualized and displayed in the plot after being normalized.

### Signal pathway enrichment analysis

The different SNP in the different states will be found using the differentGeneTest function in Monocle, and the gene in which the SNP locus is located will be found and entered into the http://metascape.org/ website to perform the bioprocess enrichment analysis (Zhou et al., 2019).

## Reference

Adachi, N., Kakuda, T., Takahashi, R., Kanzawa-Kiriyama, H., and Shinoda, K.-I. (2018). Ethnic derivation of the Ainu inferred from ancient mitochondrial DNA data. Am J Phys Anthropol 165, 139-148.

Cavalli-Sforza, L.L., and Feldman, M.W. (2003). The application of molecular genetic approaches to the study of human evolution. Nature Genetics 33, 266-275.

Crema, E.R., Habu, J., Kobayashi, K., and Madella, M. (2016). Summed Probability Distribution of 14C Dates Suggests Regional Divergences in the Population Dynamics of the Jomon Period in Eastern Japan. PLOS ONE 11, e0154809.

Diaz-Papkovich, A., Anderson-Trocmé, L., Ben-Eghan, C., and Gravel, S. (2019). UMAP reveals cryptic population structure and phenotype heterogeneity in large genomic cohorts. PLoS Genet 15, e1008432–e1008432.

Garrigan, D., and Hammer, M.F. (2006). Reconstructing human origins in the genomic era. Nature Reviews Genetics 7, 669-680.

Gaspar, H.A., and Breen, G. (2019). Probabilistic ancestry maps: a method to assess and visualize population substructures in genetics. BMC Bioinformatics 20, 116-116.

Hanihara, T., Matsumura, H., Kawakubo, Y., Nguyen, L.C., Nguyen, K.T., Oxenham, M.F., and Dodo, Y. (2012). Population history of northern Vietnamese inferred from nonmetric cranial trait variation. Anthropological Science advpub, 1202070129-1202070129.

He, J.-D., Peng, M.-S., Quang, H.H., Dang, K.P., Trieu, A.V., Wu, S.-F., Jin, J.-Q., Murphy, R.W., Yao, Y.-G., and Zhang, Y.-P. (2012). Patrilineal Perspective on the Austronesian Diffusion in Mainland Southeast Asia. PLOS ONE 7, e36437.

Iizuka, M., and Nakahashi, T. (2002). A population genetic study on the transition from Jomon people to Yayoi people. Genes & genetic systems 77, 287-300.

Janhunen, J. (2005). Tungusic: an endangered language family in Northeast Asia. In International Journal of the Sociology of Language, pp. 37.

Jinam, T., Nishida, N., Hirai, M., Kawamura, S., Oota, H., Umetsu, K., Kimura, R., Ohashi, J., Tajima, A., Yamamoto, T., et al. (2012). The history of human populations in the Japanese Archipelago inferred from genome-wide SNP data with a special reference to the Ainu and the Ryukyuan populations. Journal of Human Genetics 57, 787-795.

Kim, W., Shin, D.J., Harihara, S., and Kim, Y.J. (2000). Y chromosomal DNA variation in East Asian populations and its potential for inferring the peopling of Korea. Journal of Human Genetics 45, 76-83.

Le, V.S., Tran, K.T., Bui, H.T.P., Le, H.T.T., Nguyen, C.D., Do, D.H., Ly, H.T.T., Pham, L.T.D., Dao, L.T.M., and Nguyen, L.T. (2019). A Vietnamese human genetic variation database. Human Mutation 40, 1664-1675.

Lee, S., and Hasegawa, T. (2011). Bayesian phylogenetic analysis supports an agricultural origin of Japonic languages. Proceedings of the Royal Society B: Biological Sciences 278, 3662-3669.

Lorenzo, A.D., Medvet, E., Tu, T., #353, ar, and Bartoli, A. (2019). An analysis of dimensionality reduction techniques for visualizing evolution. In Proceedings of the Genetic and Evolutionary Computation Conference Companion (Prague, Czech Republic: ACM), pp. 1864-1872.

Maaten, L.V.D. (2014). Accelerating t-SNE using tree-based algorithms. J Mach Learn Res 15, 3221-3245.

Mao, Q., Wang, L., Goodison, S., Sun, Y., st, A.C.M.S.C.o.K.D., and Data Mining, K.D.D. (2015a). Dimensionality reduction via graph structure learning. Proc ACM SIGKDD Int Conf Knowl Discov Data Min Proceedings of the ACM SIGKDD International Conference on Knowledge Discovery and Data Mining 2015-August, 765-774.

Mao, Q., Yang, L., Wang, L., Goodison, S., and Sun, Y. (2015b). SimplePPT: A Simple Principal Tree Algorithm. In, pp. 792-800.

Nanta, A. (2008). Physical Anthropology and the Reconstruction of Japanese Identity in Postcolonial Japan. Social Science Japan Journal 11, 29-47.

Qi, M., Li, W., Tsang, I.W., and Yijun, S. (2017). Principal Graph and Structure Learning Based on Reversed Graph Embedding. IEEE Trans Pattern Anal Mach Intell 39, 2227-2241.

Qiu, X., Hill, A., Packer, J., Lin, D., Ma, Y.-A., and Trapnell, C. (2017a). Single-cell mRNA quantification and differential analysis with Census. Nat Methods 14, 309-315.

Qiu, X., Mao, Q., Tang, Y., Wang, L., Chawla, R., Pliner, H.A., and Trapnell, C. (2017b). Reversed graph embedding resolves complex single-cell trajectories. Nat Methods 14, 979-982.

Shendure, J. (2011). Next-generation human genetics. Genome Biol 12, 408-408.

Su, B., Xiao, J., Underhill, P., Deka, R., Zhang, W., Akey, J., Huang, W., Shen, D., Lu, D., Luo, J., et al. (1999). Y-Chromosome evidence for a northward migration of modern humans into Eastern Asia during the last Ice Age. Am J Hum Genet 65, 1718-1724.

Sun, H., Zhou, C., Huang, X., Lin, K., Shi, L., Yu, L., Liu, S., Chu, J., and Yang, Z. (2013). Autosomal STRs provide genetic evidence for the hypothesis that Tai people originate from southern China. PloS one 8, e60822–e60822.

Tian, C., Gregersen, P.K., and Seldin, M.F. (2008). Accounting for ancestry: population substructure and genome-wide association studies. Hum Mol Genet 17, R143–R150.

Tishkoff, S.A., and Verrelli, B.C. (2003). Patterns of Human Genetic Diversity: Implications for Human Evolutionary History and Disease. Annual Review of Genomics and Human Genetics 4, 293-340.

Trapnell, C., Cacchiarelli, D., Grimsby, J., Pokharel, P., Li, S., Morse, M., Lennon, N.J., Livak, K.J., Mikkelsen, T.S., and Rinn, J.L. (2014). The dynamics and regulators of cell fate decisions are revealed by pseudotemporal ordering of single cells. Nature Biotechnology 32, 381-386.

van der Maaten, L., and Hinton, G. (2009). Visualizing Data using t-SNE. Journal of machine learning research : JMLR 9, 2579-2606.

Watanobe, T., Ishiguro, N., Nakano, M., Matsui, A., Hongo, H., Yamazaki, K., and Takahashi, O. (2004). Prehistoric Sado Island Populations of Sus scrofa Distinguished from Contemporary Japanese Wild Boar by Ancient Mitochondrial DNA. Zoological Science 21, 219-228, 210.

Xia, Z.-Y., Yan, S., Wang, C.-C., Zheng, H.-X., Zhang, F., Liu, Y.-C., Yu, G., Yu, B.-X., Shu, L.-L., and Jin, L. (2019). Inland-coastal bifurcation of southern East Asians revealed by Hmong-Mien genomic history. bioRxiv, 730903.

Zhang, J., Wang, X., Kruger, U., and Wang, F. (2011). Principal Curve Algorithms for Partitioning High-Dimensional Data Spaces. IEEE Transactions on Neural Networks 22, 367-380.

Zhou, Y., Zhou, B., Pache, L., Chang, M., Khodabakhshi, A.H., Tanaseichuk, O., Benner, C., and Chanda, S.K. (2019). Metascape provides a biologist-oriented resource for the analysis of systems-level datasets. Nature communications 10.

