## Supplementary Figure Legends for "Graph Embedding Method Based Genetical Trajectory Reveals Migration History Among East Asians"

**Fig S1** Reversed graph embedding method learned dimension reduction plot by using tSNE and DDRTree combined method of Genome 1000 program whole world data. Plots color indicated the areas.

**Fig S2** tSNE dimension reduction of whole world population of Genome 1000 program data. Each point indicated one individua, Plots color indicated the area groups from Genome 1000 program.

**Fig S3** The complex trajectory of East Asian population of Genome 1000 program data generated by Monocle. The level of the color indicates the score of E11.

**Fig S4** Umap one-step dimensionality reduction analysis for Genome 1000 program whole world data. The level of the color indicates the score of E11, K13 or K12B.

**Fig S5** Umap-DDRTree two-step dimensionality reduction analysis for Genome 1000 program whole world data. The level of the color indicates the score of E11, K13 or K12B.
