## Supplementary figures and images for "Graph Embedding Method Based Genetical Trajectory Reveals Migration History Among East Asians"

### Supplementary Figures 1-3

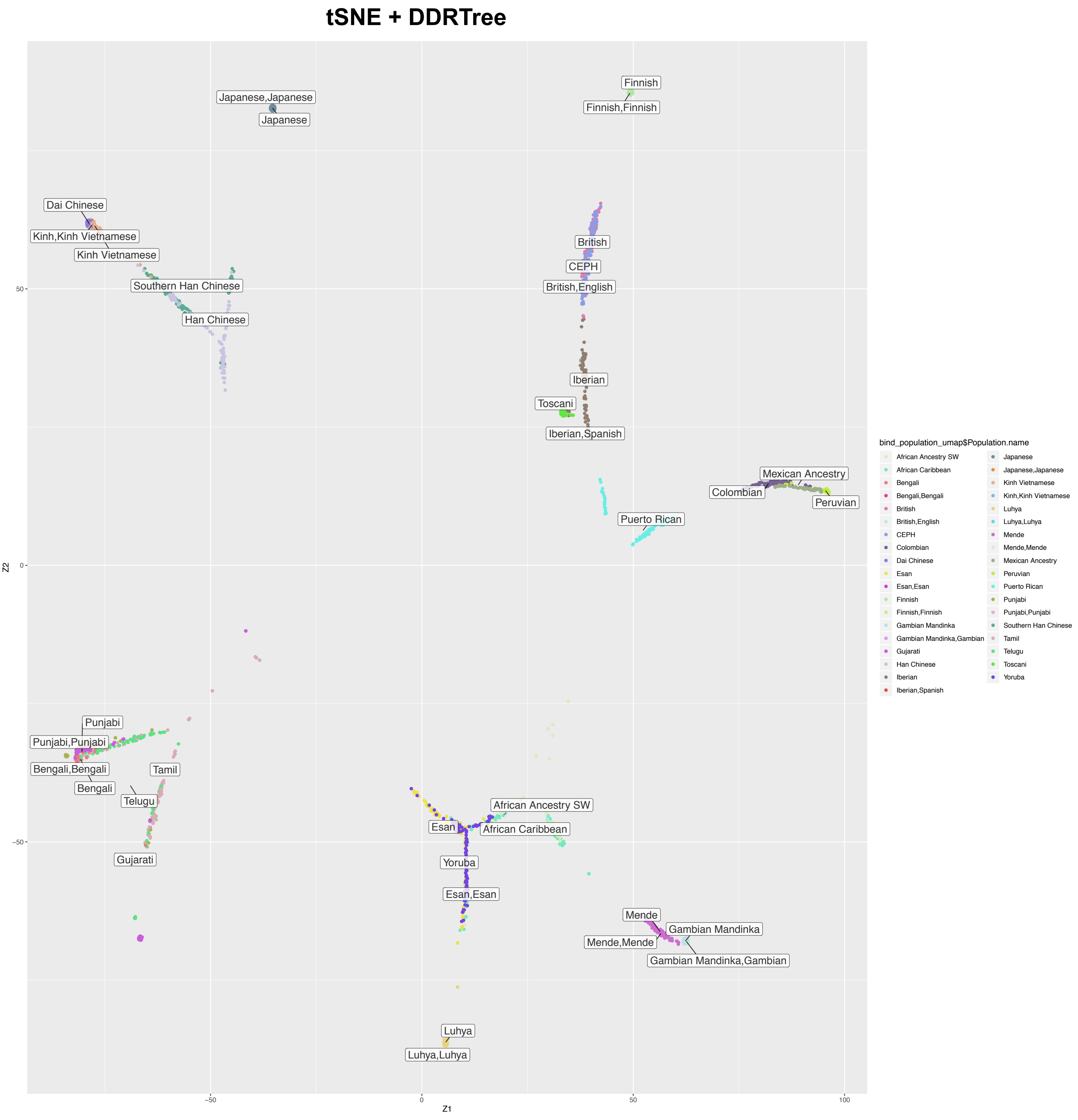

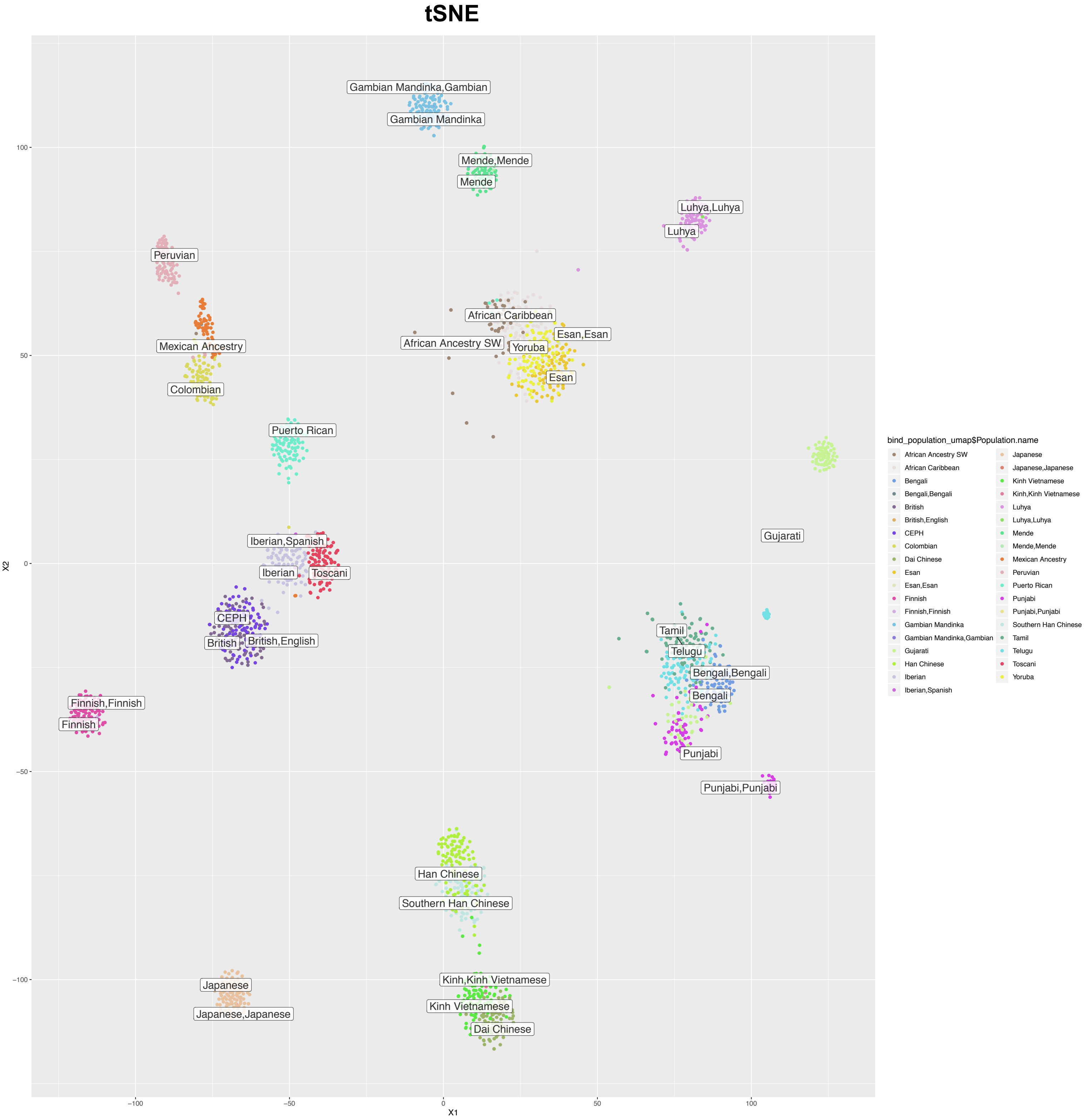

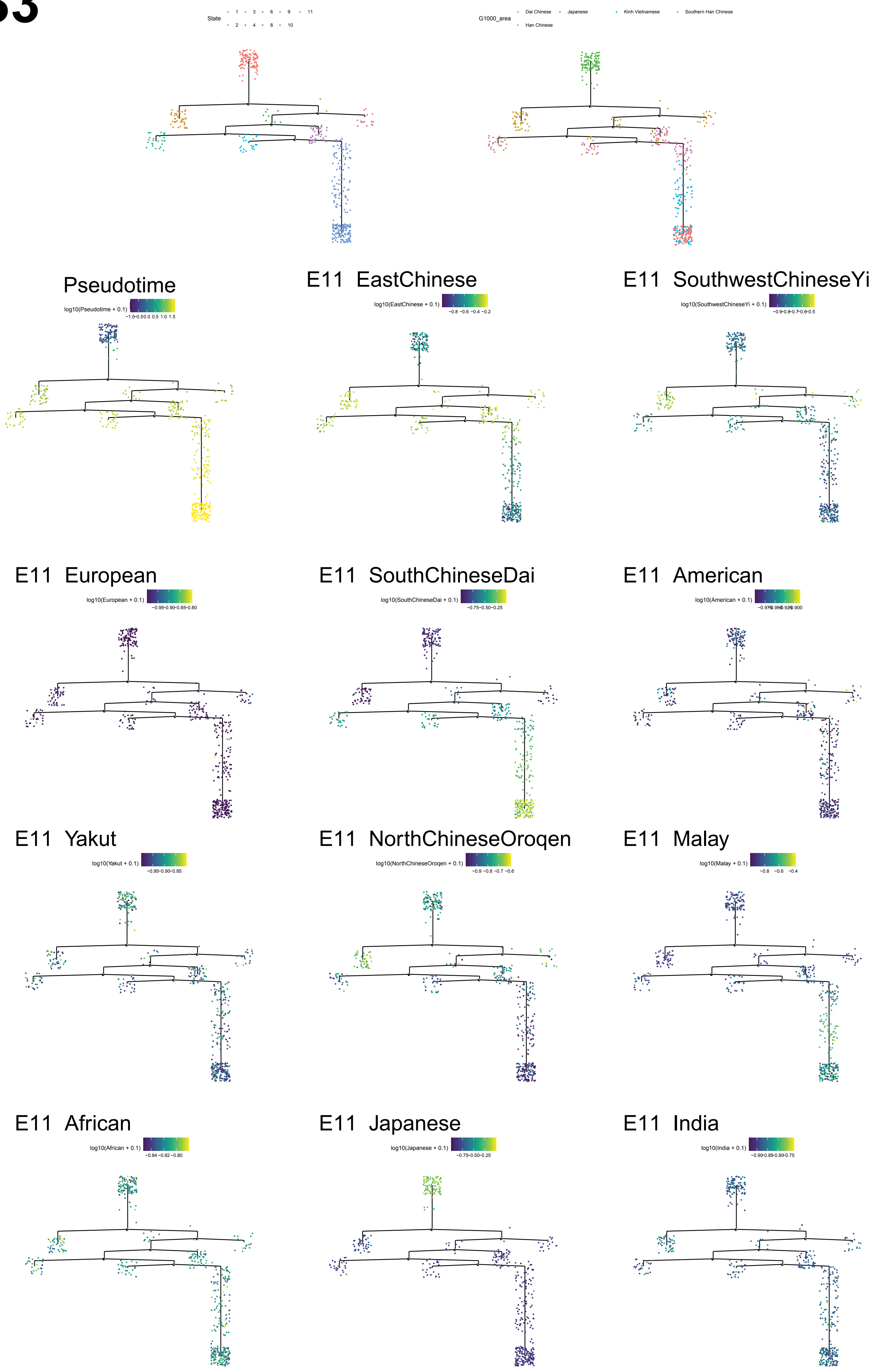
