## Supplementary Figures 4 for "Graph Embedding Method Based Genetical Trajectory Reveals Migration History Among East Asians"

### K13_Siberian

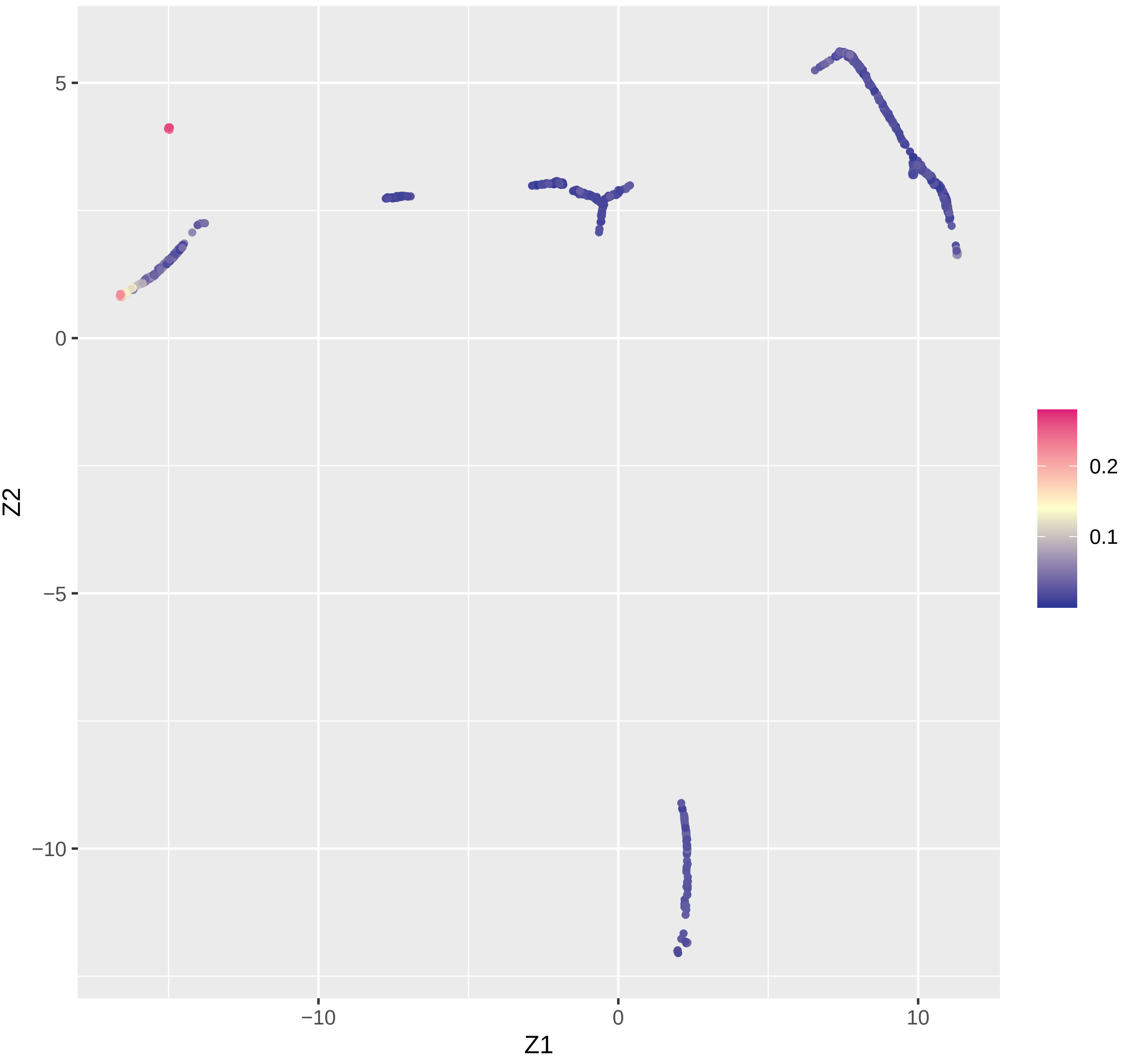

### K13_Amerindian

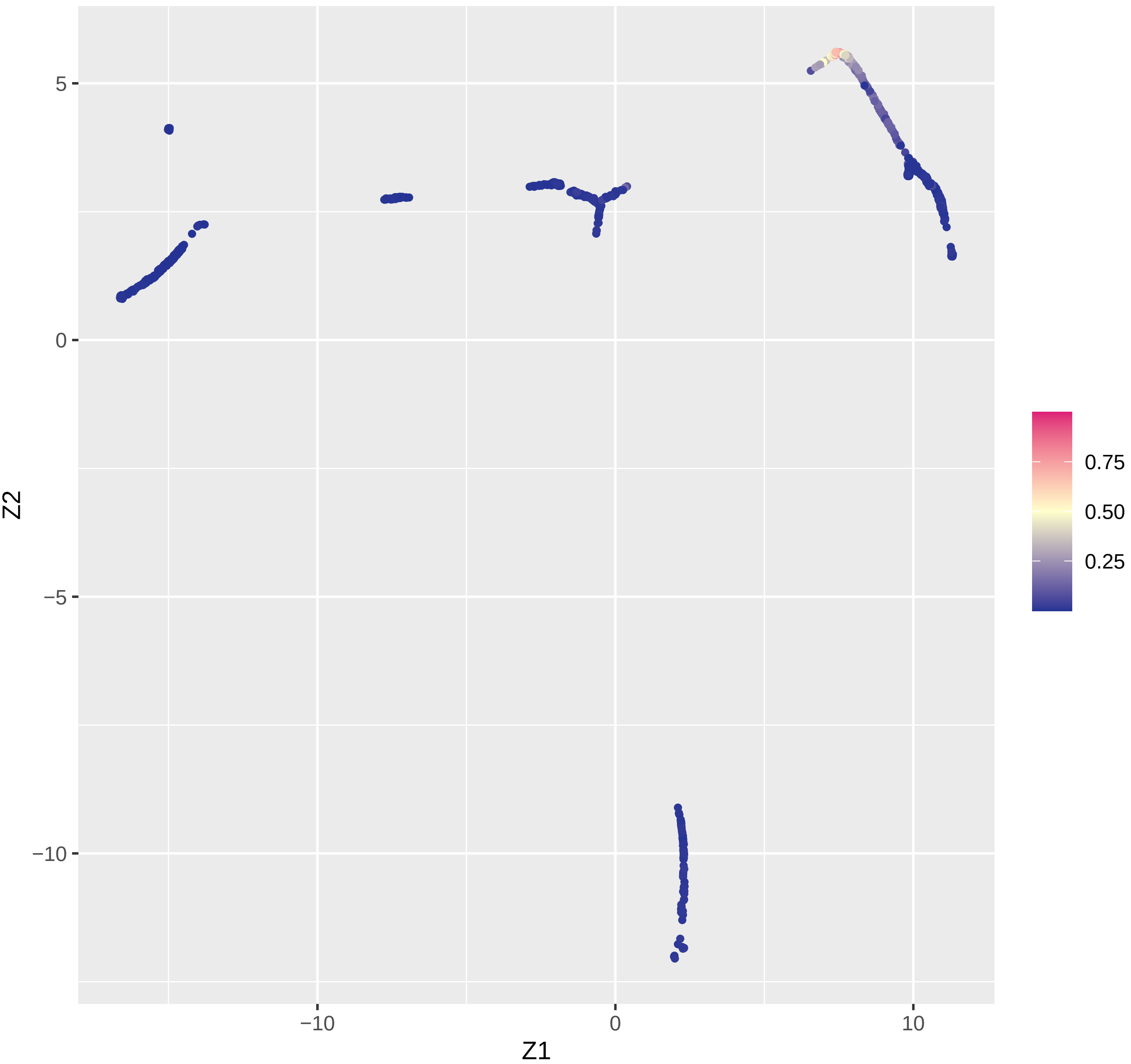

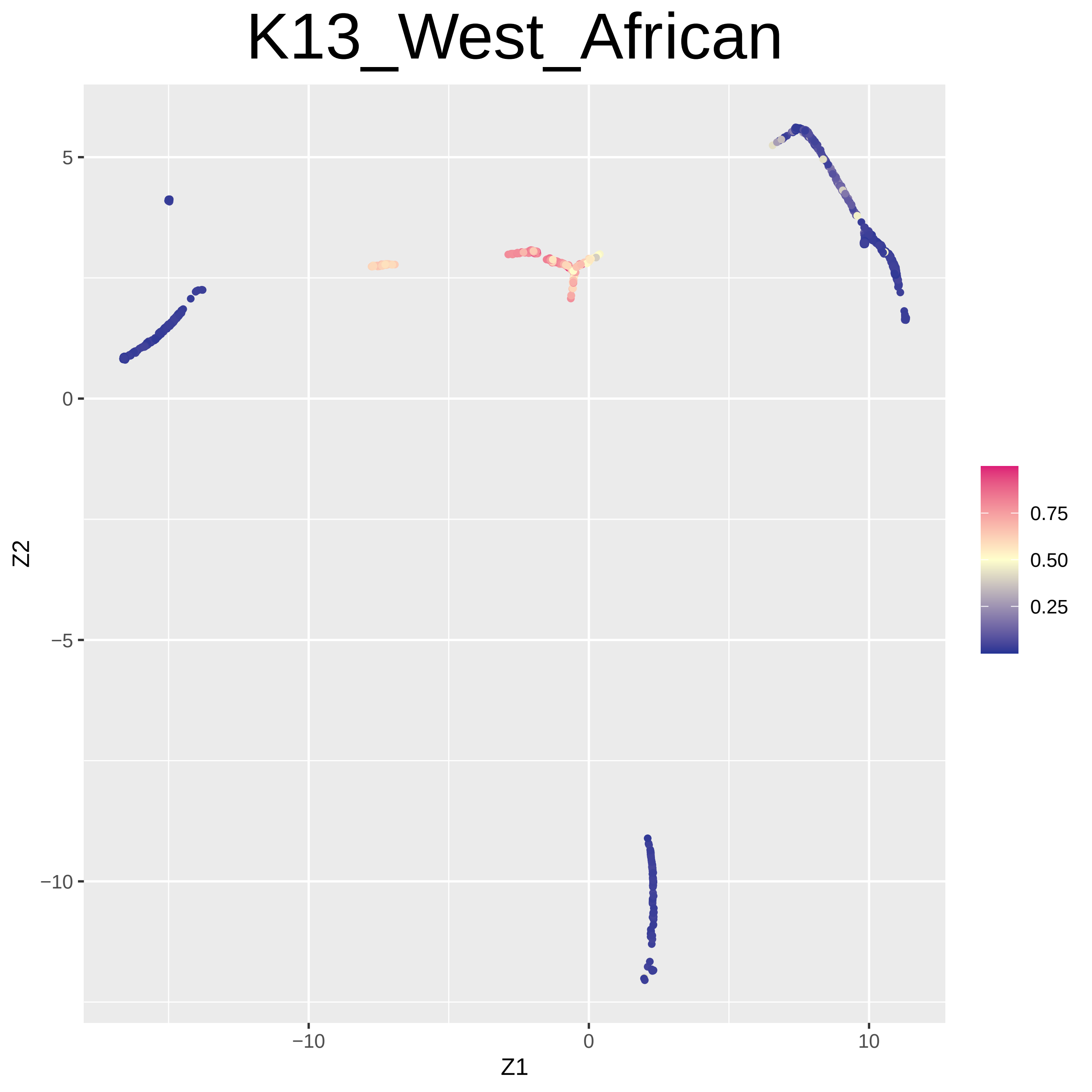

K13_West_African
5
0
−5
−10
−10
0
10
Z1
Z2
0.75
0.50
0.25

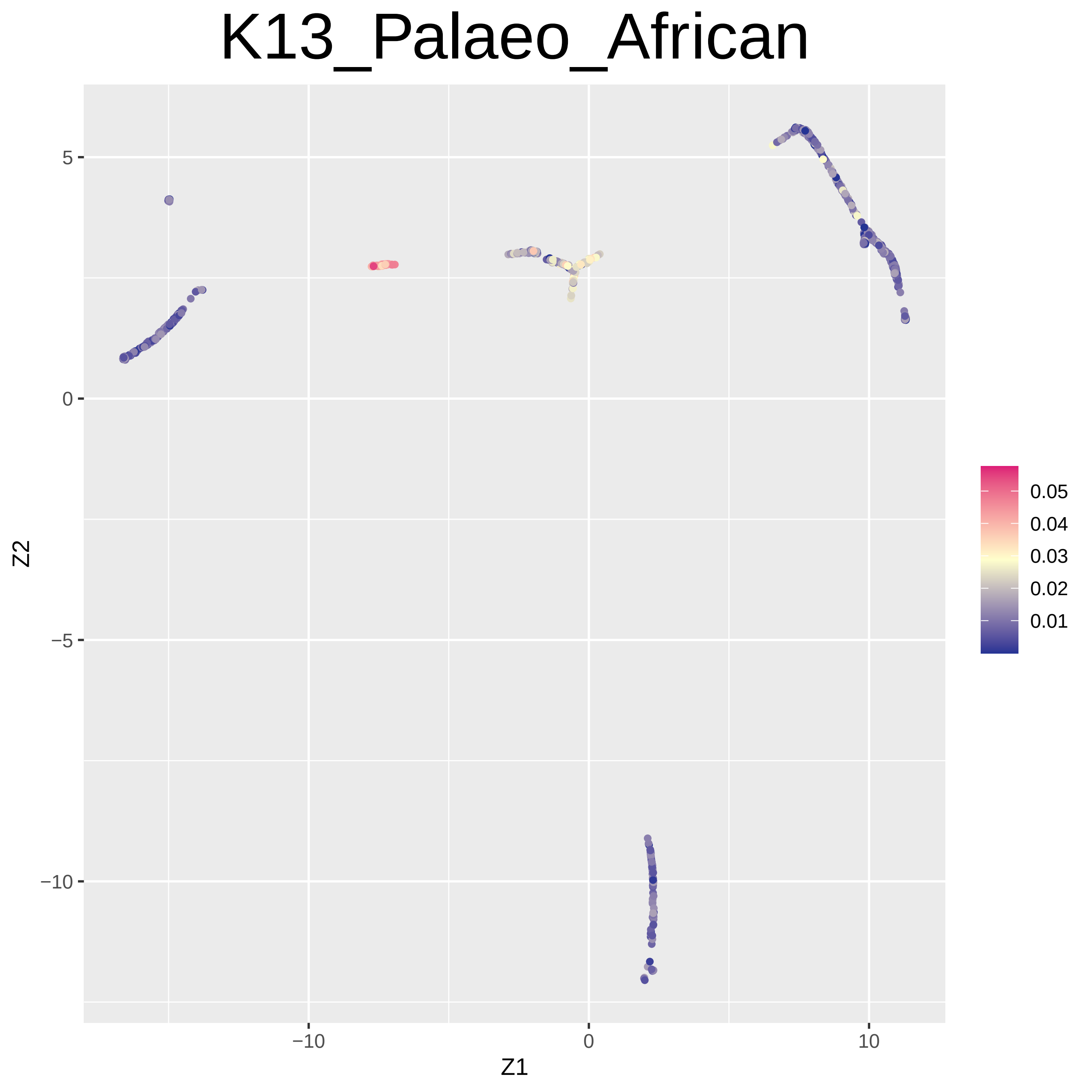

K13_Palaeo_African
Z2
Z1
5
0
−5
−10
−10
0
10
0.05
0.04
0.03
0.02
0.01

### K13_Southwest_Asian

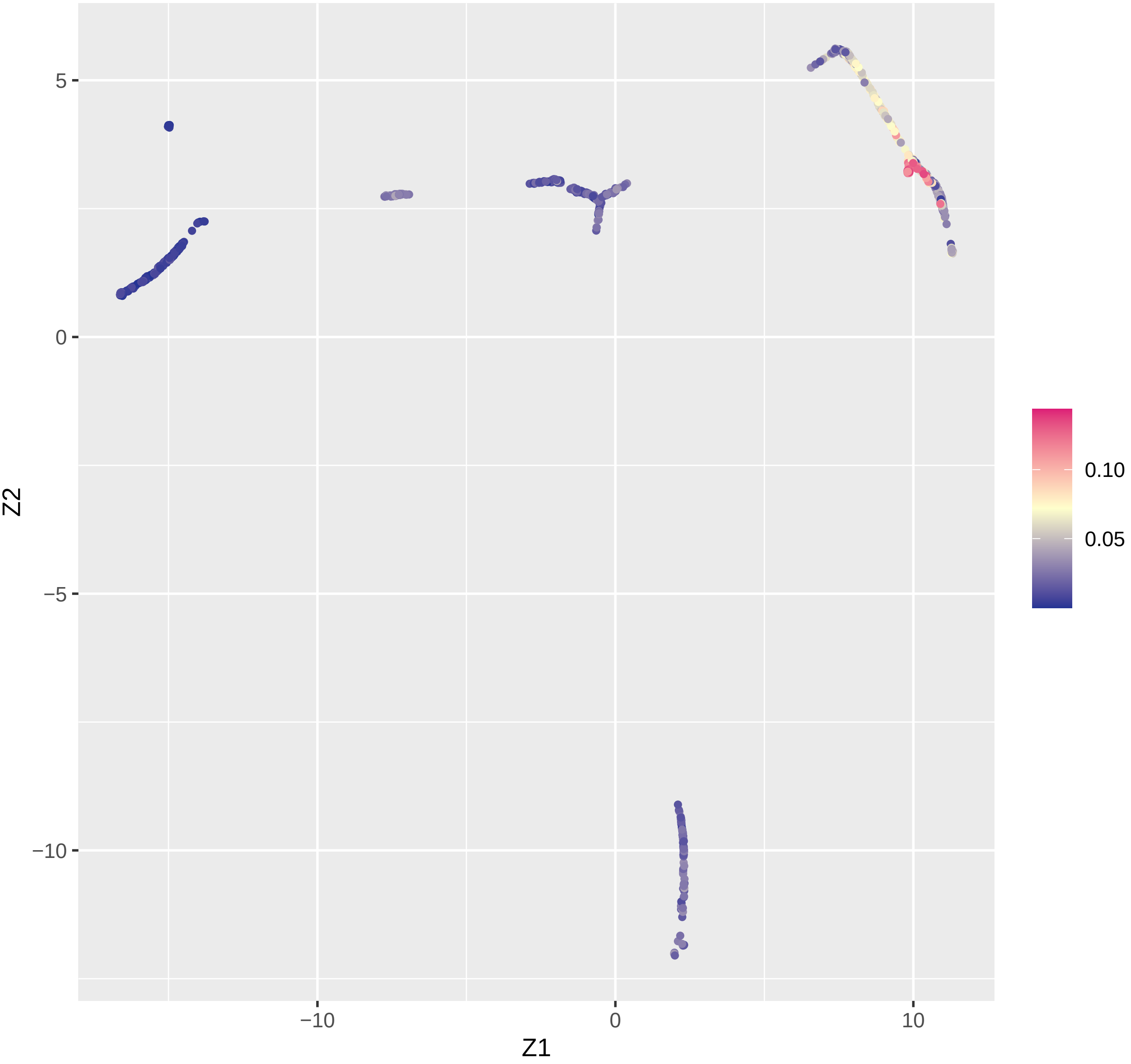

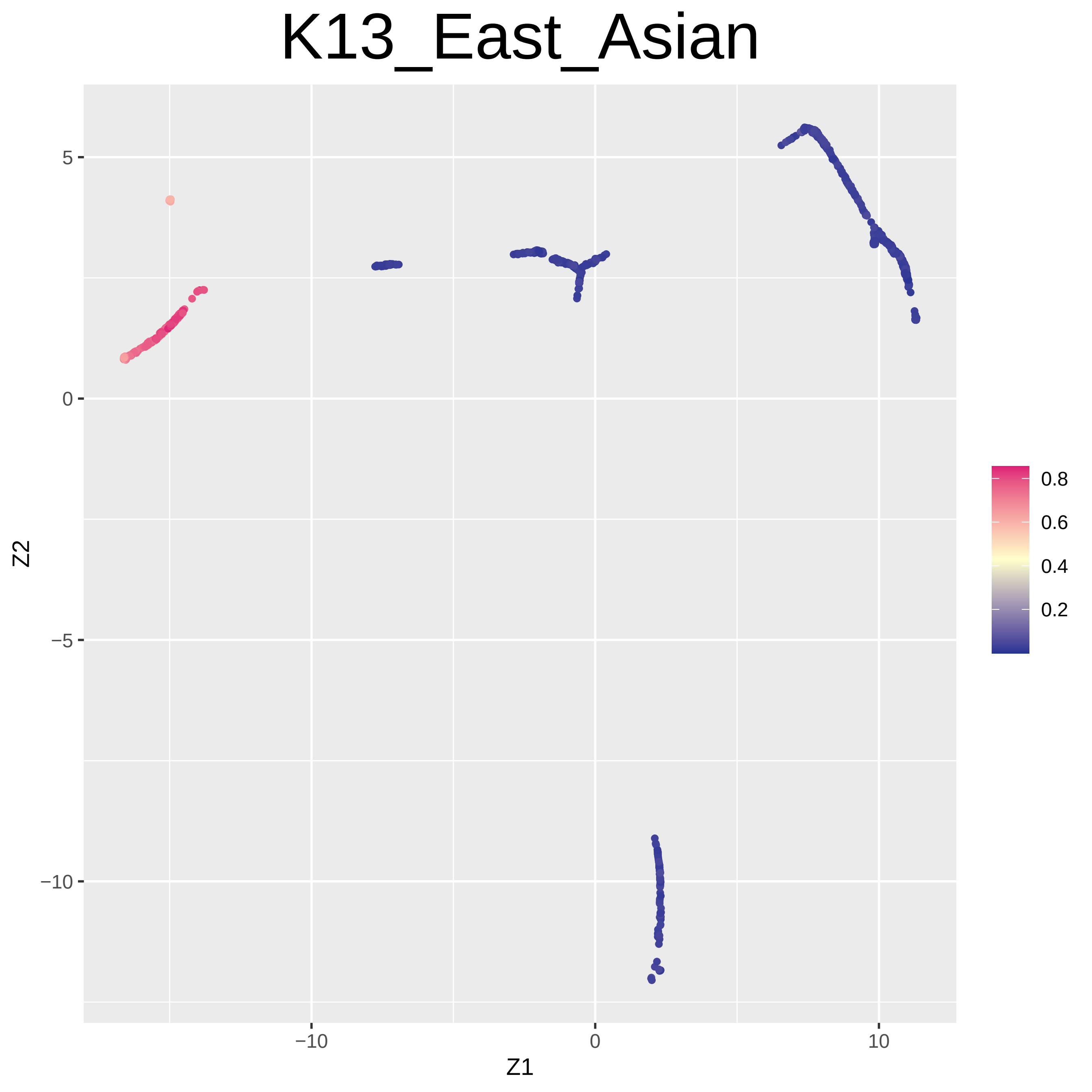

K13_East_Asian
5
0
−5
−10
Z2
−10
0
10
Z1
0.8
0.6
0.4
0.2

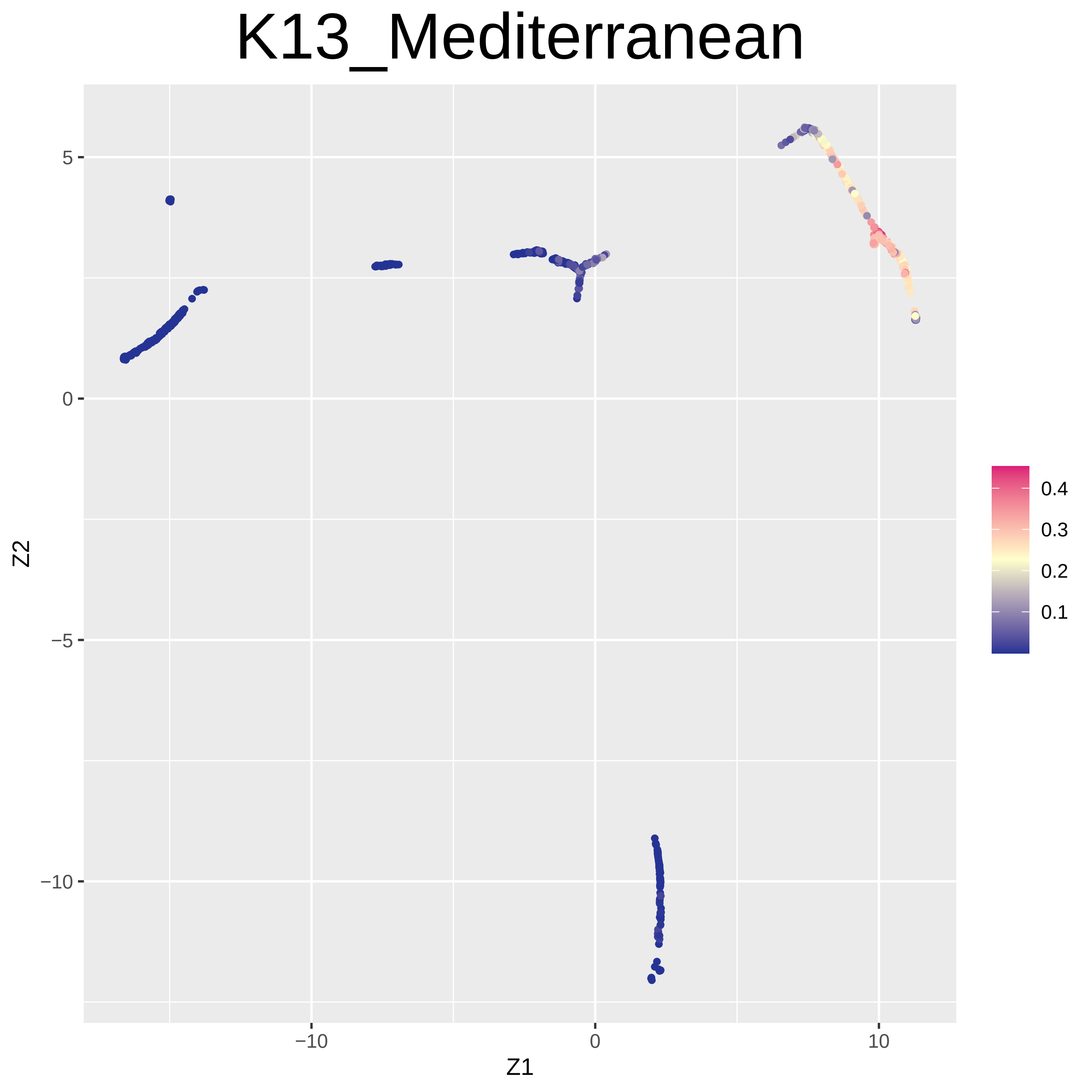

K13_Mediterranean
Z2
Z1
5
0
−5
−10
−10
0
10
0.4
0.3
0.2
0.1

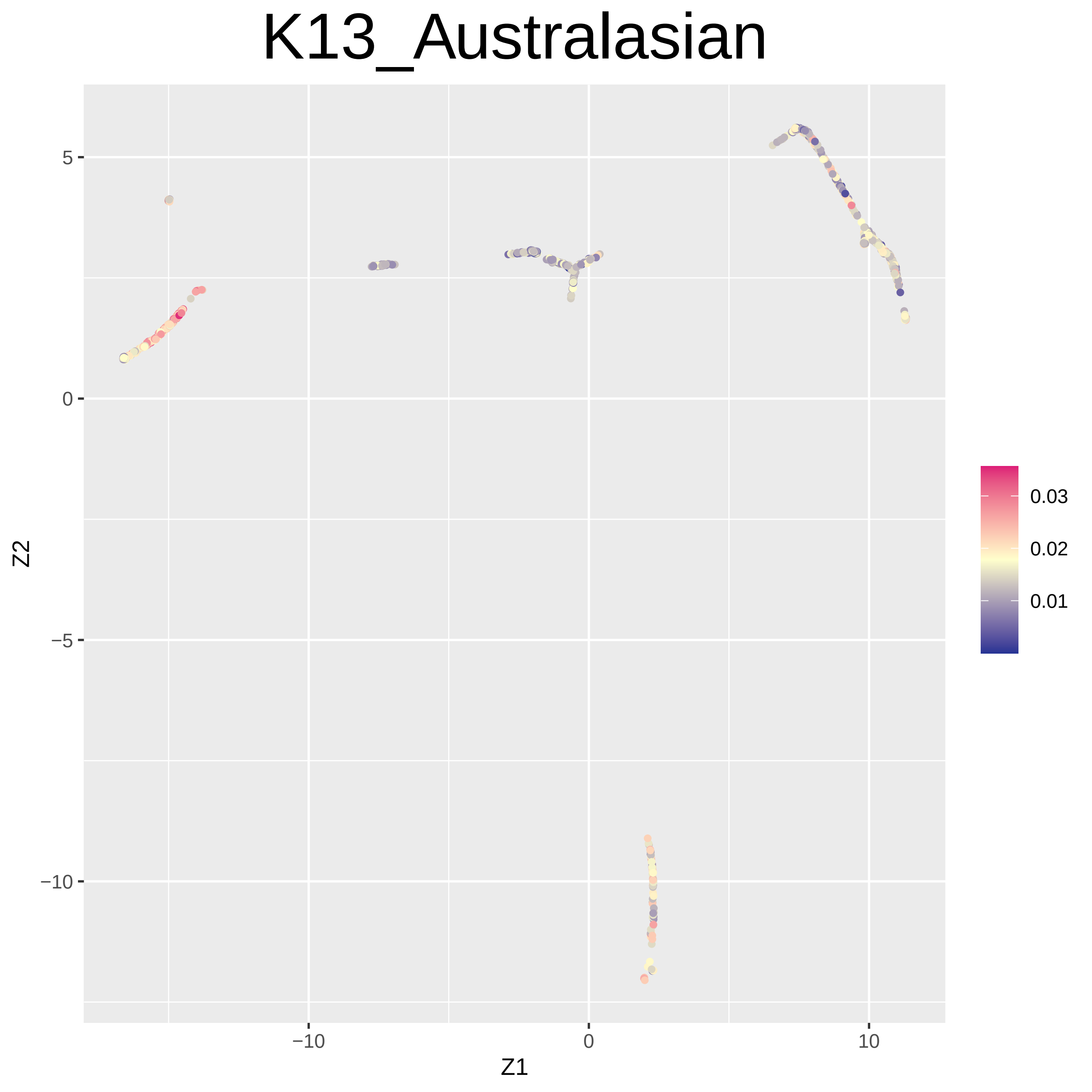

K13_Australasian
5
0
−5
−10
−10
0
10
Z1
Z2
0.03
0.02
0.01

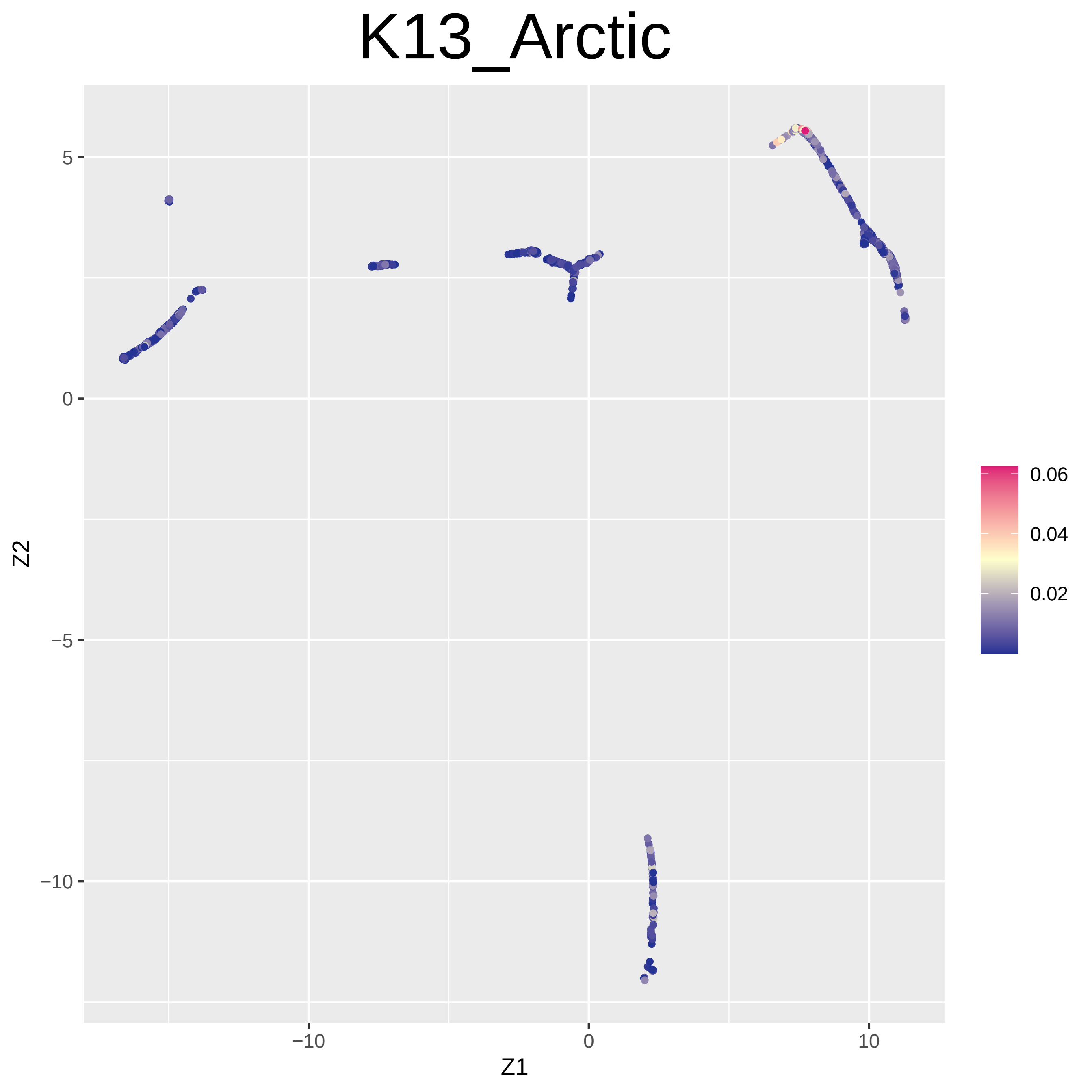

K13_Arctic
5
0
−5
−10
Z2
−10
0
10
Z1
0.06
0.04
0.02

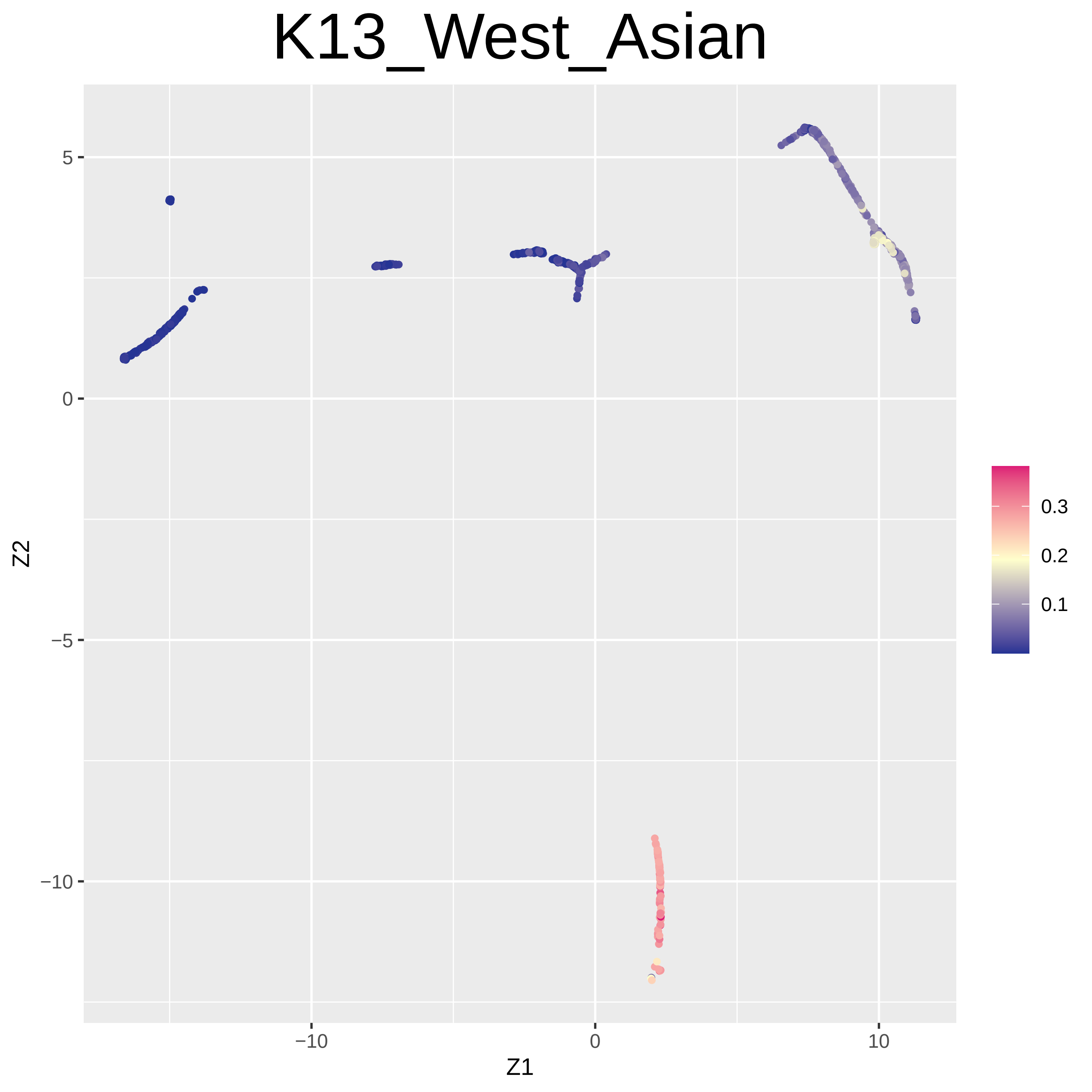
K13_West_Asian
Z2
Z1
5
0
−5
−10
−10
0
10
0.3
0.2
0.1

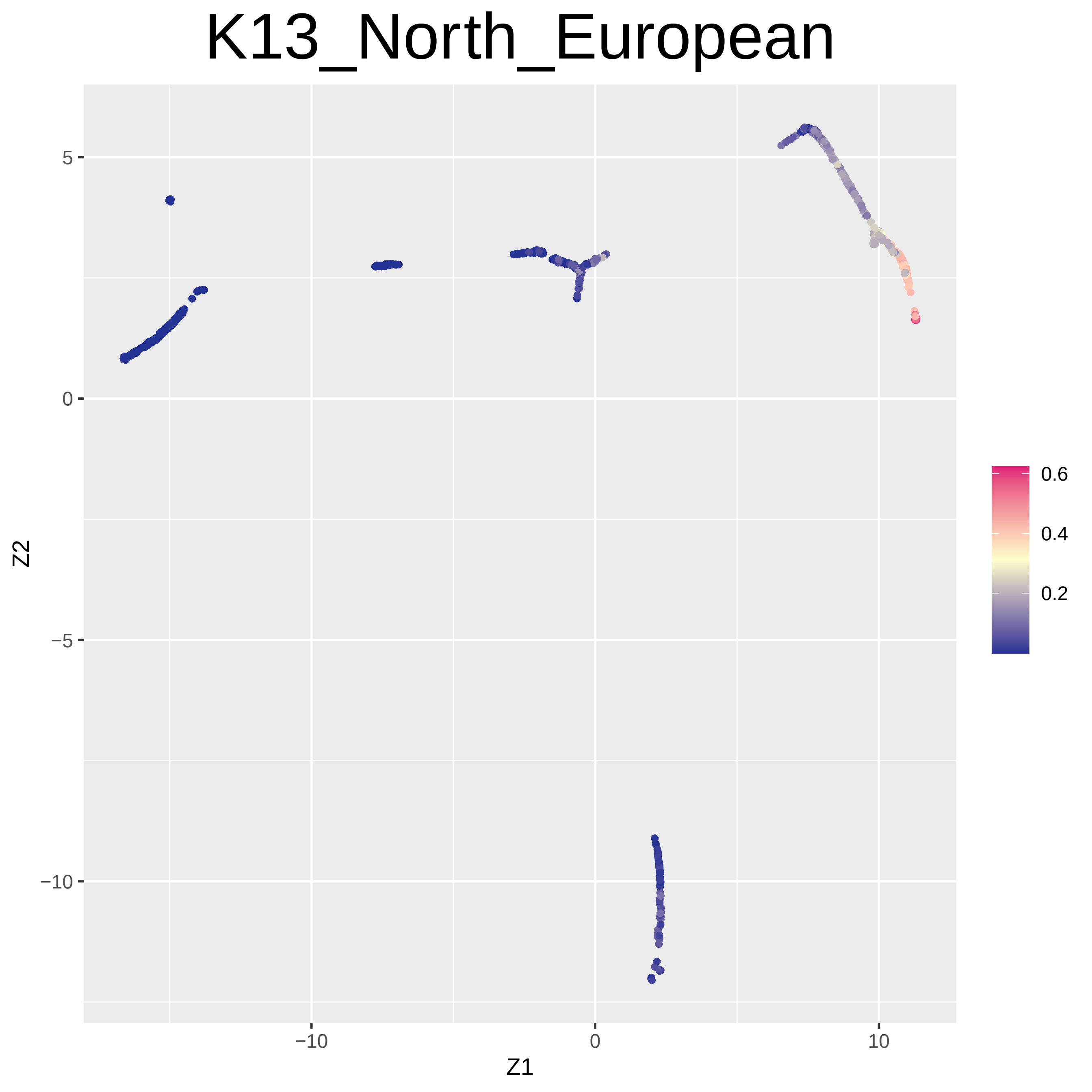

K13_North_European
5
0
−5
−10
Z2
−10
0
10
Z1
0.6
0.4
0.2

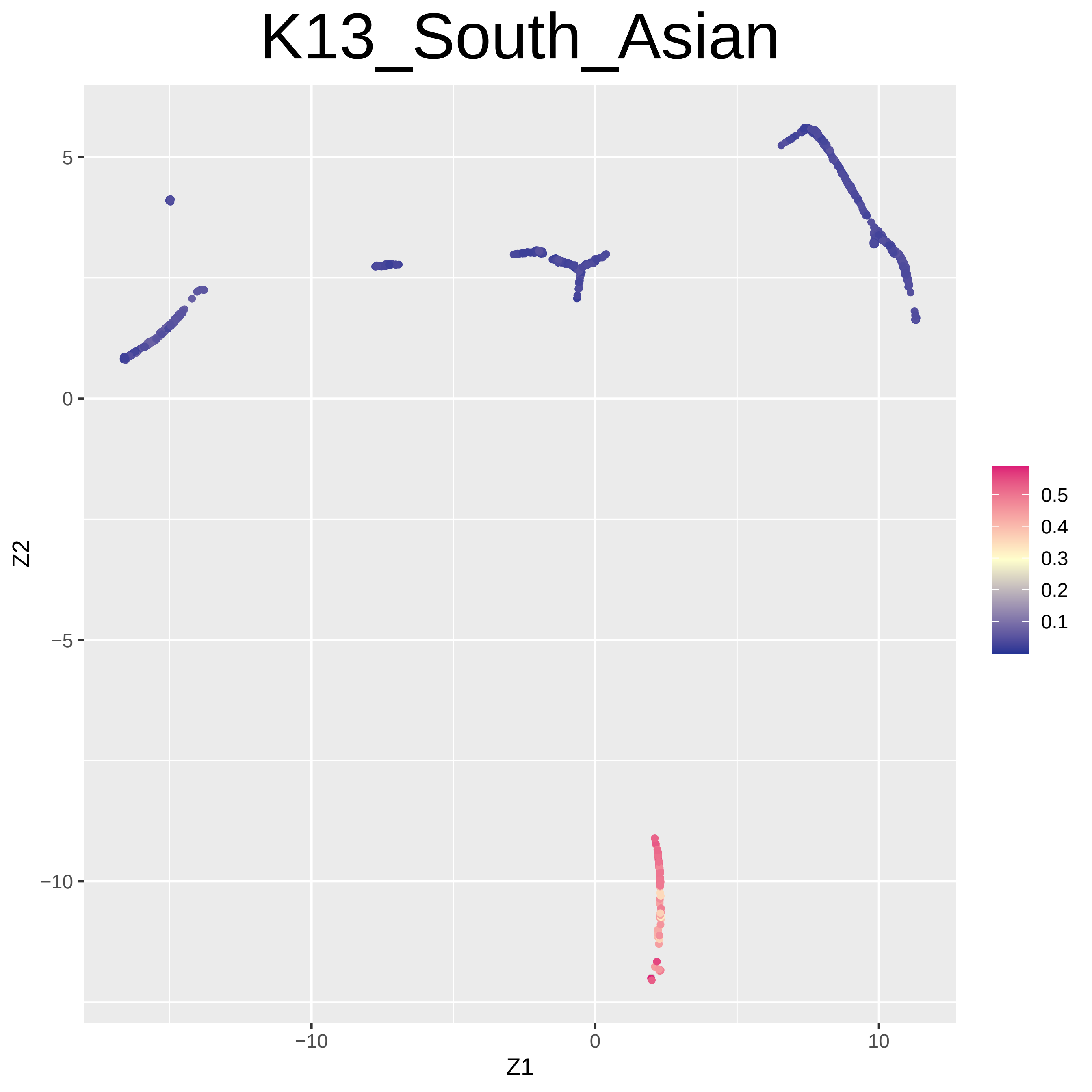

K13_South_Asian
5
0
−5
−10
−10
0
10
Z1
Z2
0.5
0.4
0.3
0.2
0.1

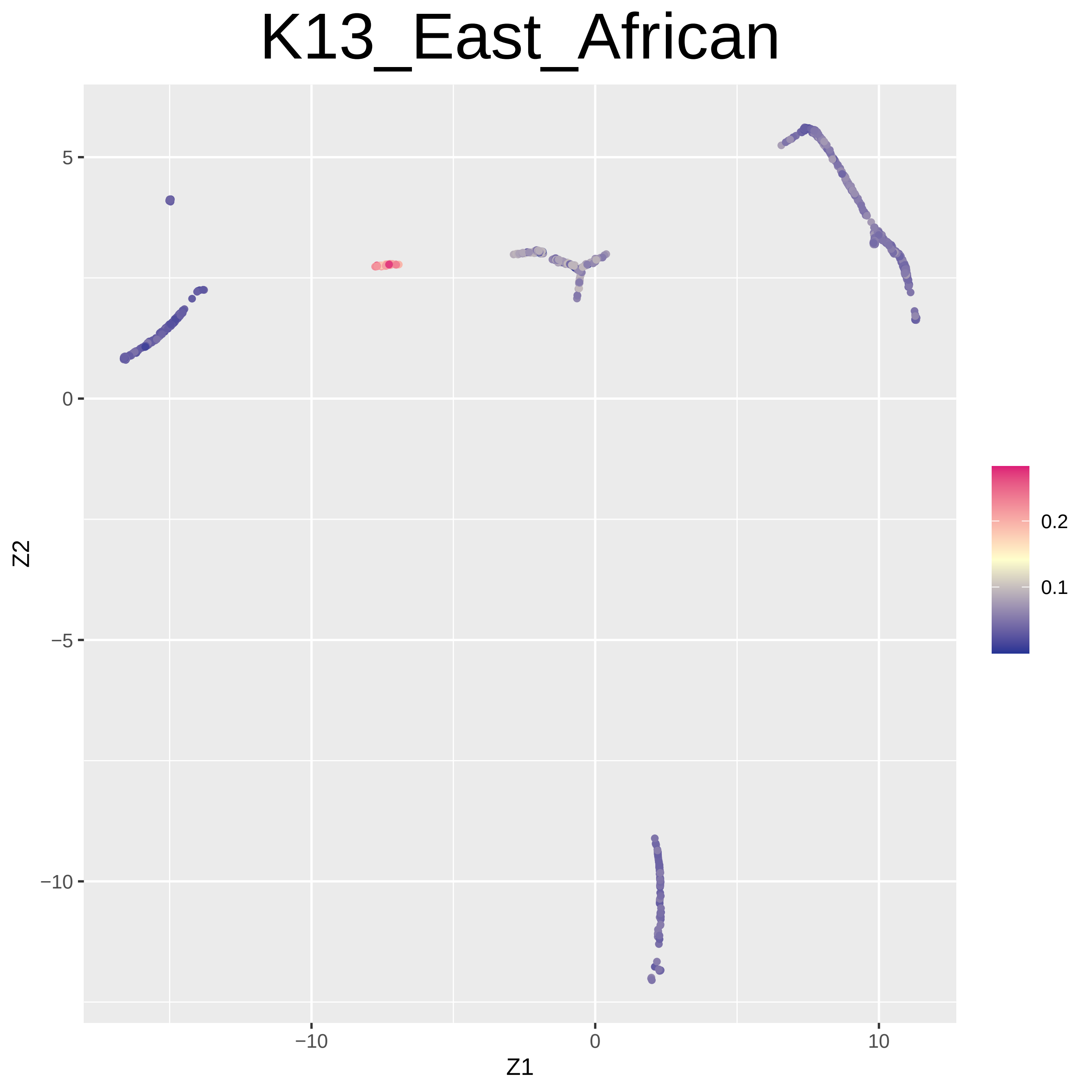

K13_East_African
5
0
−5
−10
−10
0
10
Z1
Z2
0.2
0.1

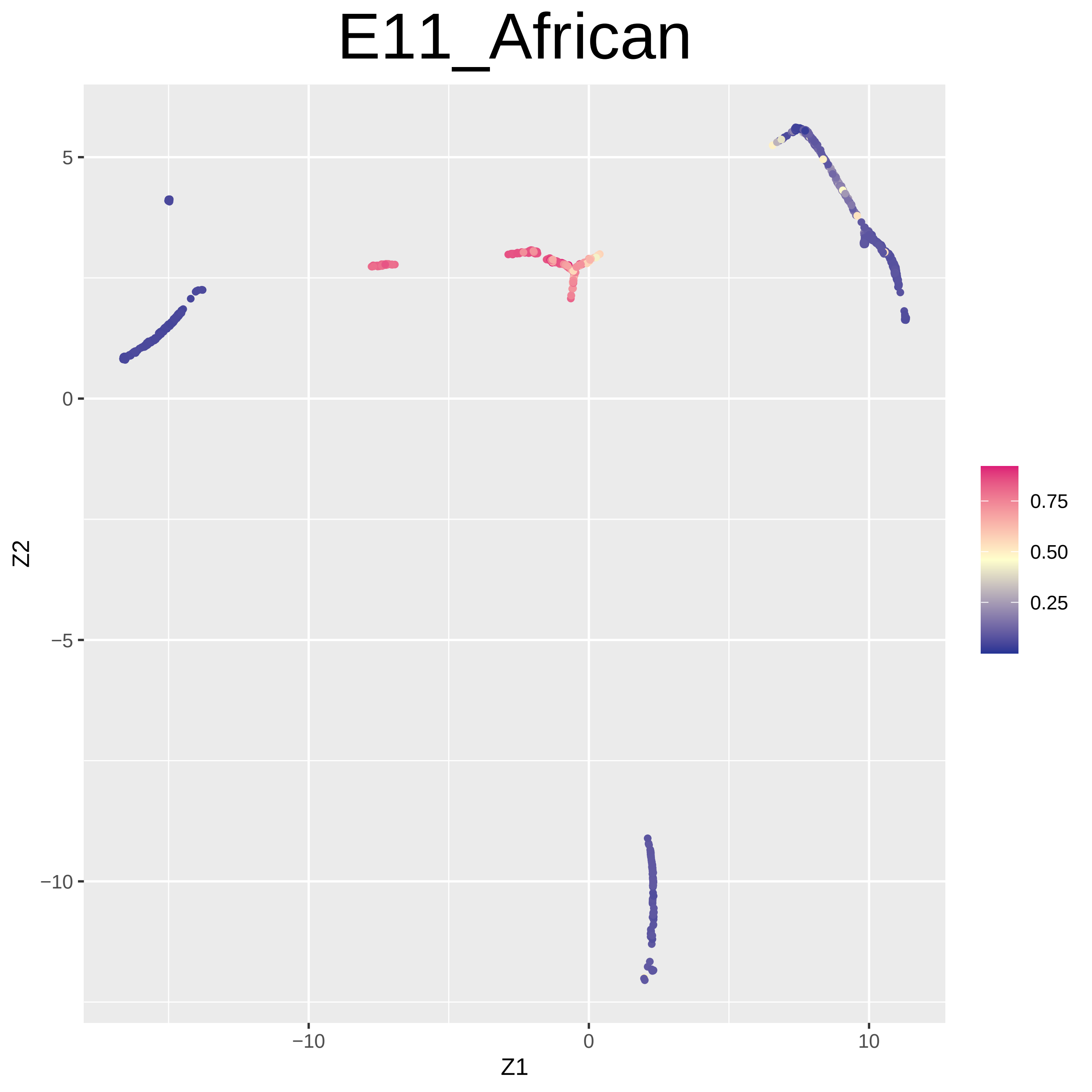

E11_African
Z2
Z1
5
0
−5
−10
−10
0
10
0.75
0.50
0.25

### E11_European

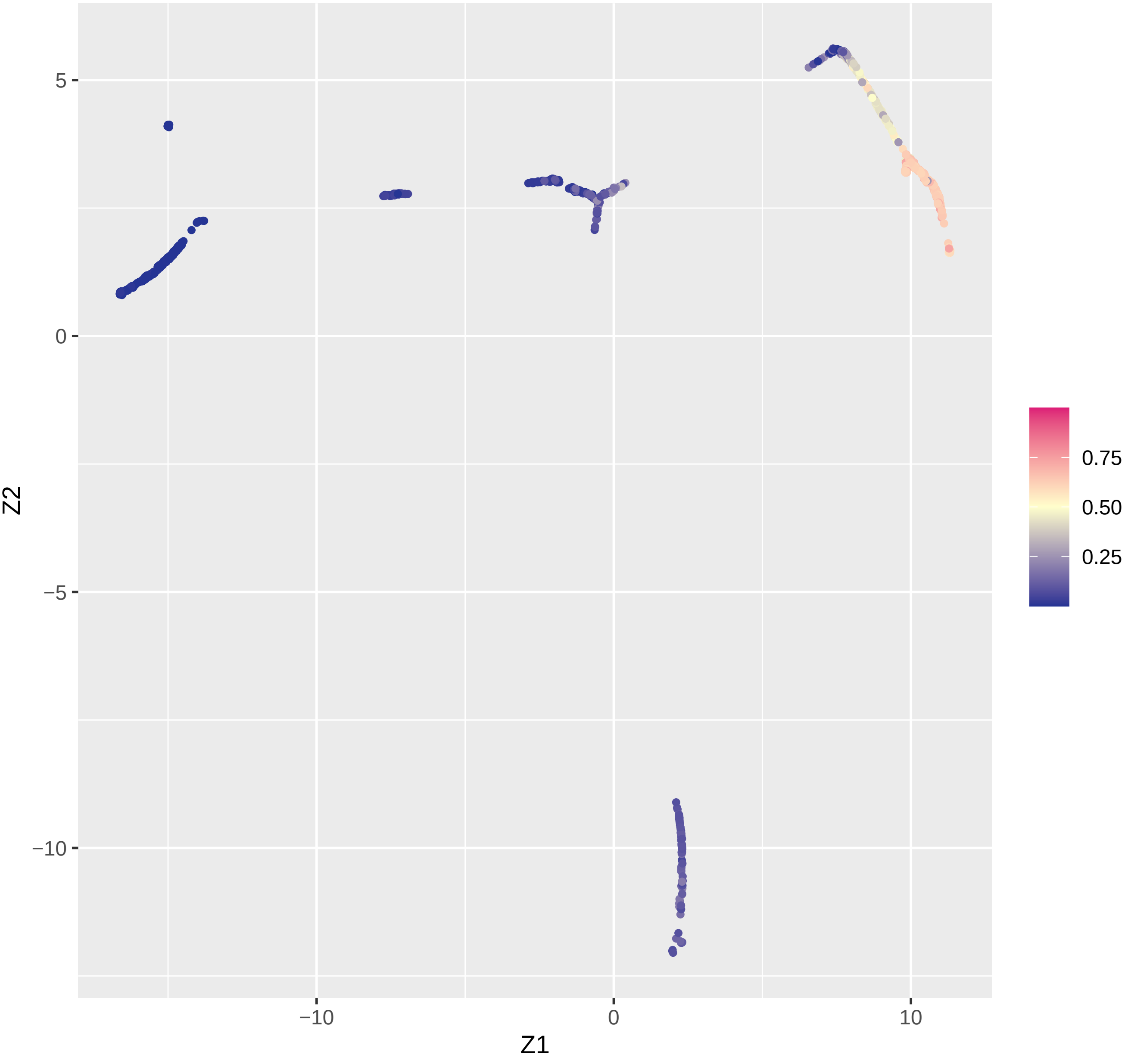

### E11_India

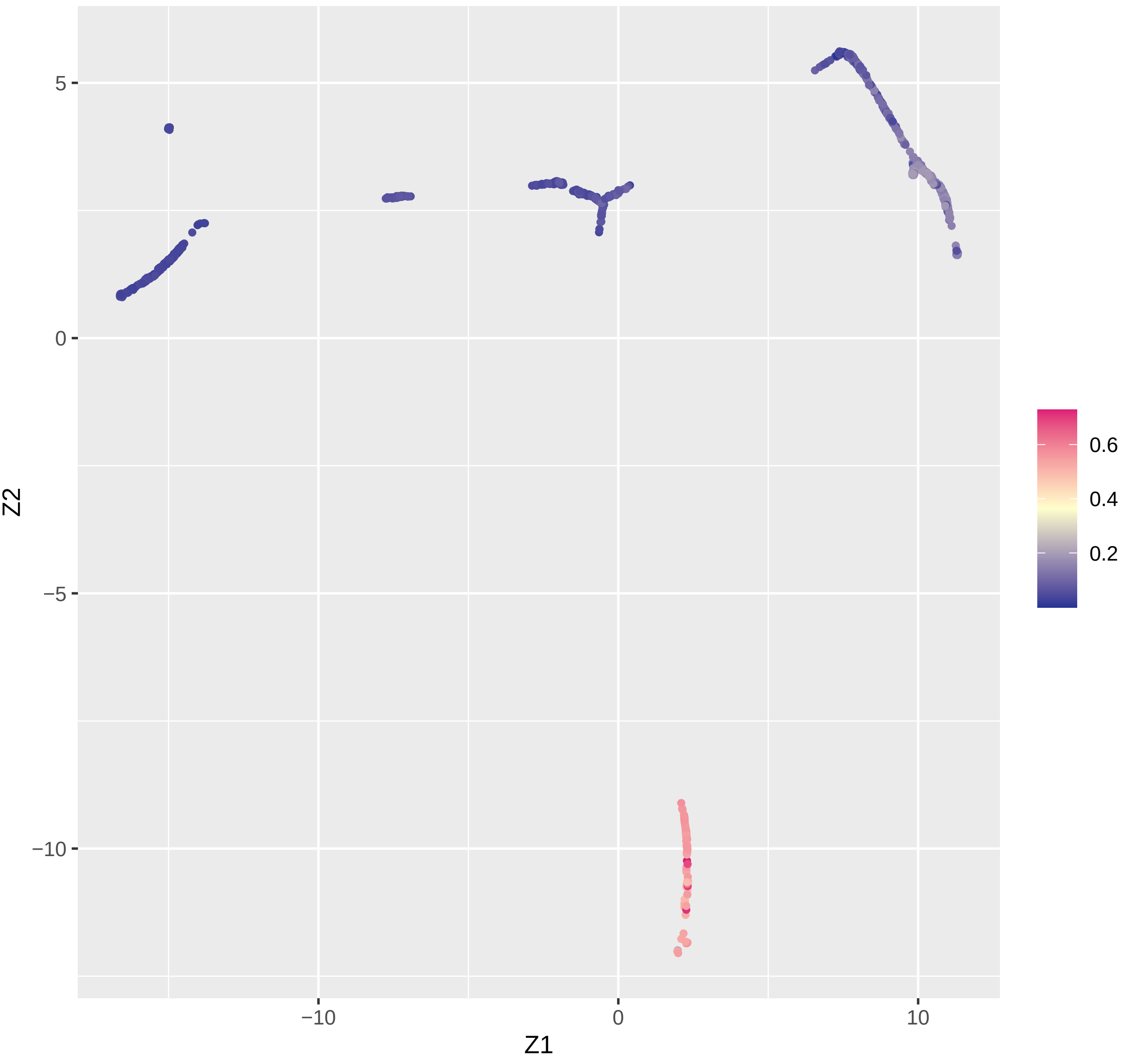

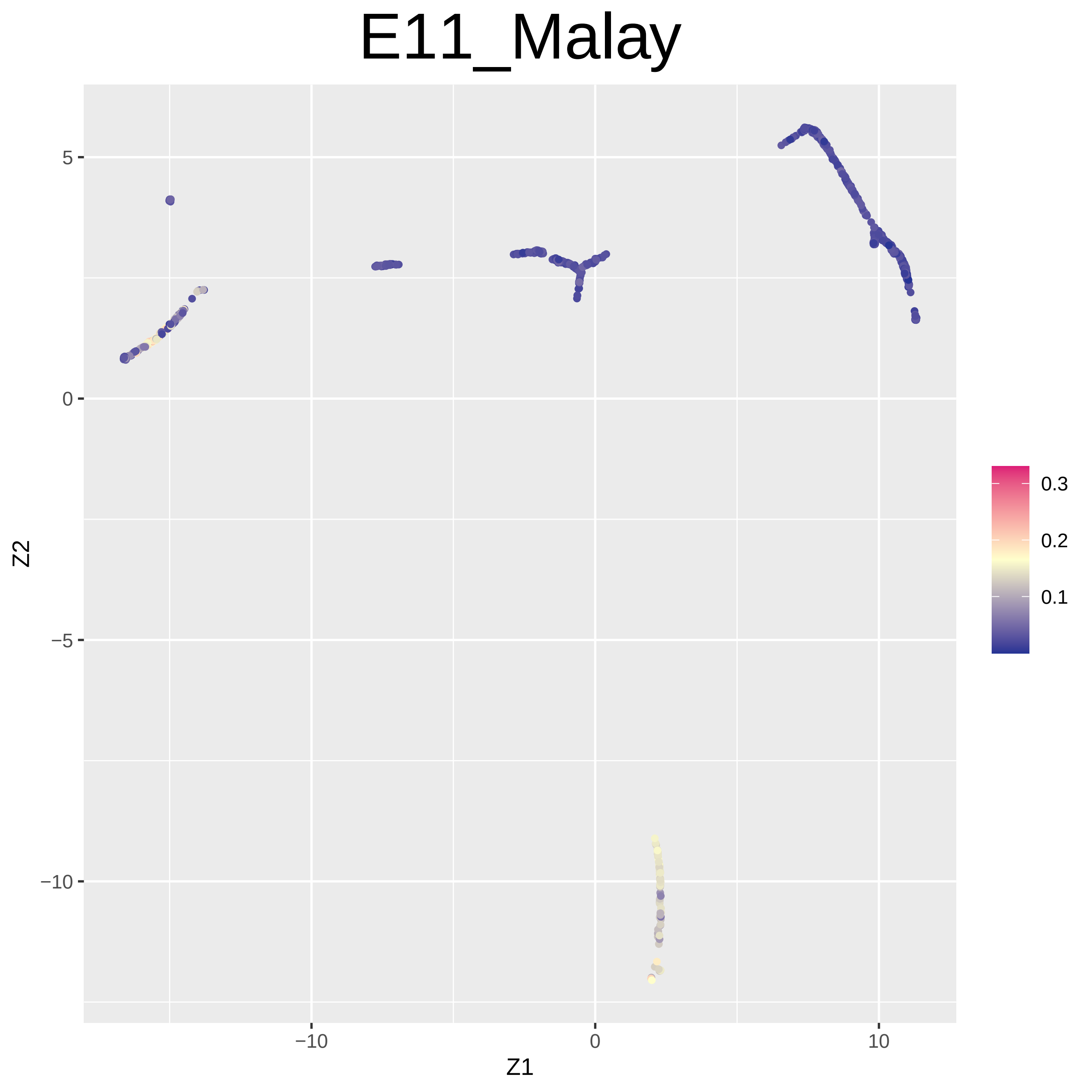
E11_Malay
Z2
Z1
5
0
−5
−10
−10
0
10
0.3
0.2
0.1

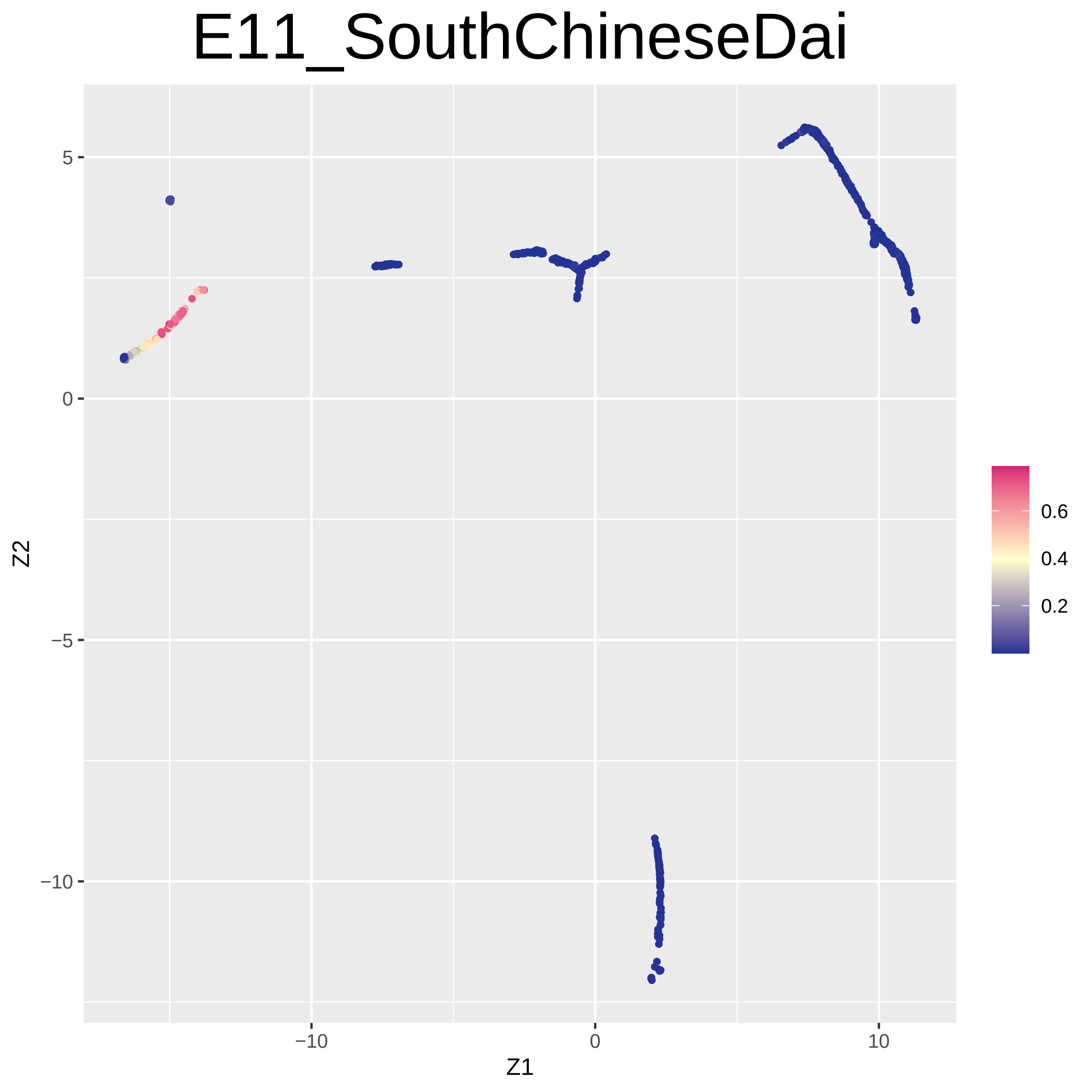

E11_SouthChineseDai
Z2
Z1
5
0
−5
−10
−10
0
10
0.6
0.4
0.2

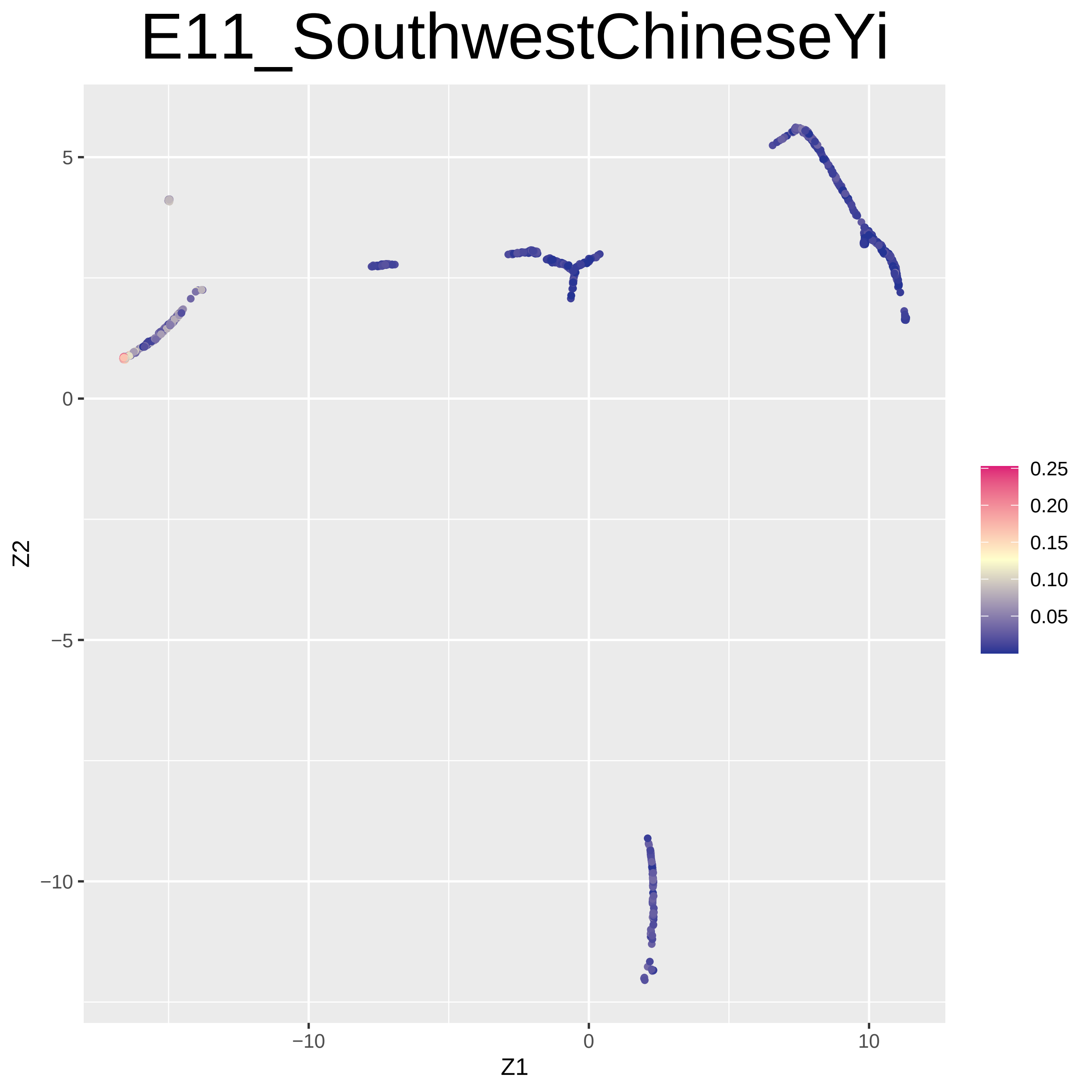
E11_SouthwestChineseYi
Z2
Z1
5
0
−5
−10
−10
0
10
0.25
0.20
0.15
0.10
0.05

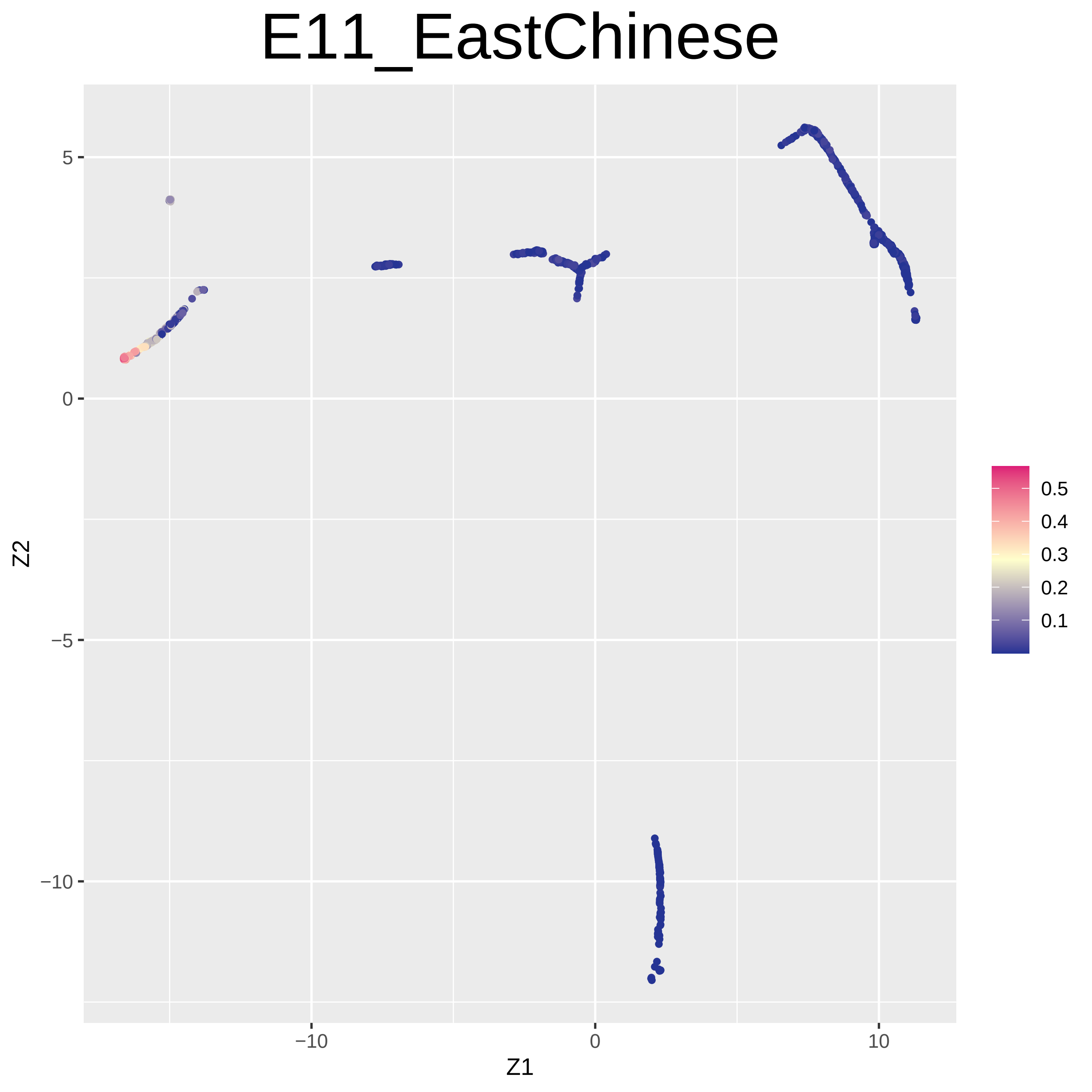
E11_EastChinese
5
0
−5
−10
Z2
−10
0
10
Z1
0.5
0.4
0.3
0.2
0.1

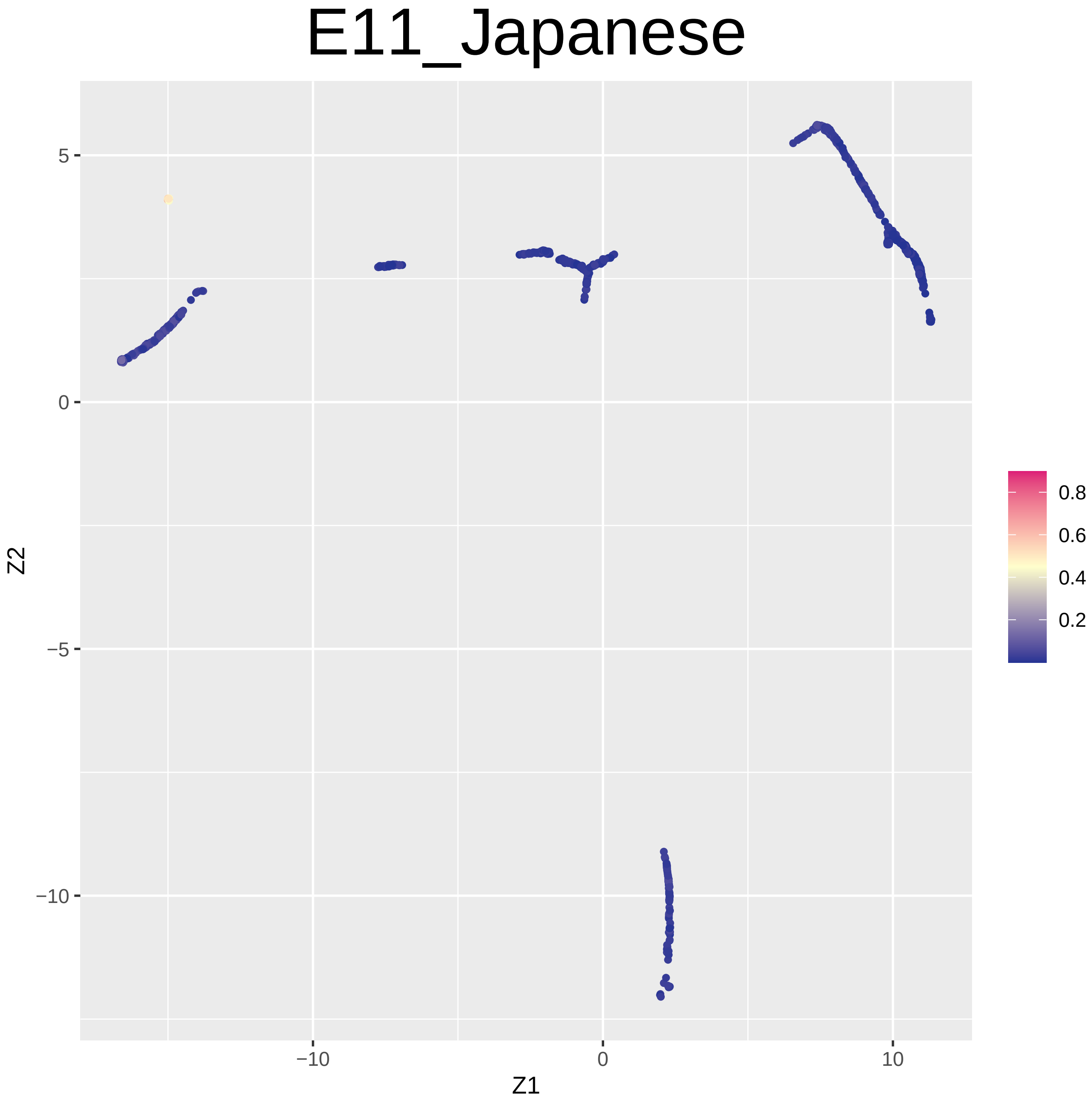

E11_Japanese
5
0
−5
−10
−10
0
10
Z1
Z2
0.8
0.6
0.4
0.2

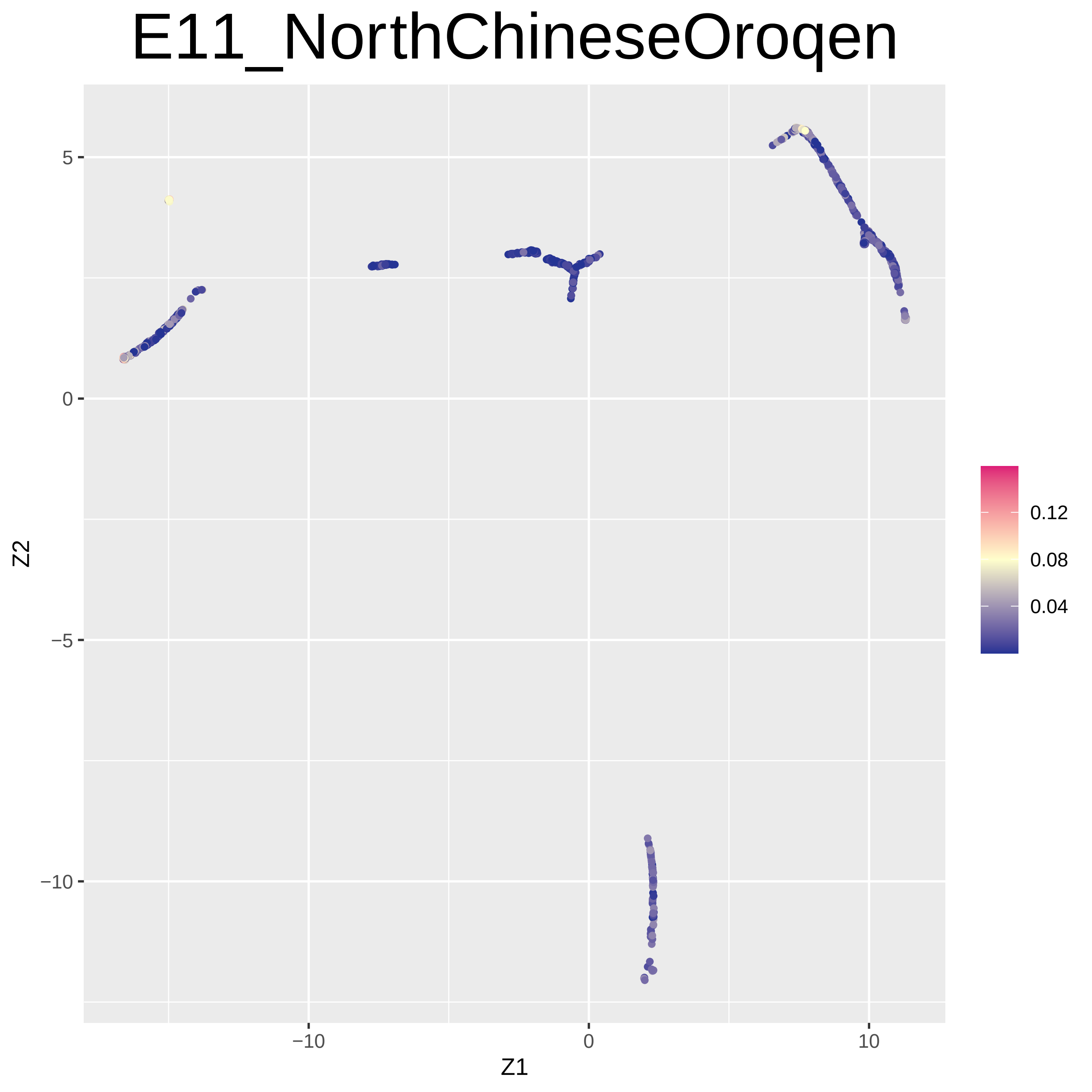

E11_NorthChineseOroqen
Z2
Z1
5
0
−5
−10
−10
0
10
0.12
0.08
0.04

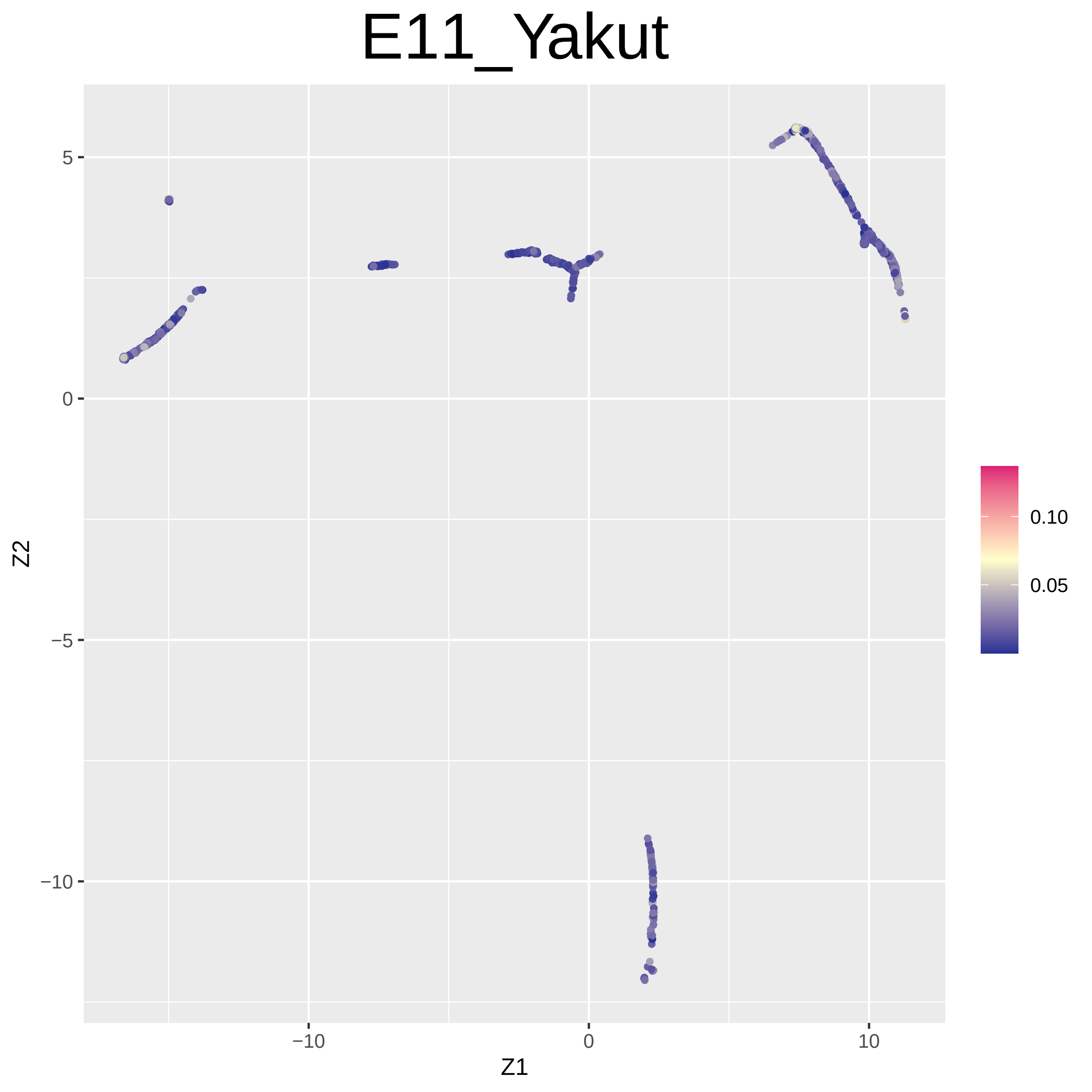
E11_Yakut
5
0
−5
−10
Z2
−10
0
10
Z1
0.10
0.05

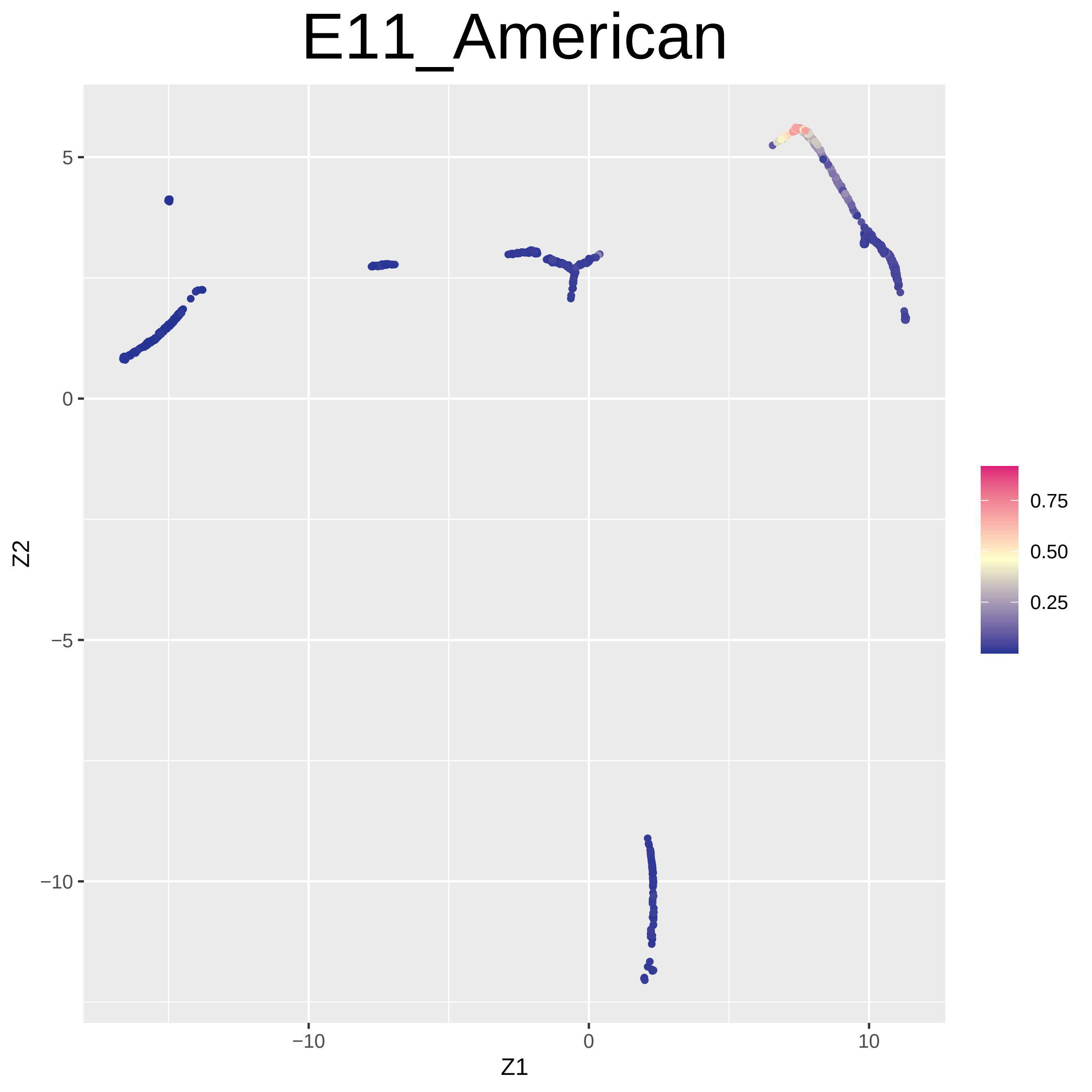

E11_American
5
0
−5
−10
−10
0
10
Z1
Z2
0.75
0.50
0.25

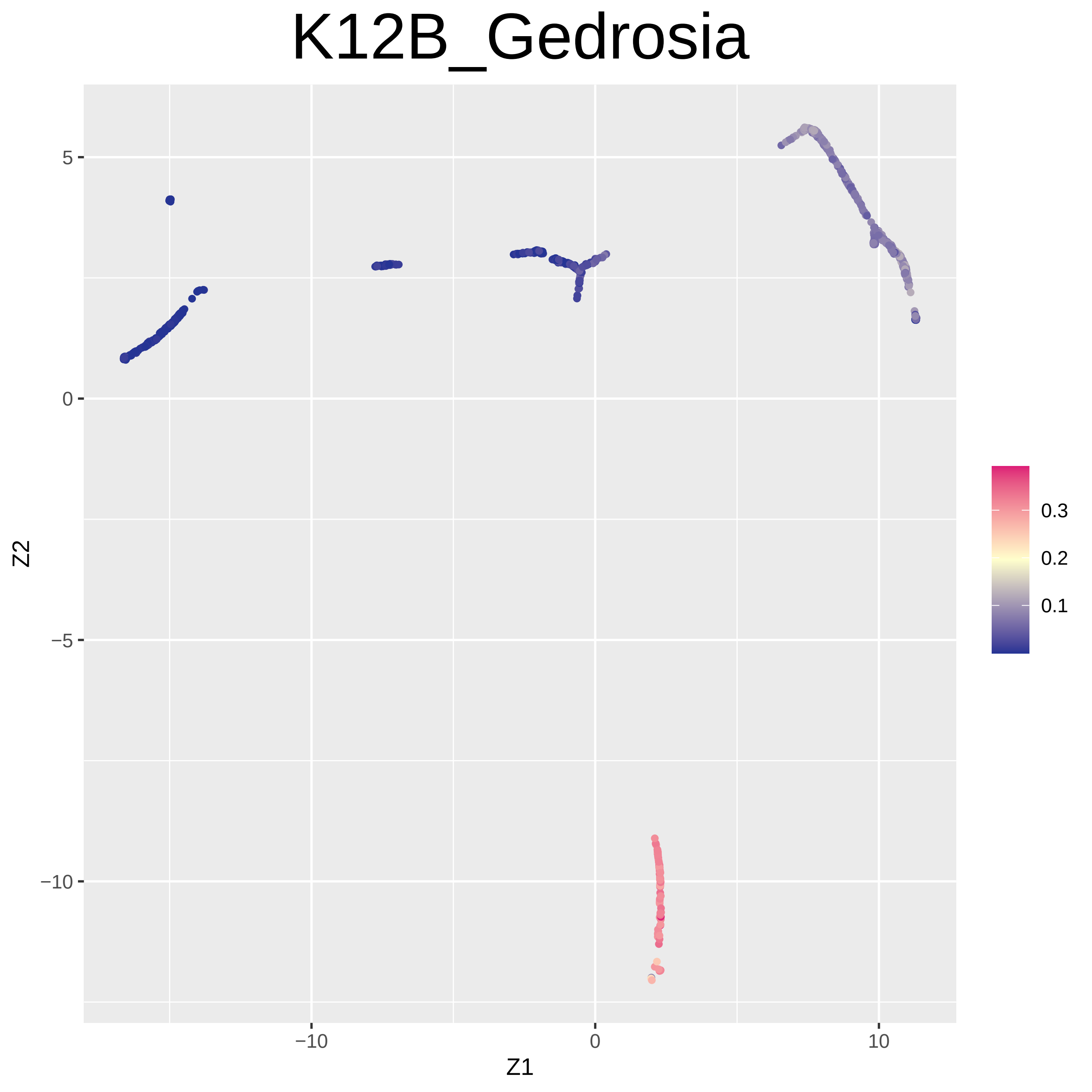

K12B_Gedrosia
5
0
−5
−10
−10
0
10
Z1
Z2
0.3
0.2
0.1

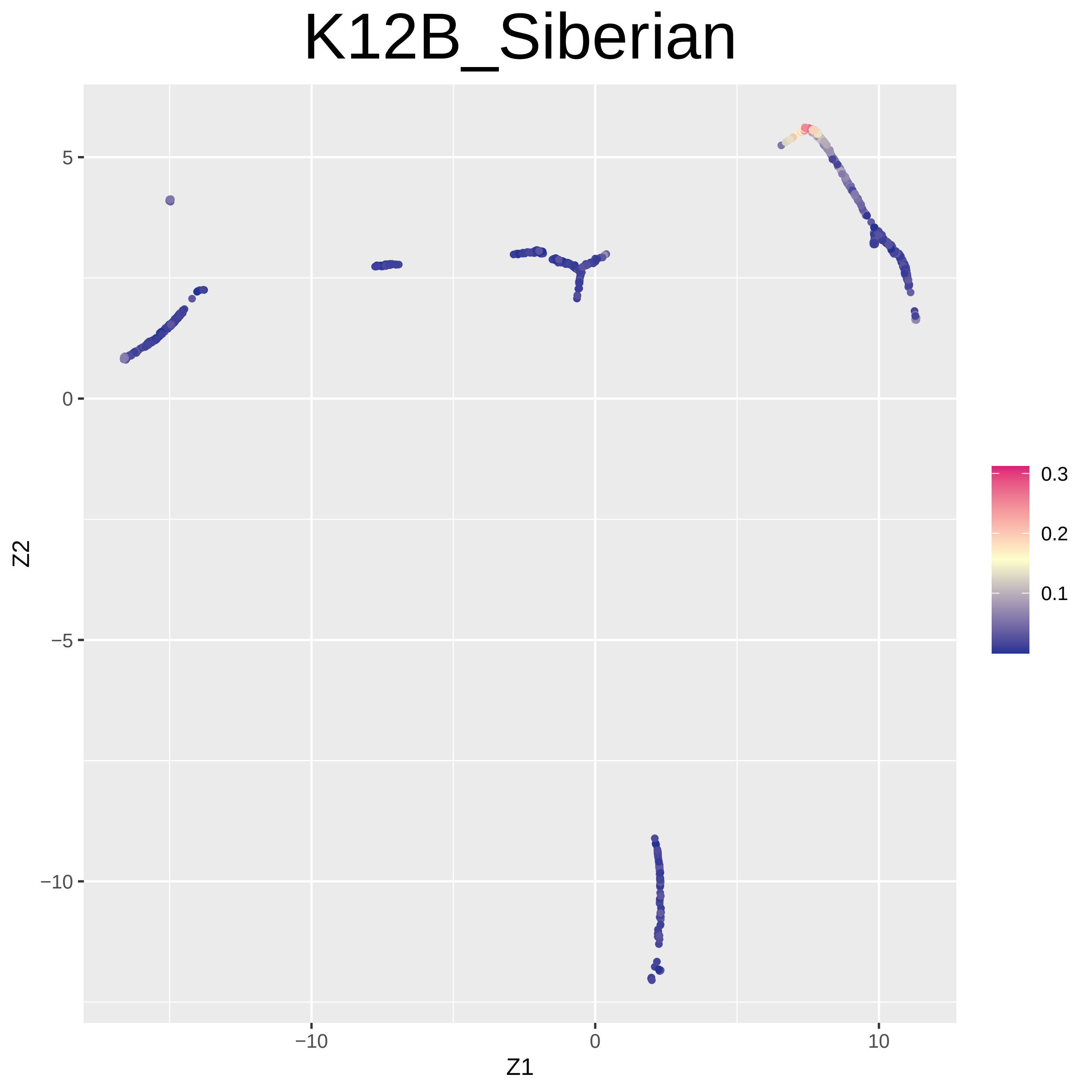

K12B_Siberian
Z2
Z1
5
0
−5
−10
−10
0
10
0.3
0.2
0.1

### K12B_Northwest_African

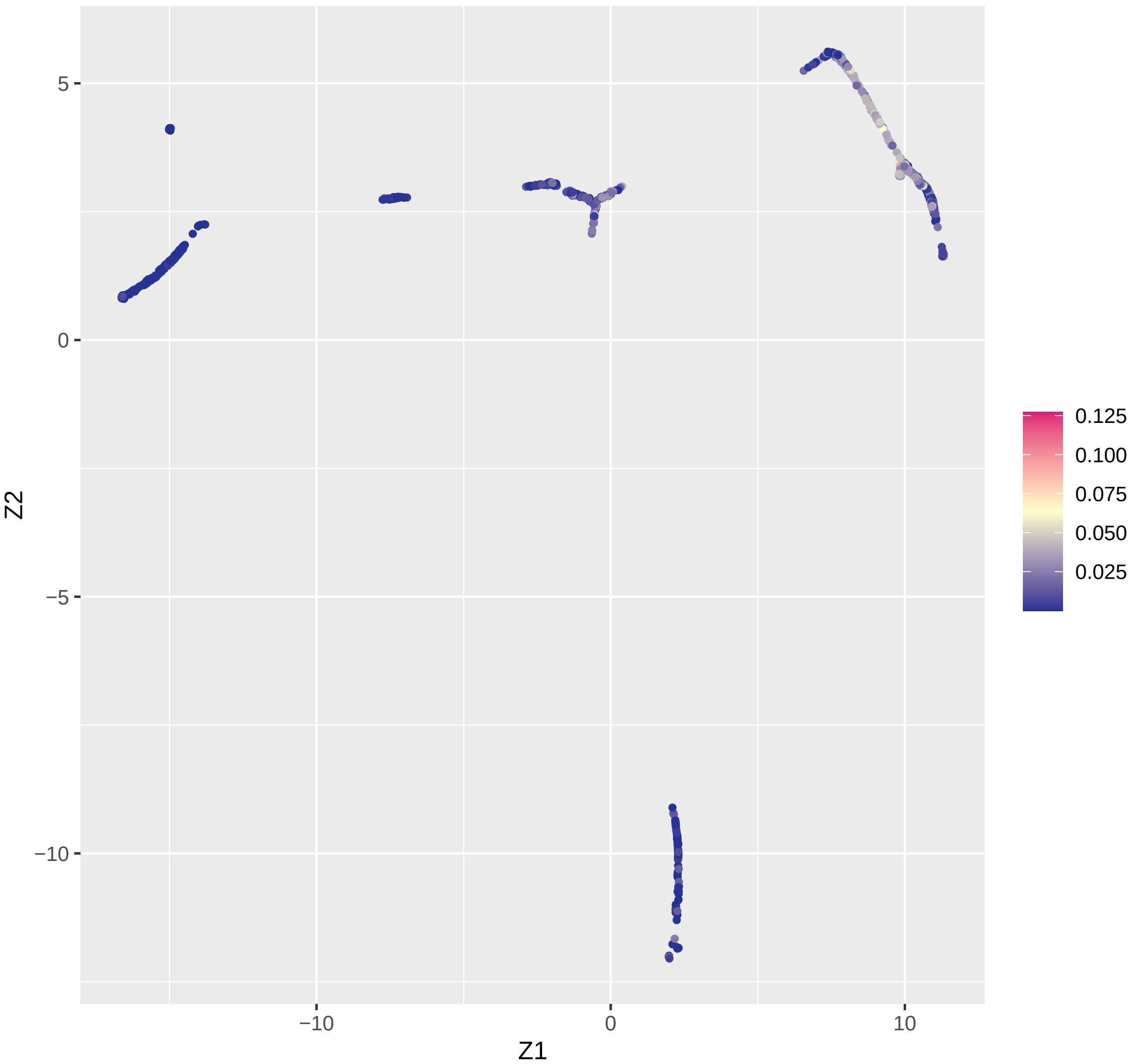

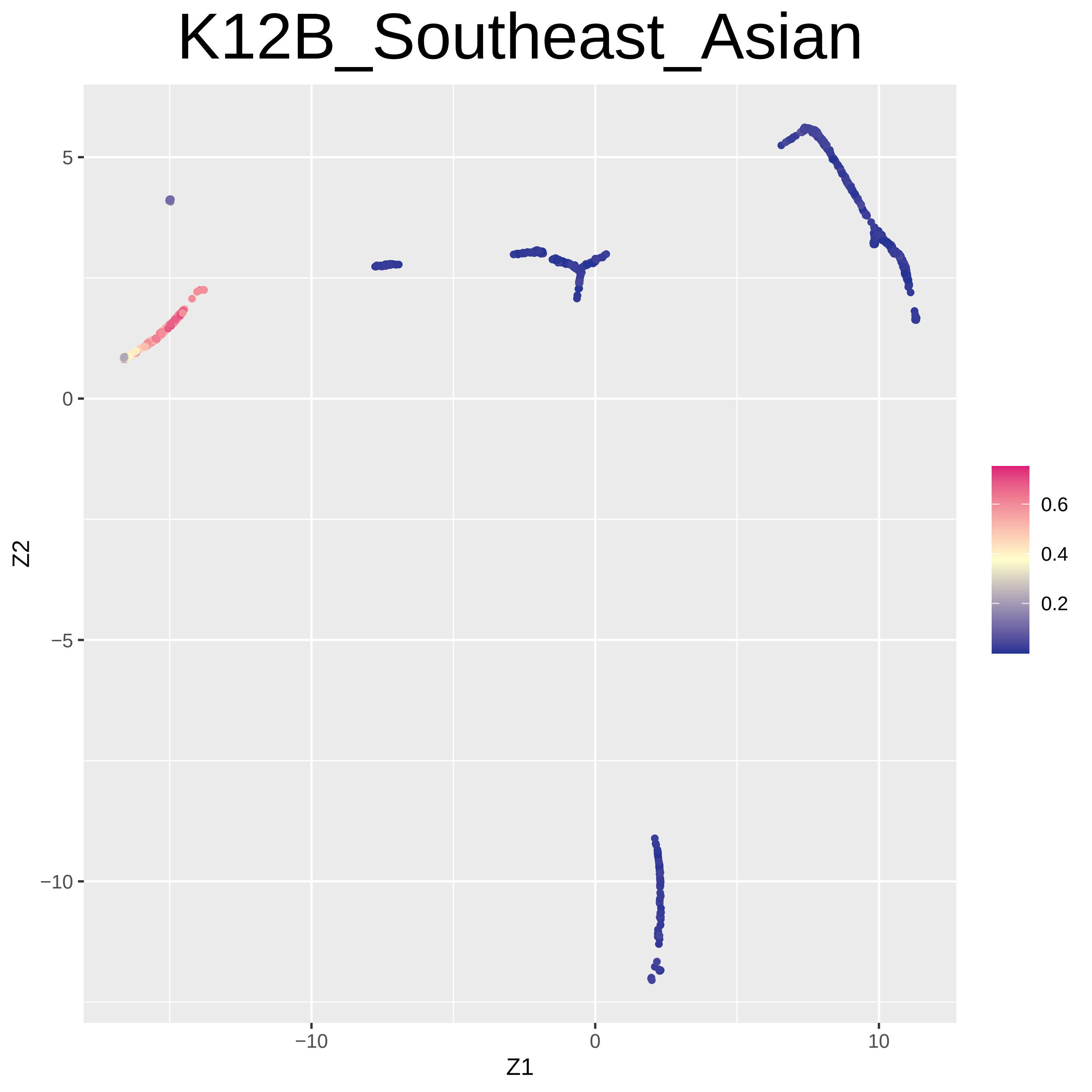

K12B_Southeast_Asian
Z2
Z1
5
0
−5
−10
−10
0
10
0.6
0.4
0.2

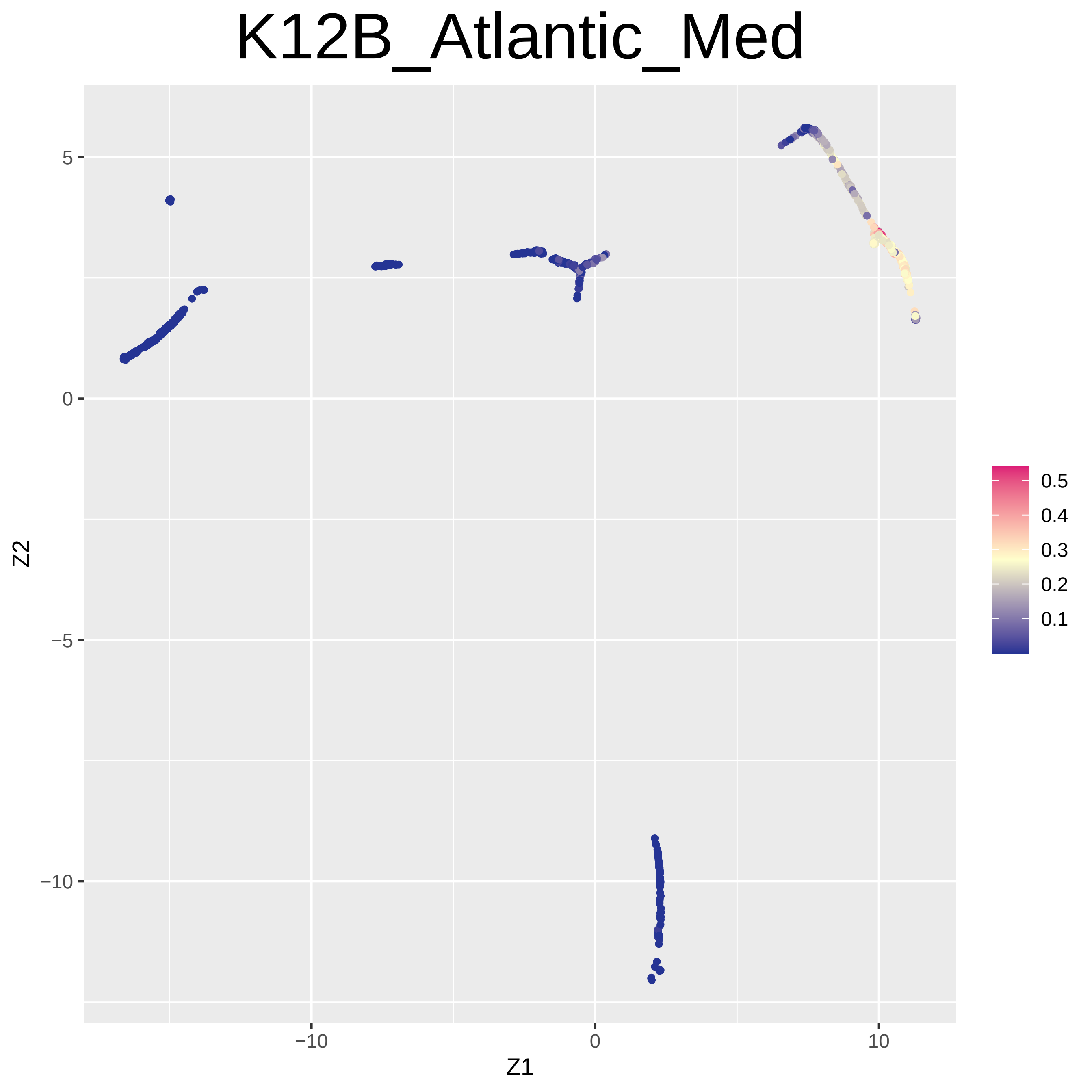
K12B_Atlantic_Med
Z2
5
0
−5
−10
−10
0
10
Z1
0.5
0.4
0.3
0.2
0.1

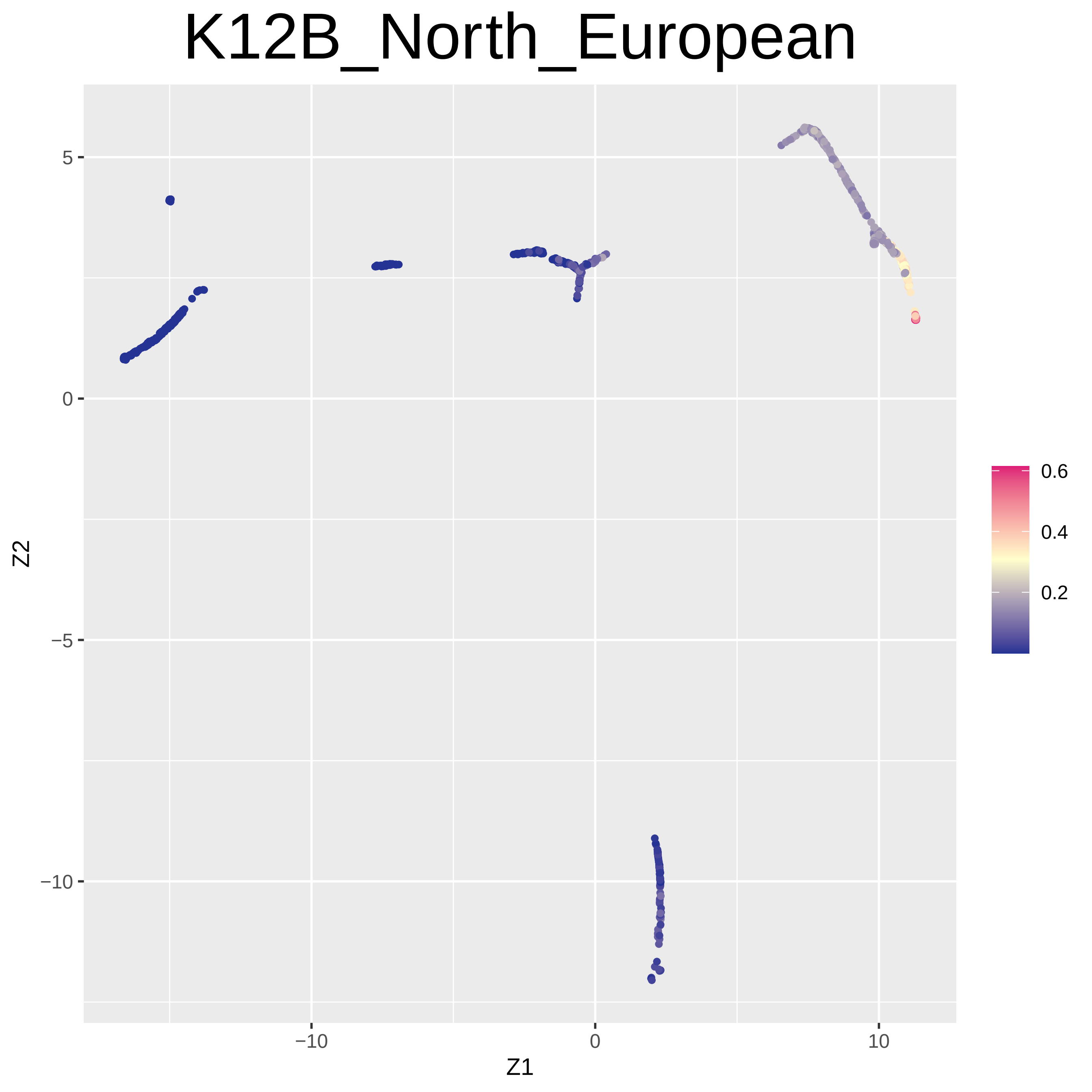
K12B_North_European
Z1
Z2
5
0
−5
−10
−10
0
10
0.6
0.4
0.2

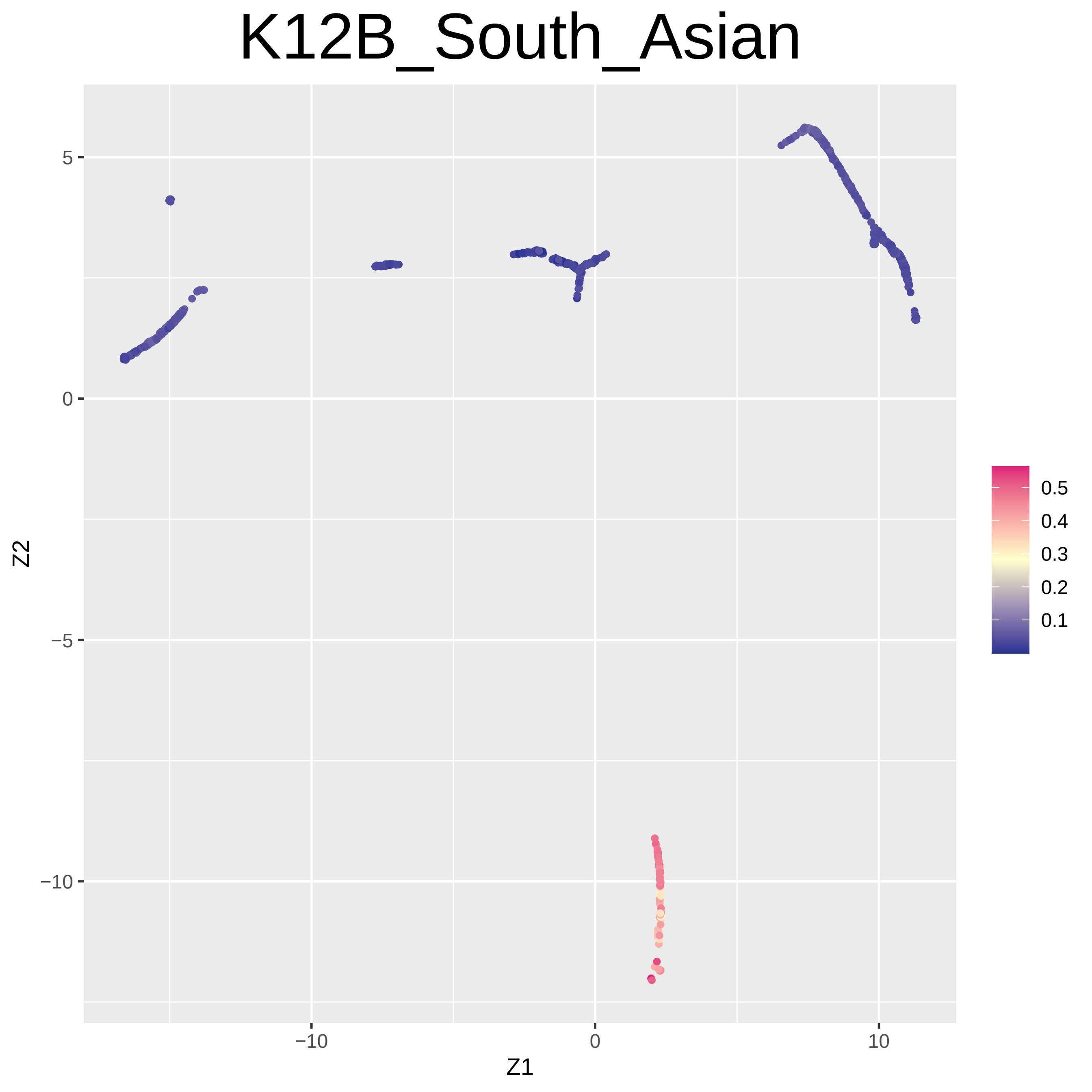
K12B_South_Asian
Z2
Z1
5
0
−5
−10
−10
0
10
0.5
0.4
0.3
0.2
0.1

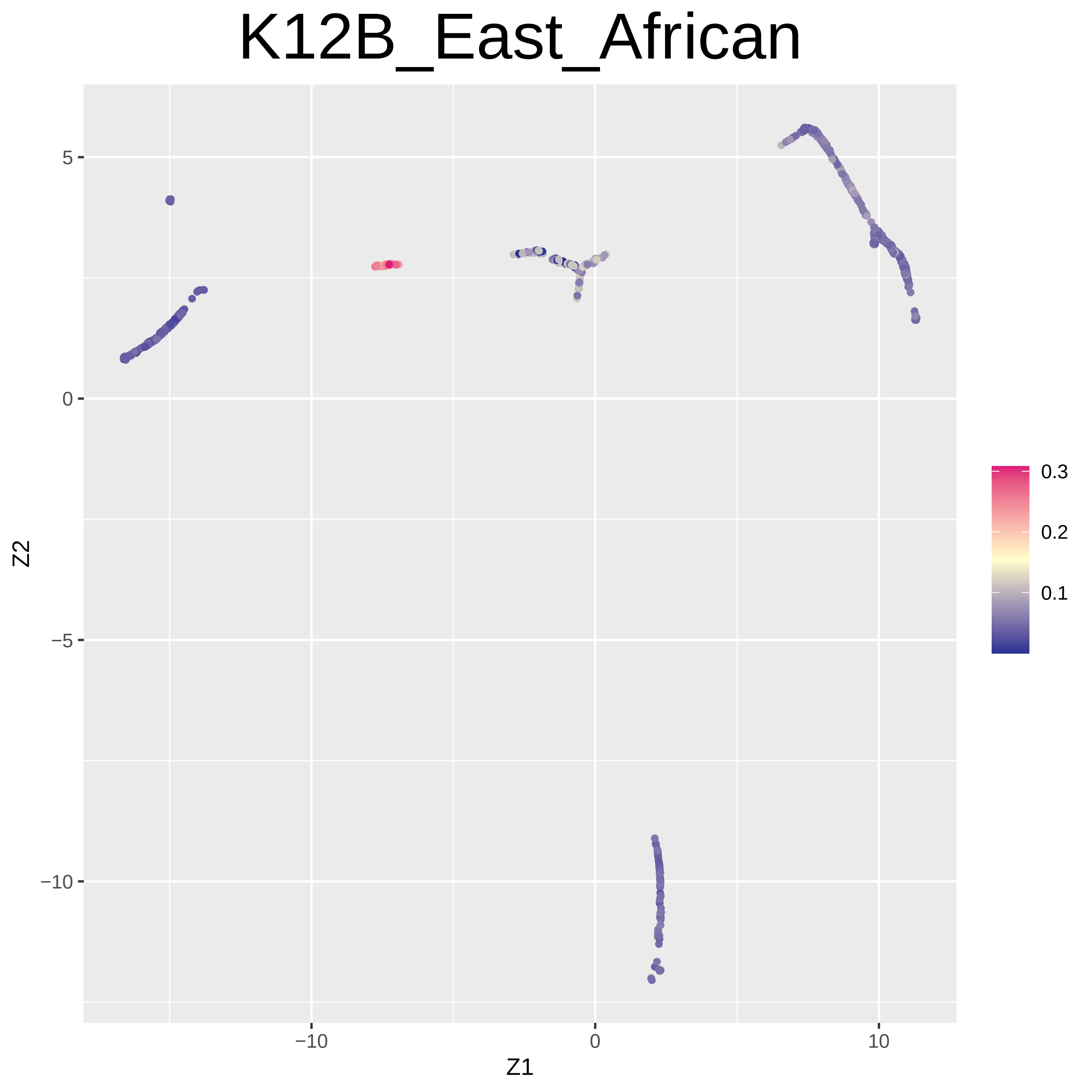
K12B_East_African
5
0
−5
−10
Z2
−10
0
10
Z1
0.3
0.2
0.1

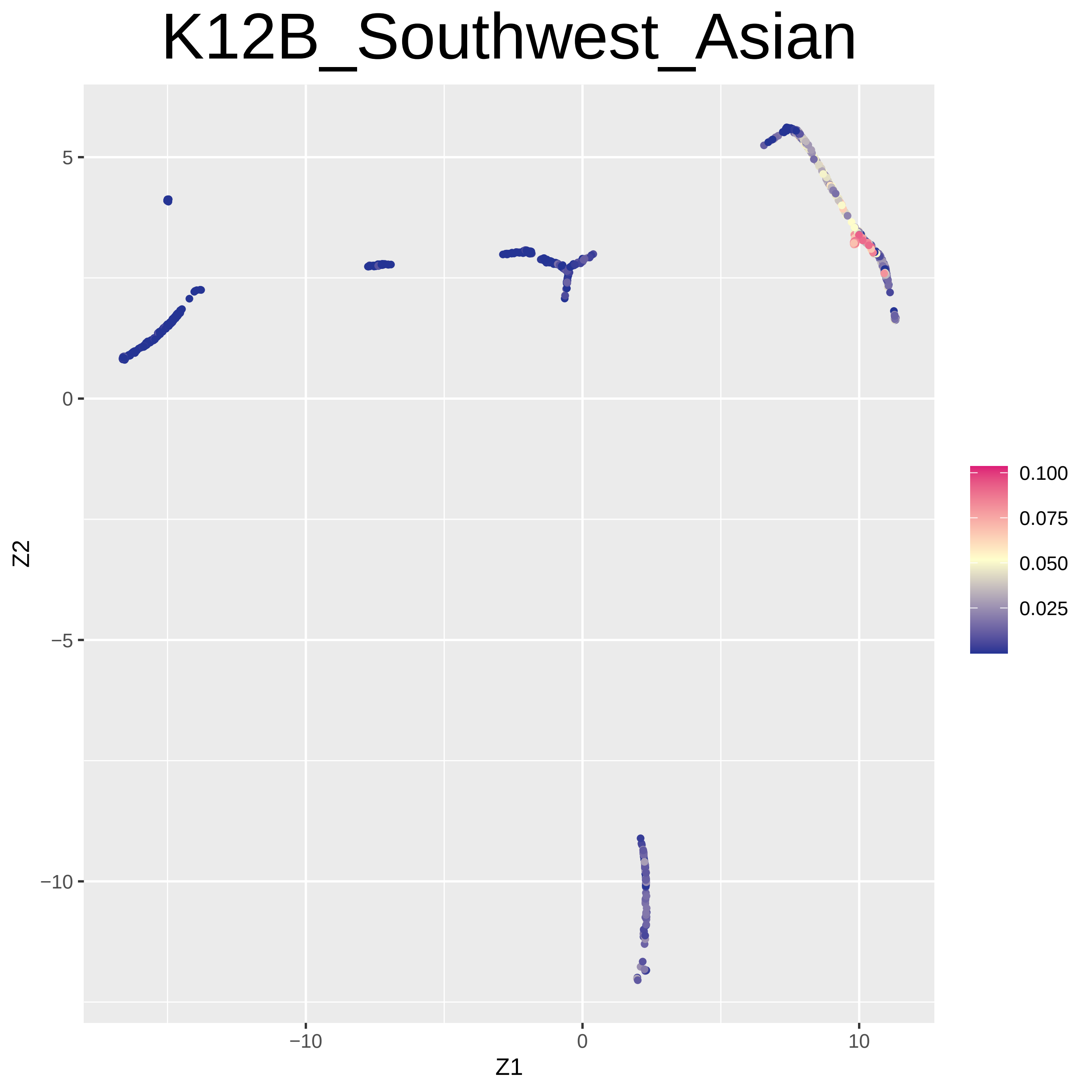
K12B_Southwest_Asian
Z2
Z1
5
0
−5
−10
−10
0
10
0.100
0.075
0.050
0.025

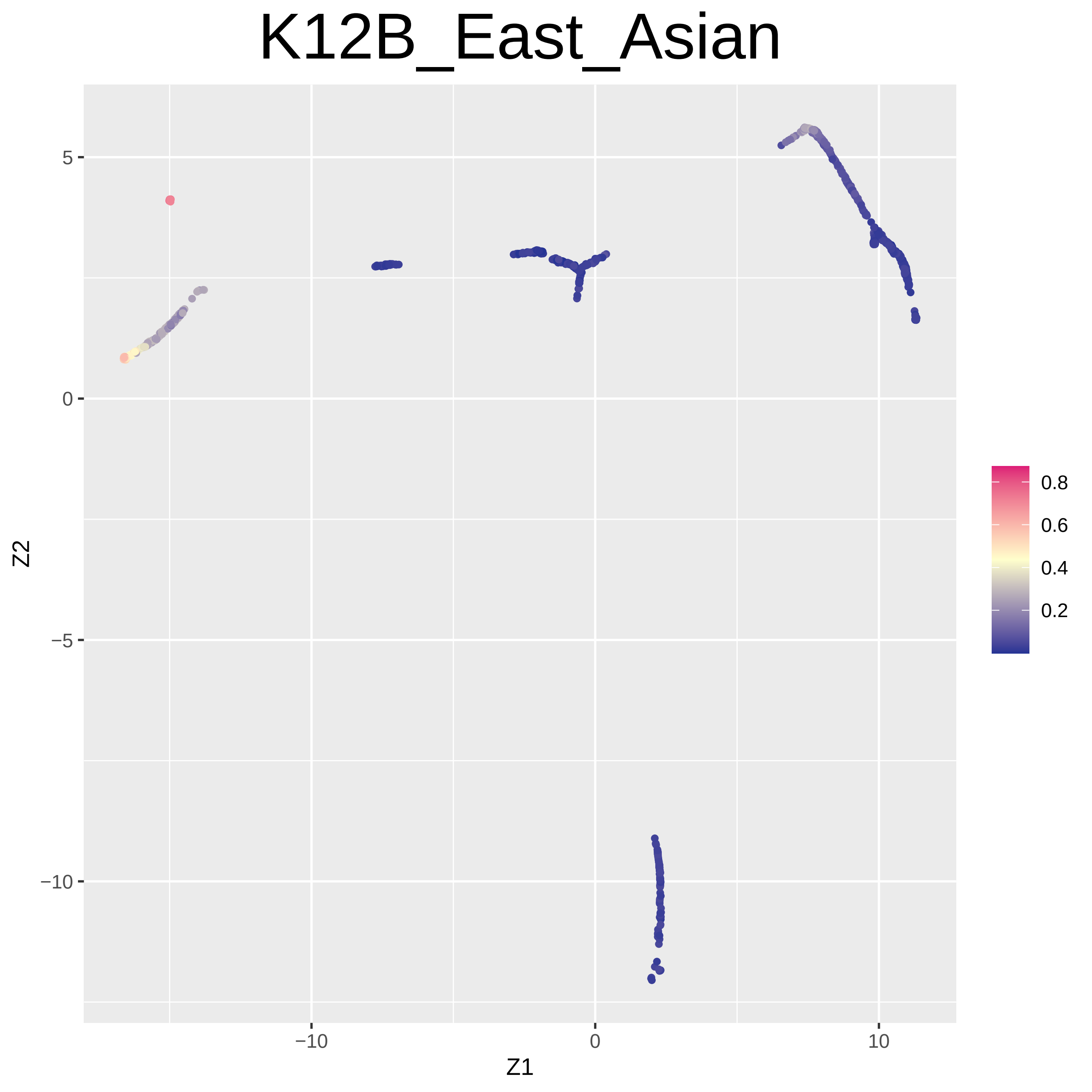
K12B_East_Asian
Z2
Z1
5
0
−5
−10
−10
0
10
0.8
0.6
0.4
0.2

### K12B_Caucasus

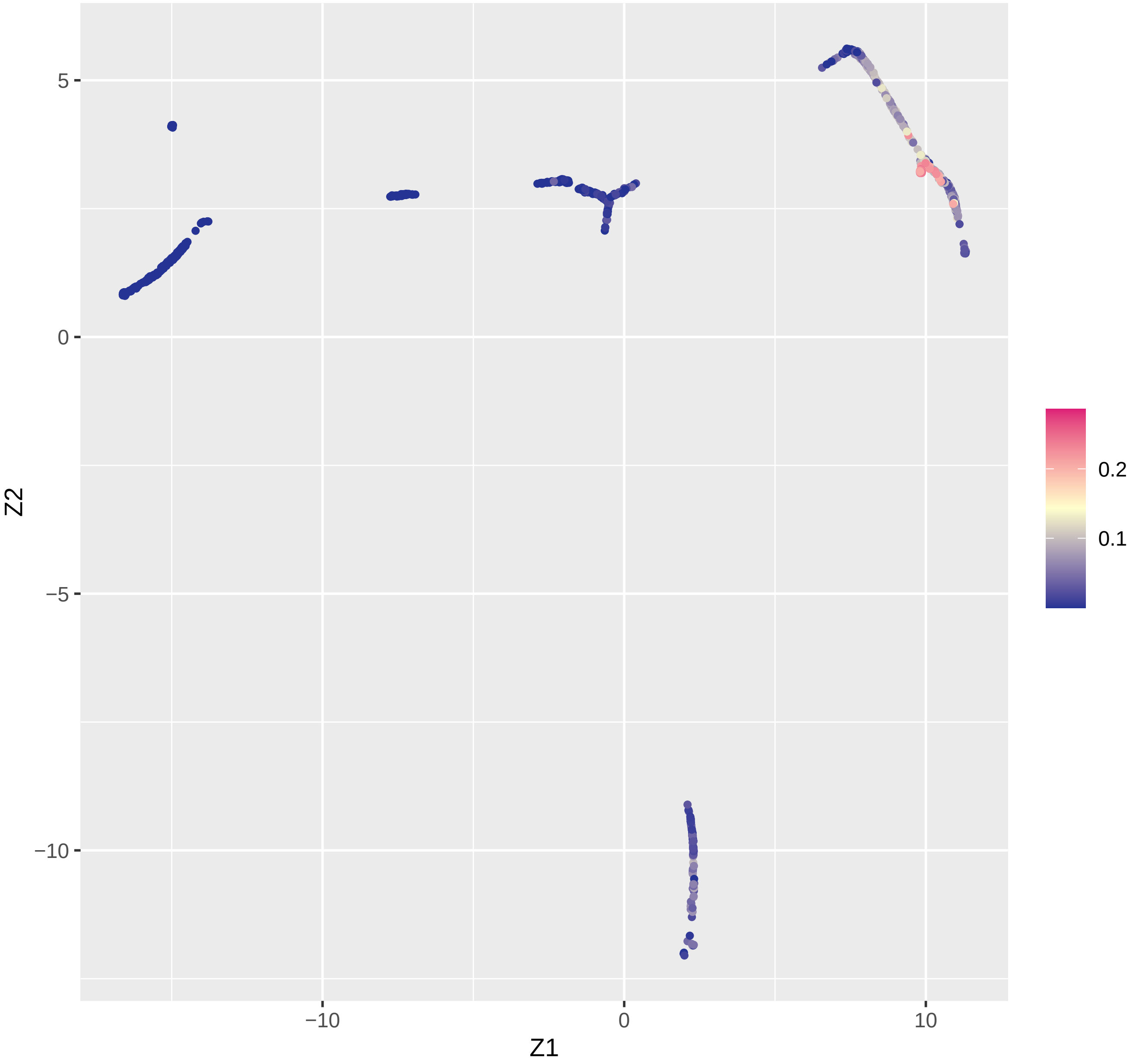

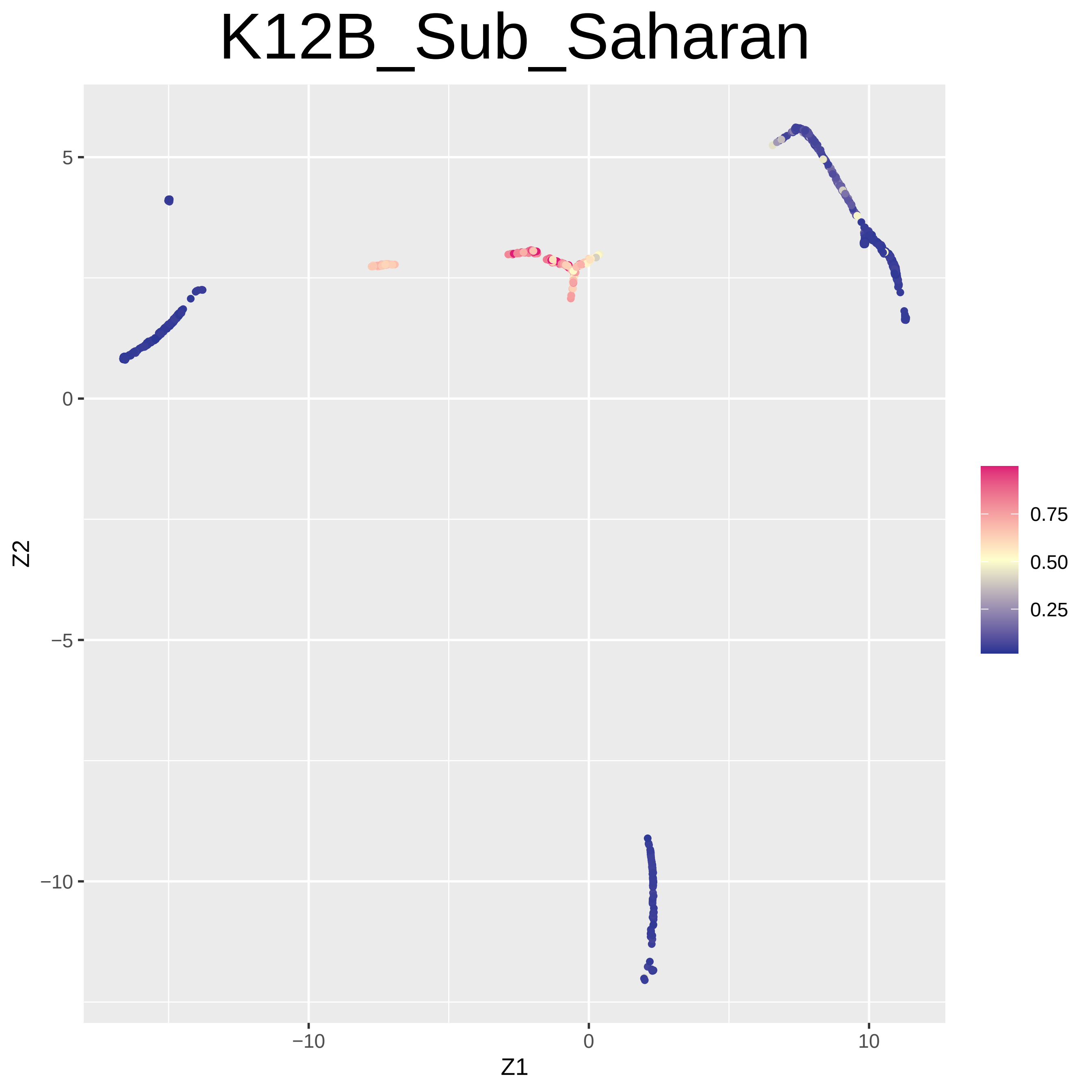
K12B_Sub_Saharan
Z2
Z1
5
0
−5
−10
−10
0
10
0.75
0.50
0.25
