## Supplementary Figures 5 for "Graph Embedding Method Based Genetical Trajectory Reveals Migration History Among East Asians"

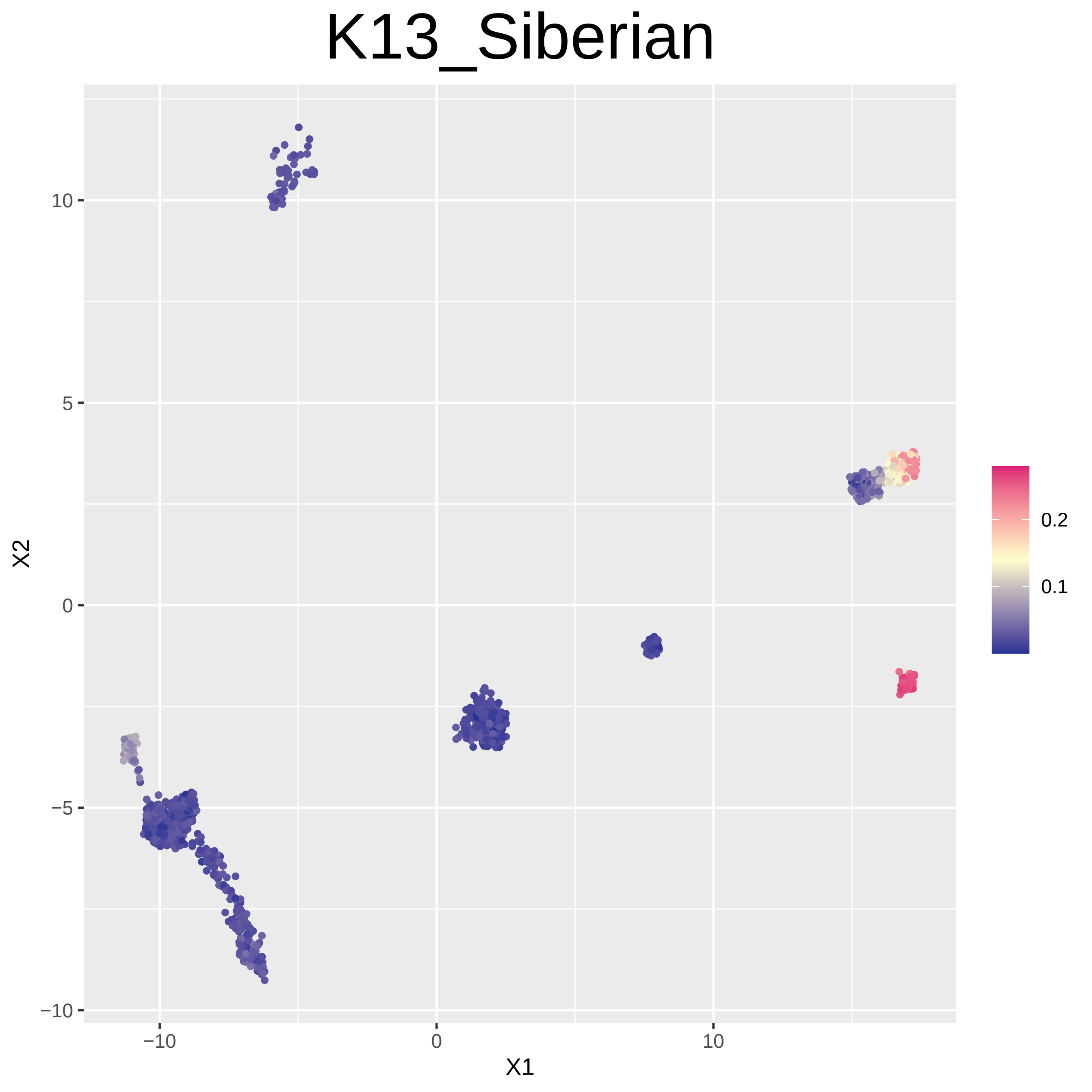

K13_Siberian
10
5
0
−5
−10
X2
−10
0
10
X1
0.2
0.1

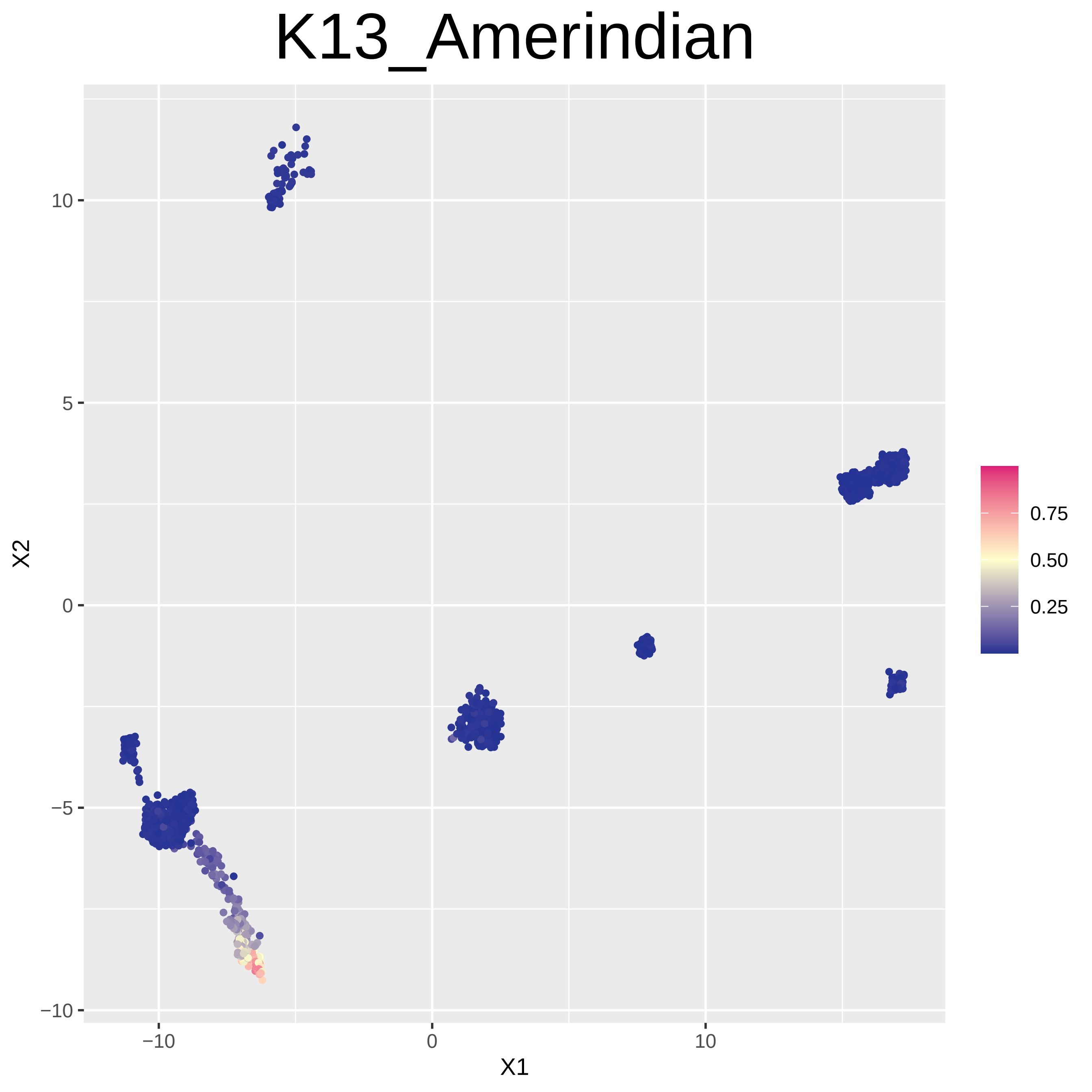
K13_Amerindian
X1
X2
−10
0
10
−10
−5
0
5
10
0.75
0.50
0.25

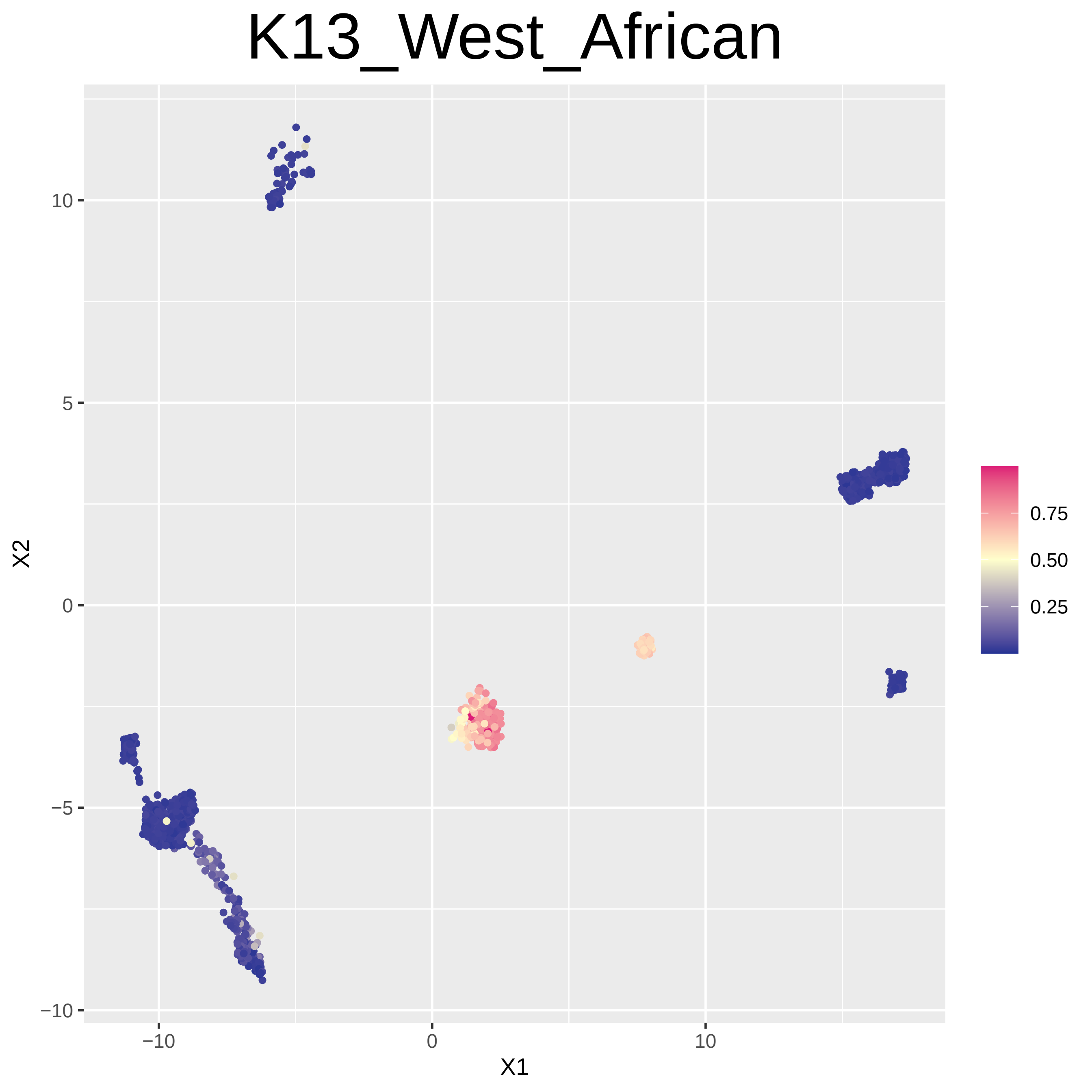

K13_West_African
X2
X1
10
5
0
−5
−10
−10
0
10
0.75
0.50
0.25

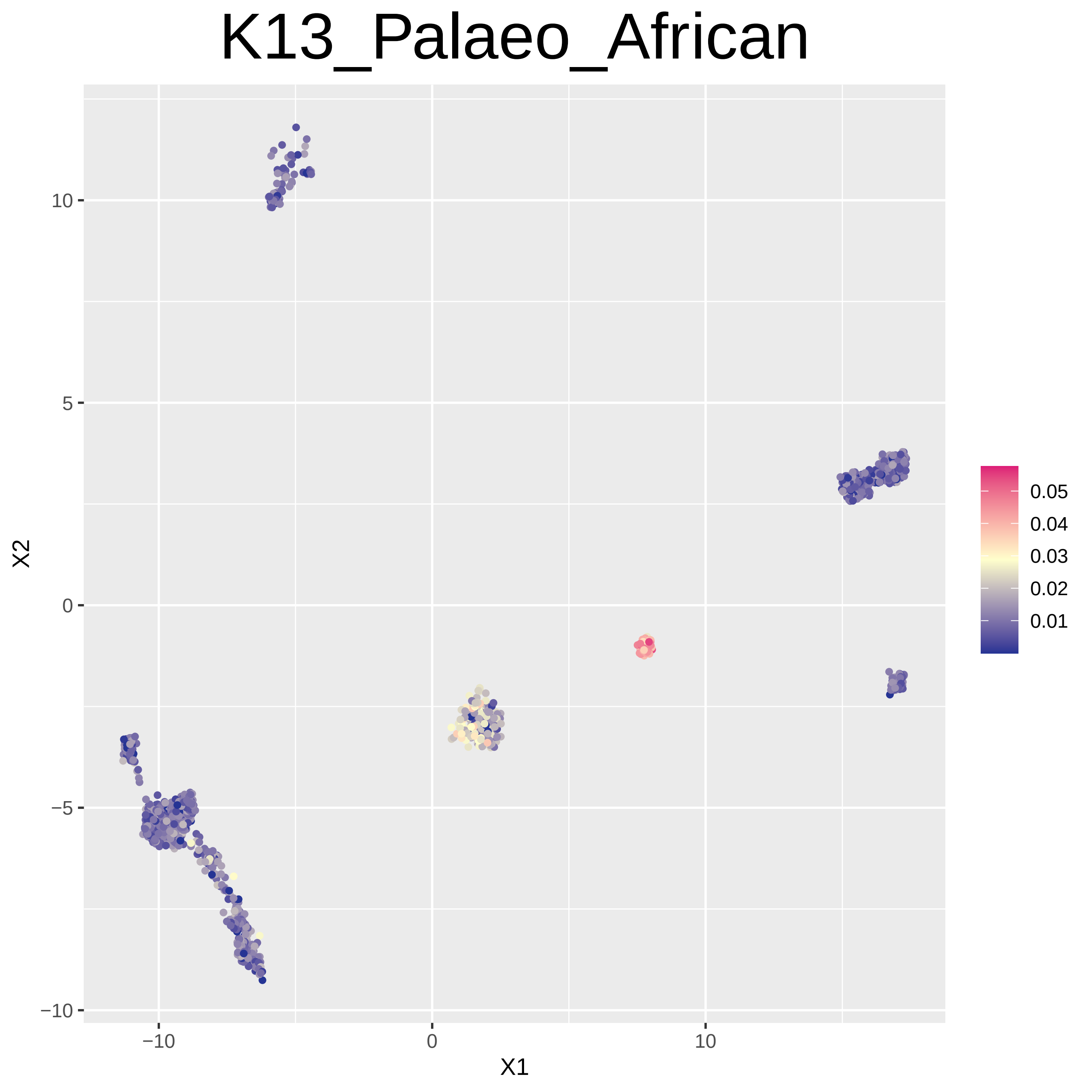
K13_Palaeo_African
X2
X1
10
5
0
−5
−10
−10
0
10
0.05
0.04
0.03
0.02
0.01

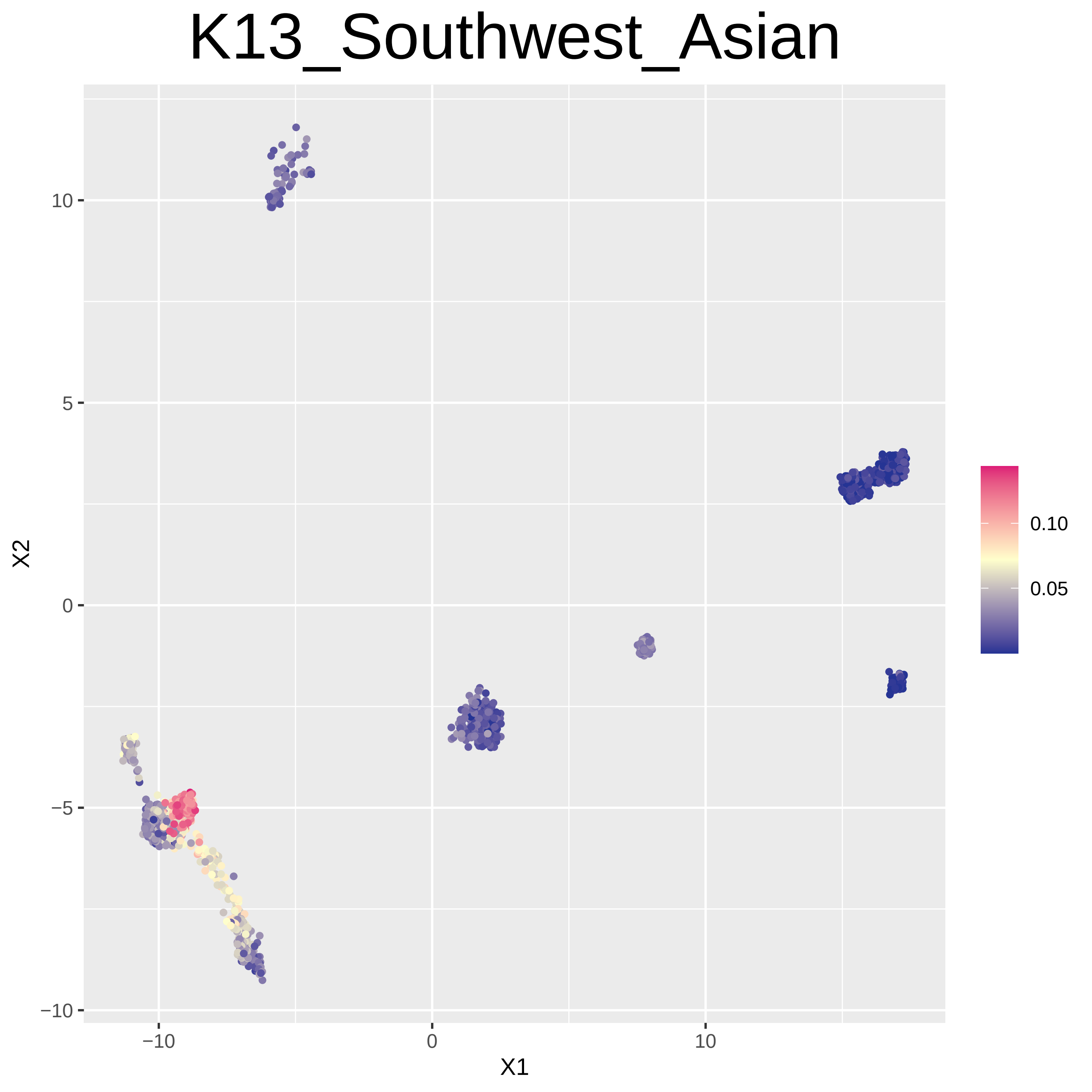

K13_Southwest_Asian
X2
X1
10
5
0
−5
−10
−10
0
10
0.10
0.05

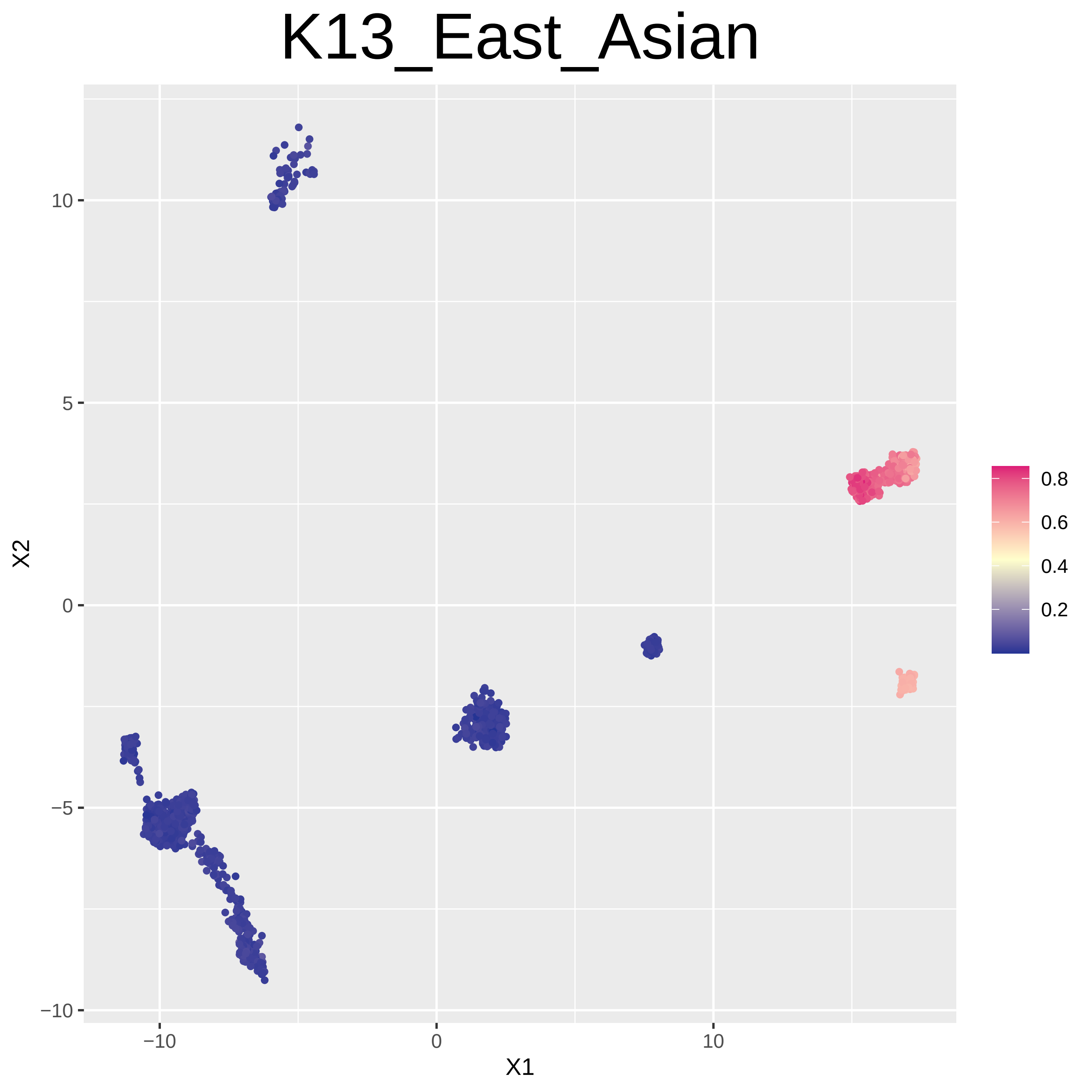
K13_East_Asian
X2
X1
10
5
0
−5
−10
−10
0
10
0.8
0.6
0.4
0.2

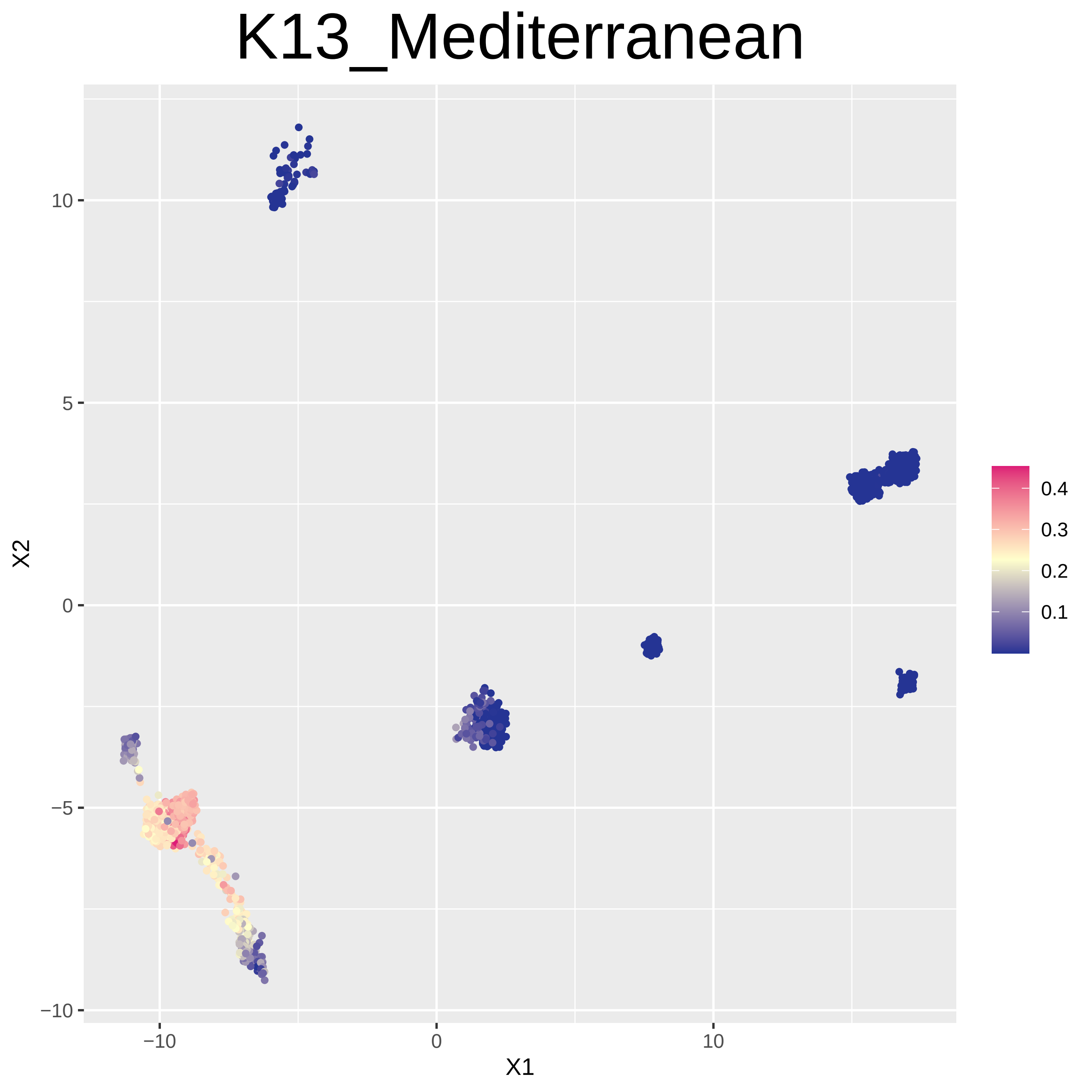
K13_Mediterranean
10
5
0
−5
−10
−10
0
10
X1
X2
0.4
0.3
0.2
0.1

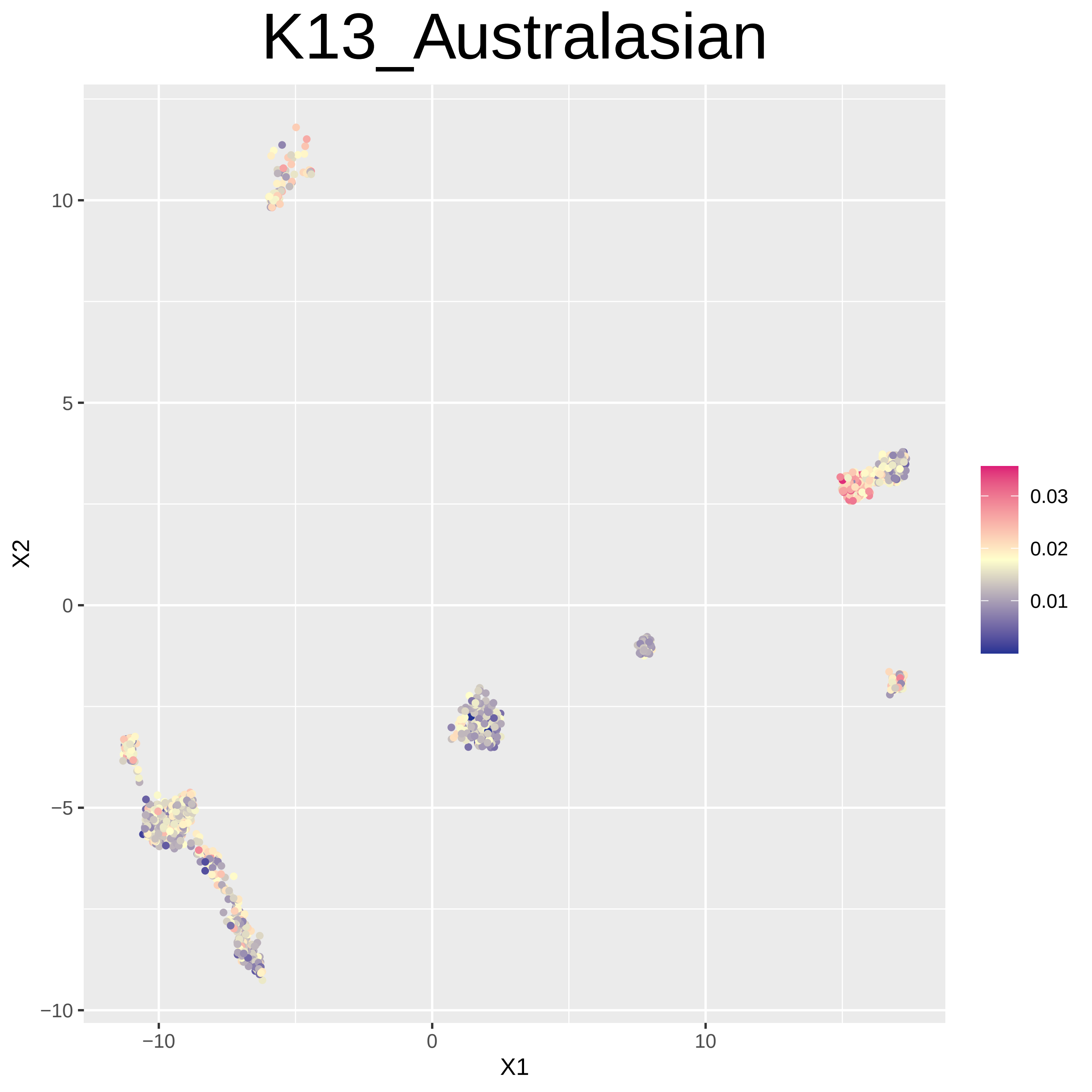
K13_Australasian
X1
X2
−10
0
10
−10
−5
0
5
10
0.03
0.02
0.01

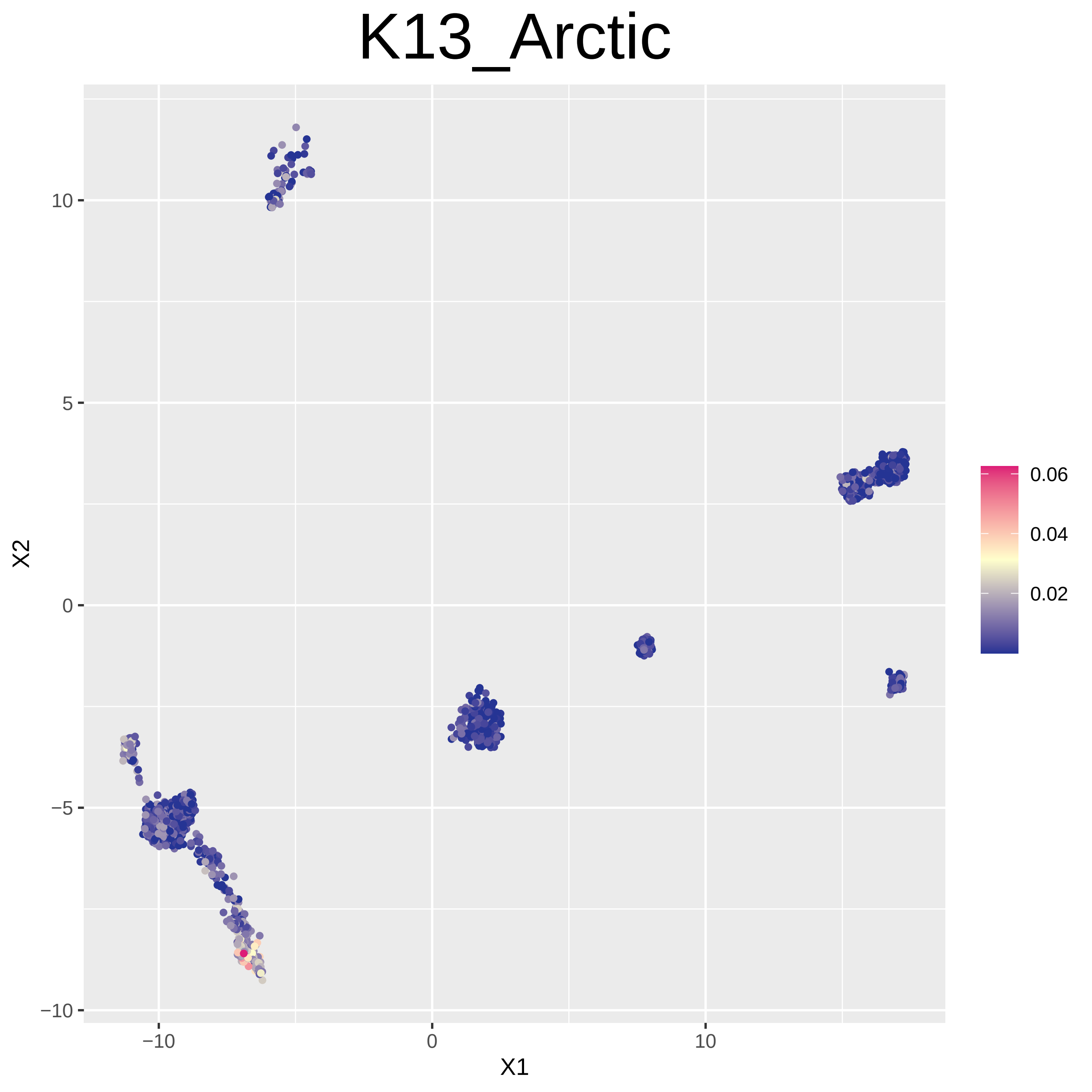

K13_Arctic
X2
X1
10
5
0
−5
−10
−10
0
10
0.06
0.04
0.02

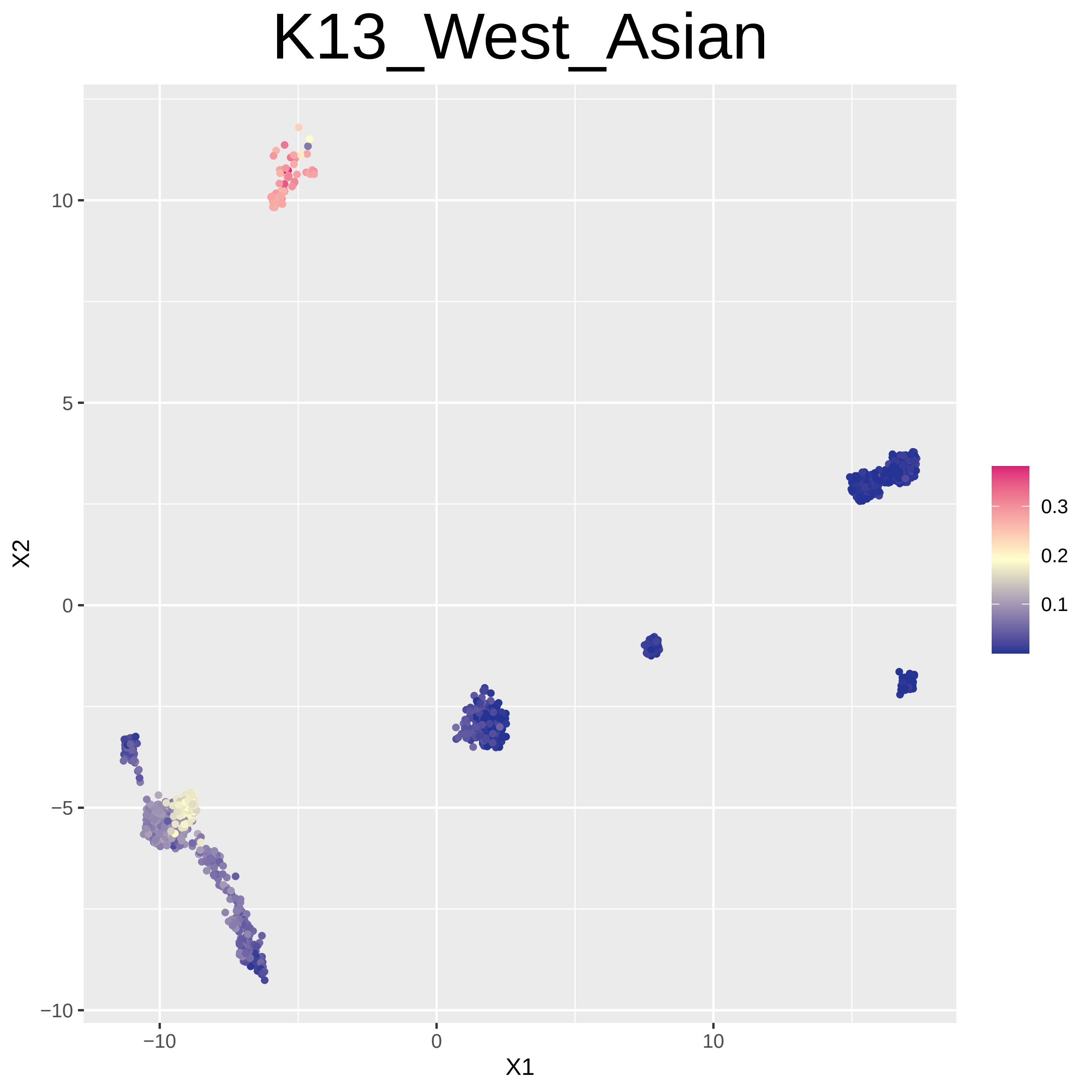
K13_West_Asian
X1
X2
−10
0
10
5
−5
0.3
0.2
0.1

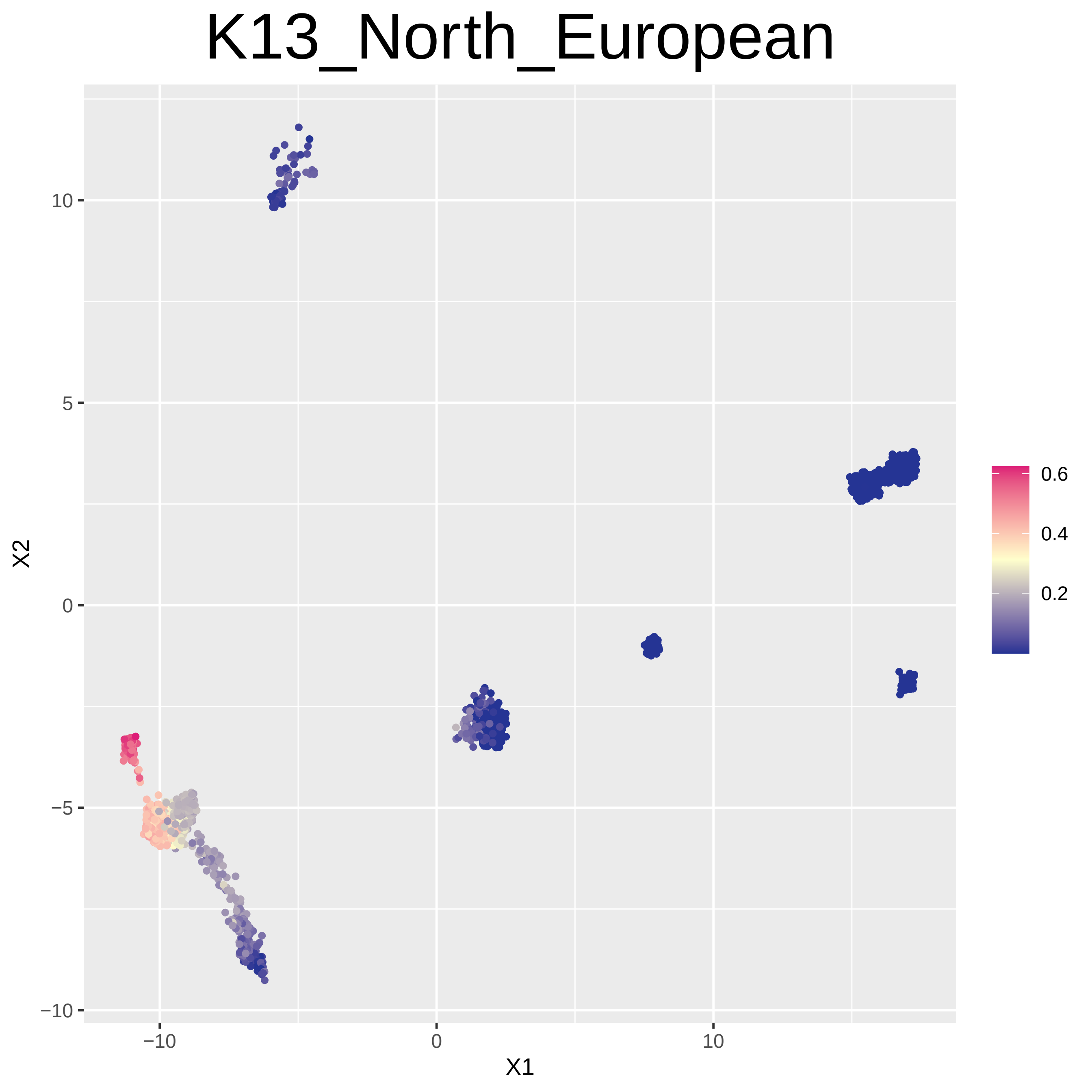

K13_North_European
X1
X2
−10
0
10
10
5
0
−5
−10
0.6
0.4
0.2

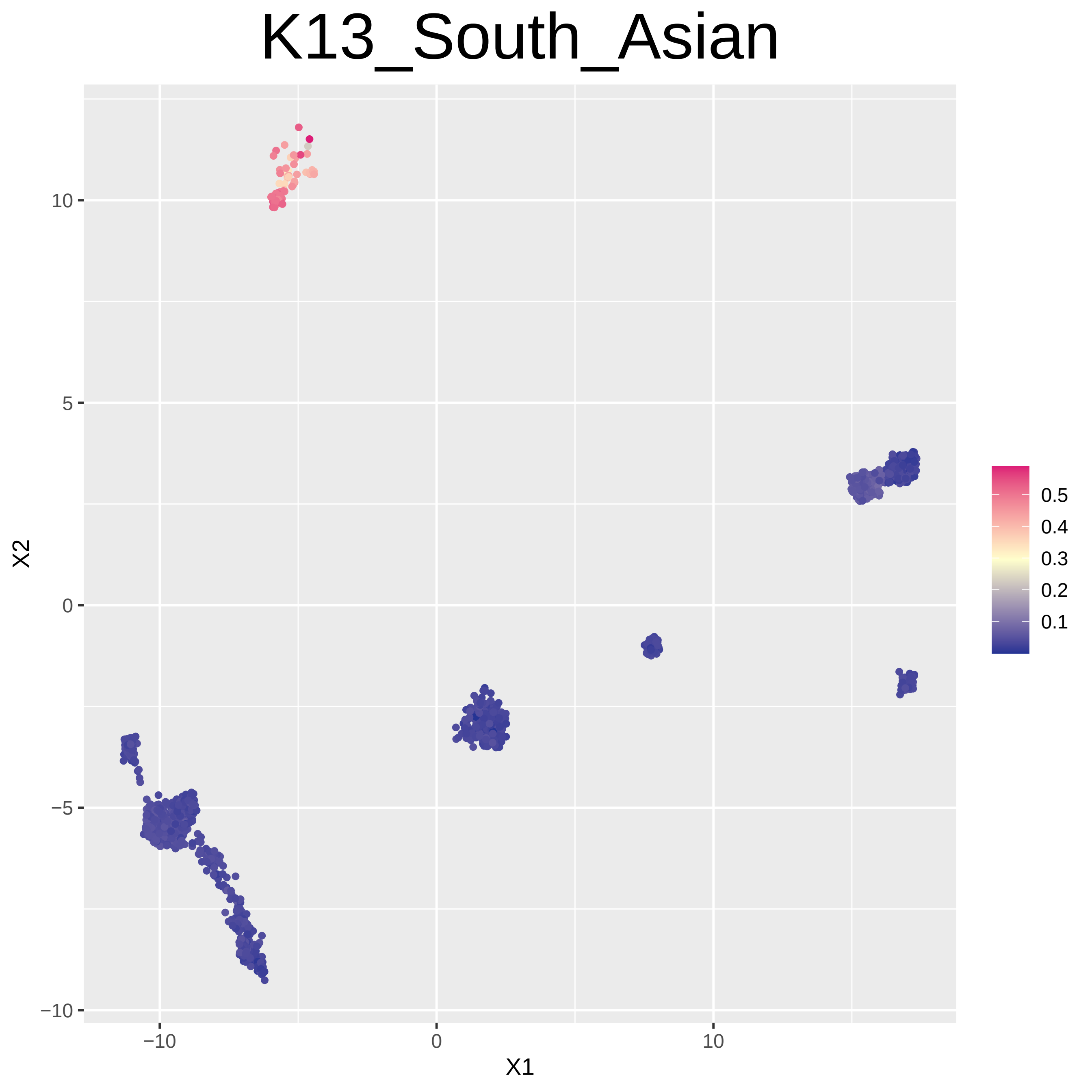

K13_South_Asian
X1
X2
10
5
0
−5
−10
0
10
0.5
0.4
0.3
0.2
0.1

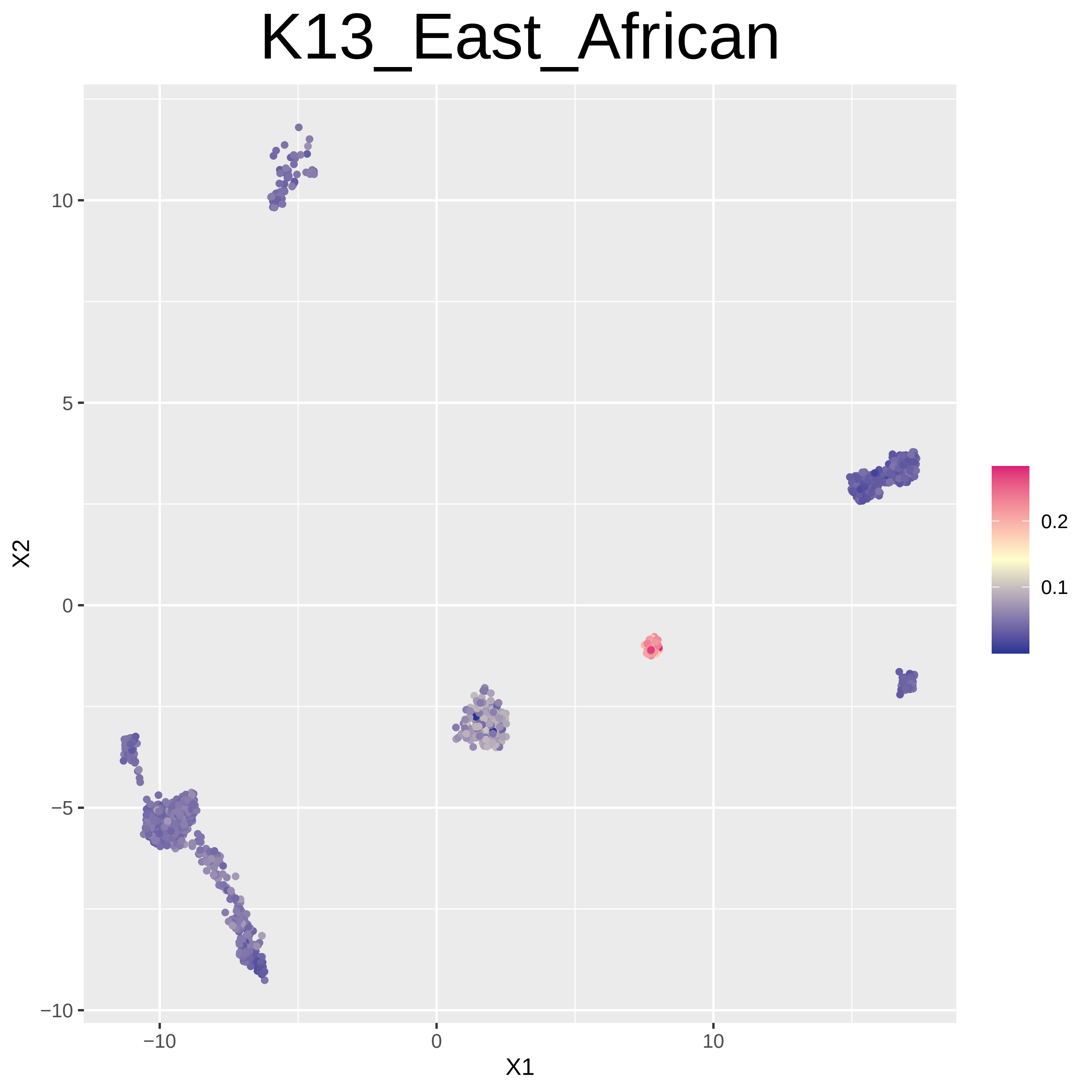
K13_East_African
X1
X2
−10
0
10
−10
−5
0
5
10
0.1
0.2

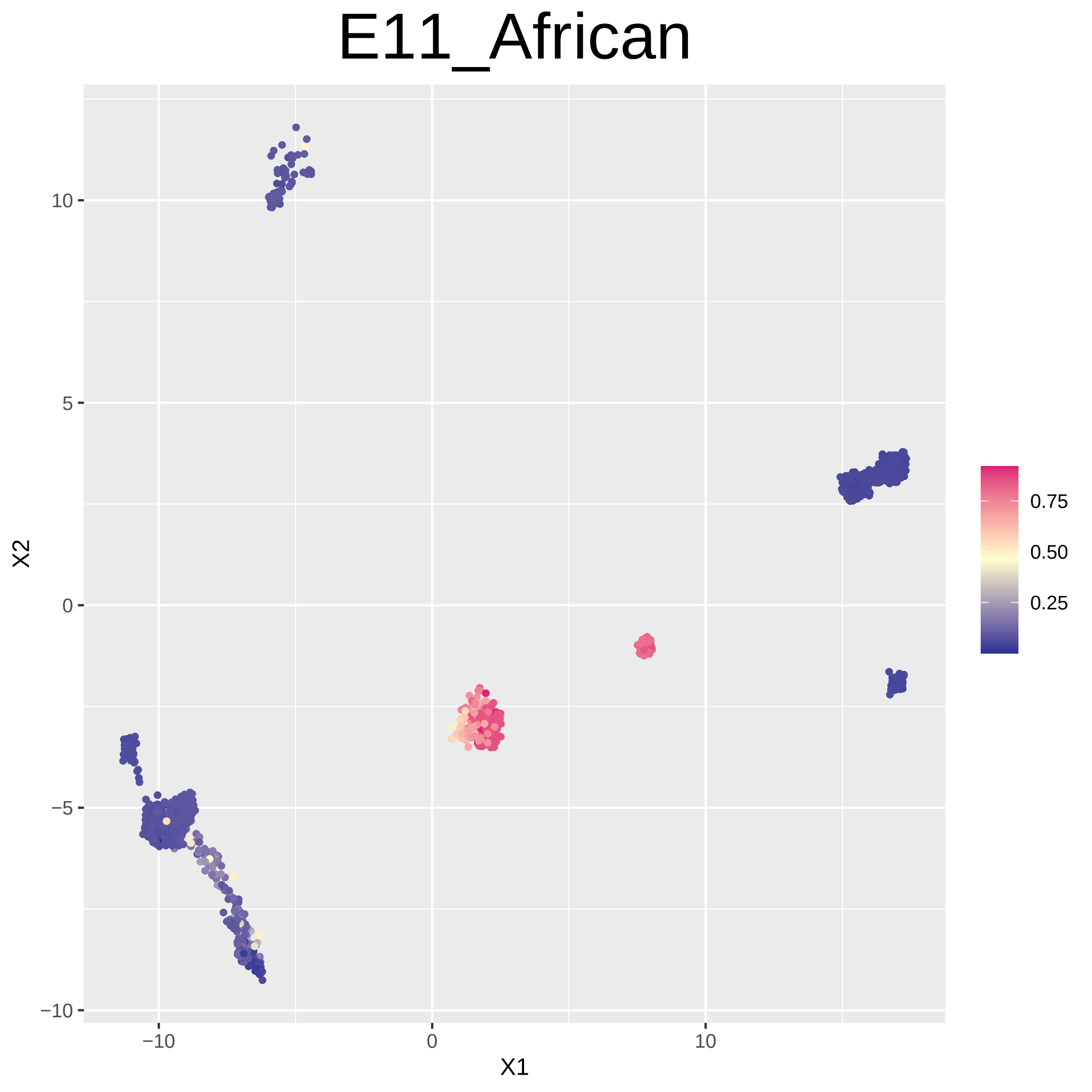

E11_African
X2
X1
10
5
0
−5
−10
−10
0
10
0.75
0.50
0.25

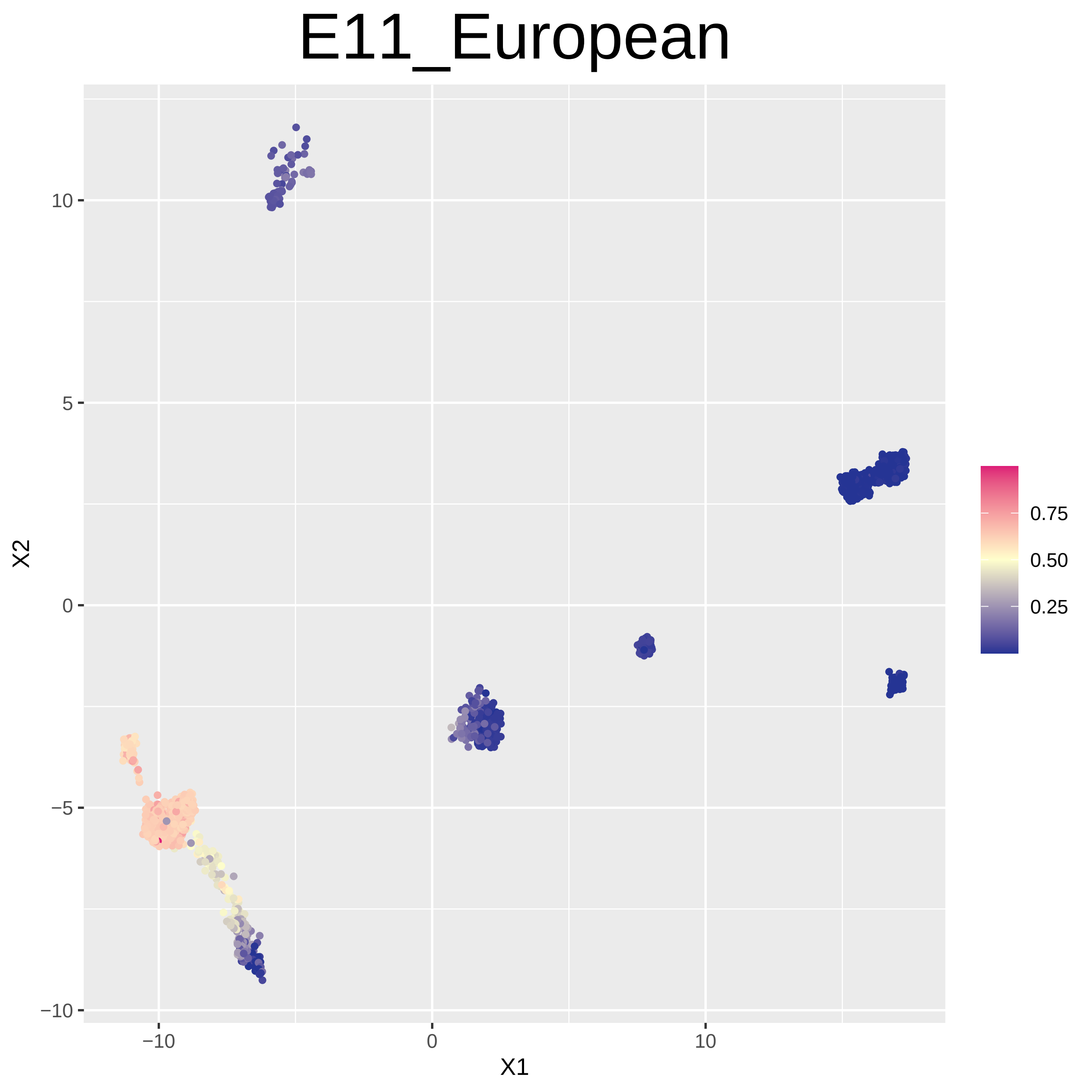
E11_European
X2
X1
10
5
0
−5
−10
−10
0
10
0.75
0.50
0.25

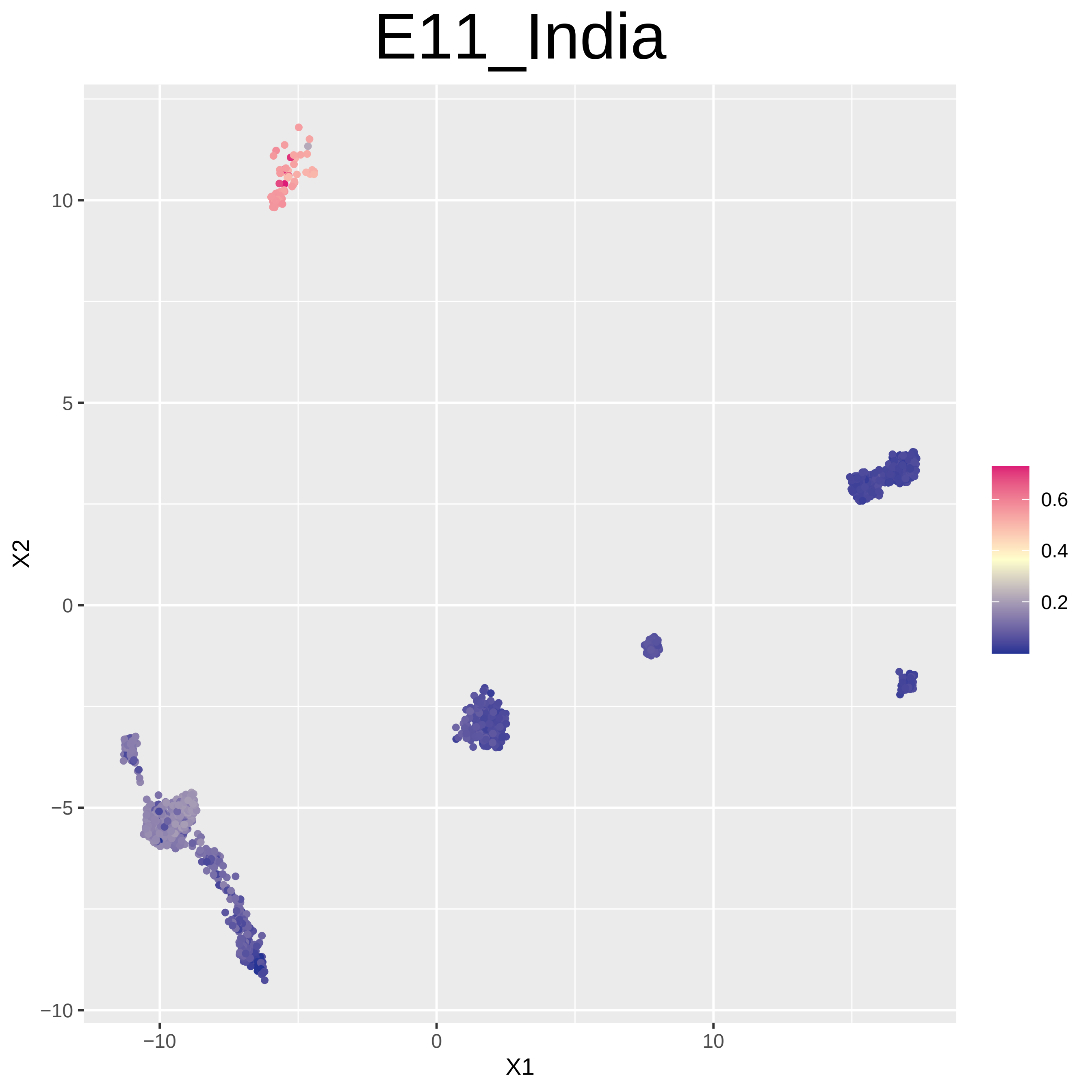

E11_India
X1
X2
−10
0
10
−10
−5
0
5
10
0.6
0.4
0.2

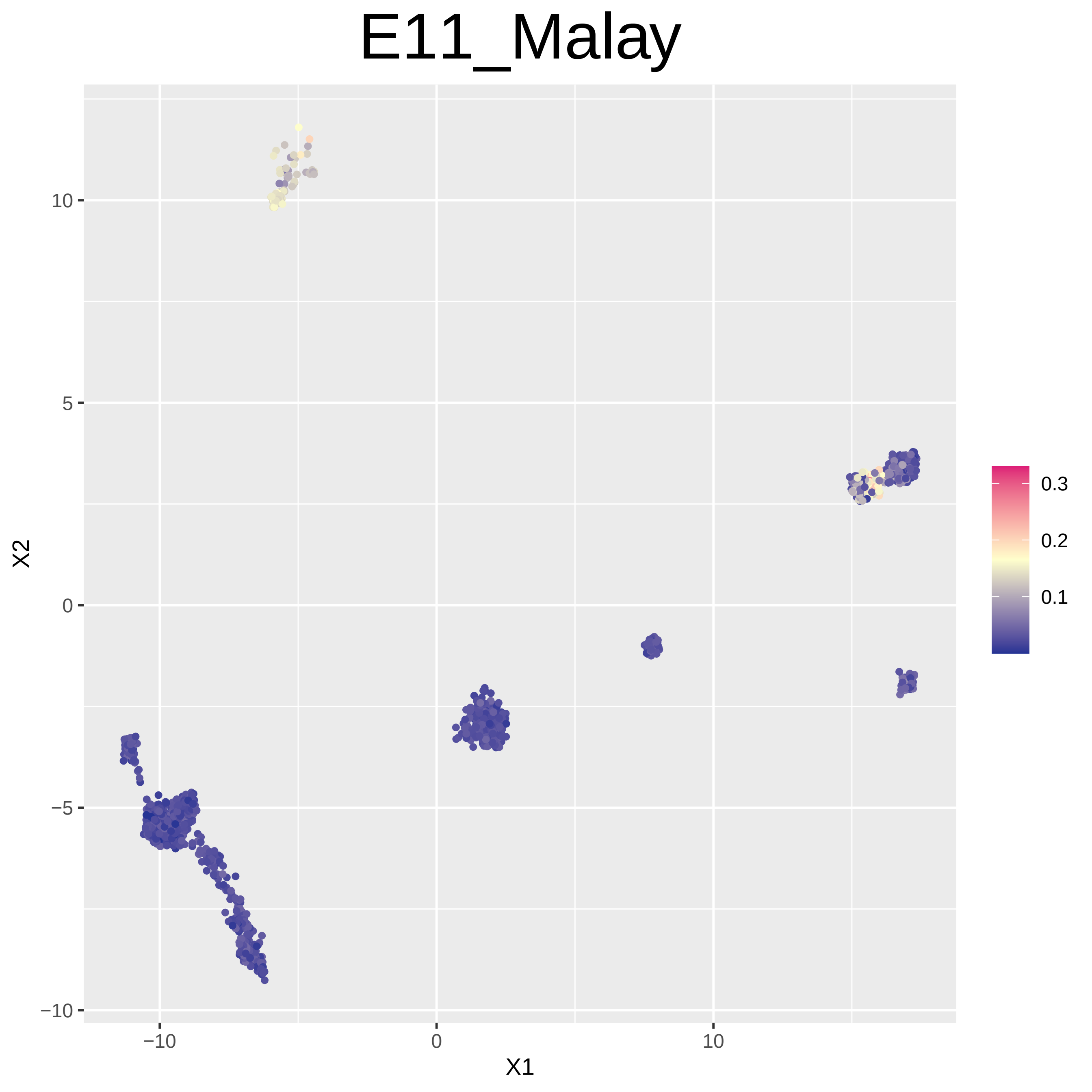
E11_Malay
X1
X2
−10
0
10
5
−5
0.1
0.2
0.3

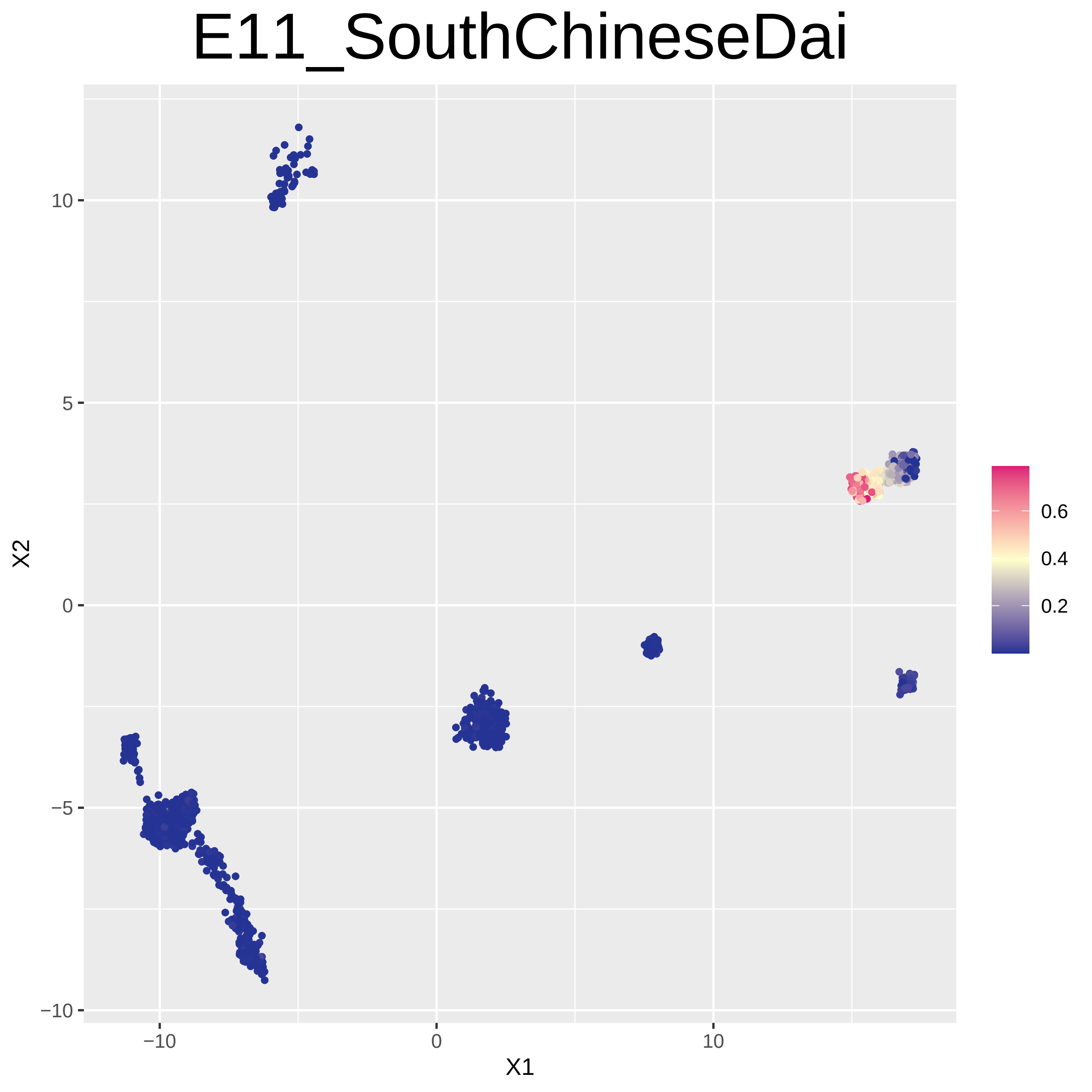

E11_SouthChineseDai
X2
X1
10
5
0
−5
−10
−10
0
10
0.6
0.4
0.2

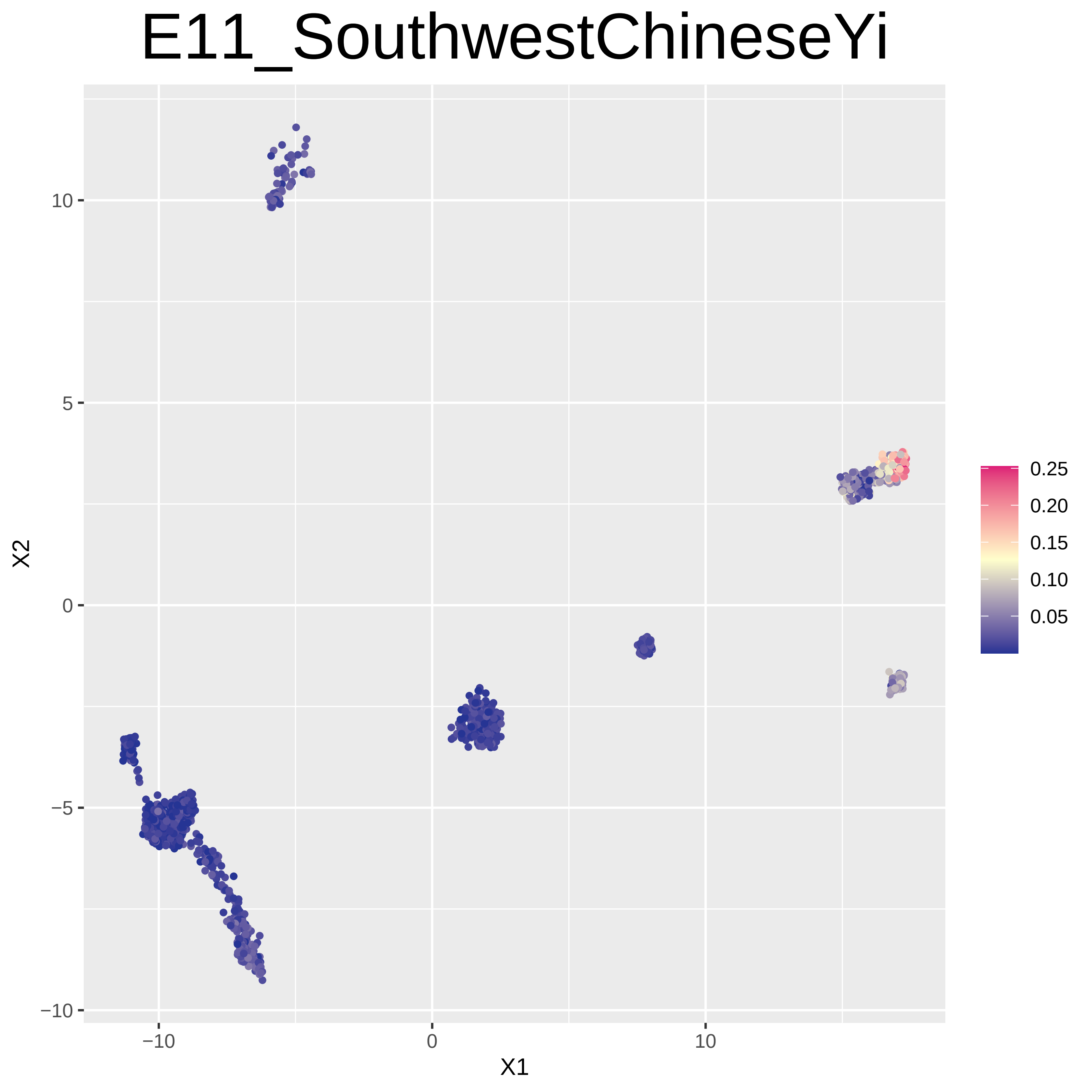
E11_SouthwestChineseYi
X1
X2
−10
0
10
10
5
0
−5
−10
0.25
0.20
0.15
0.10
0.05

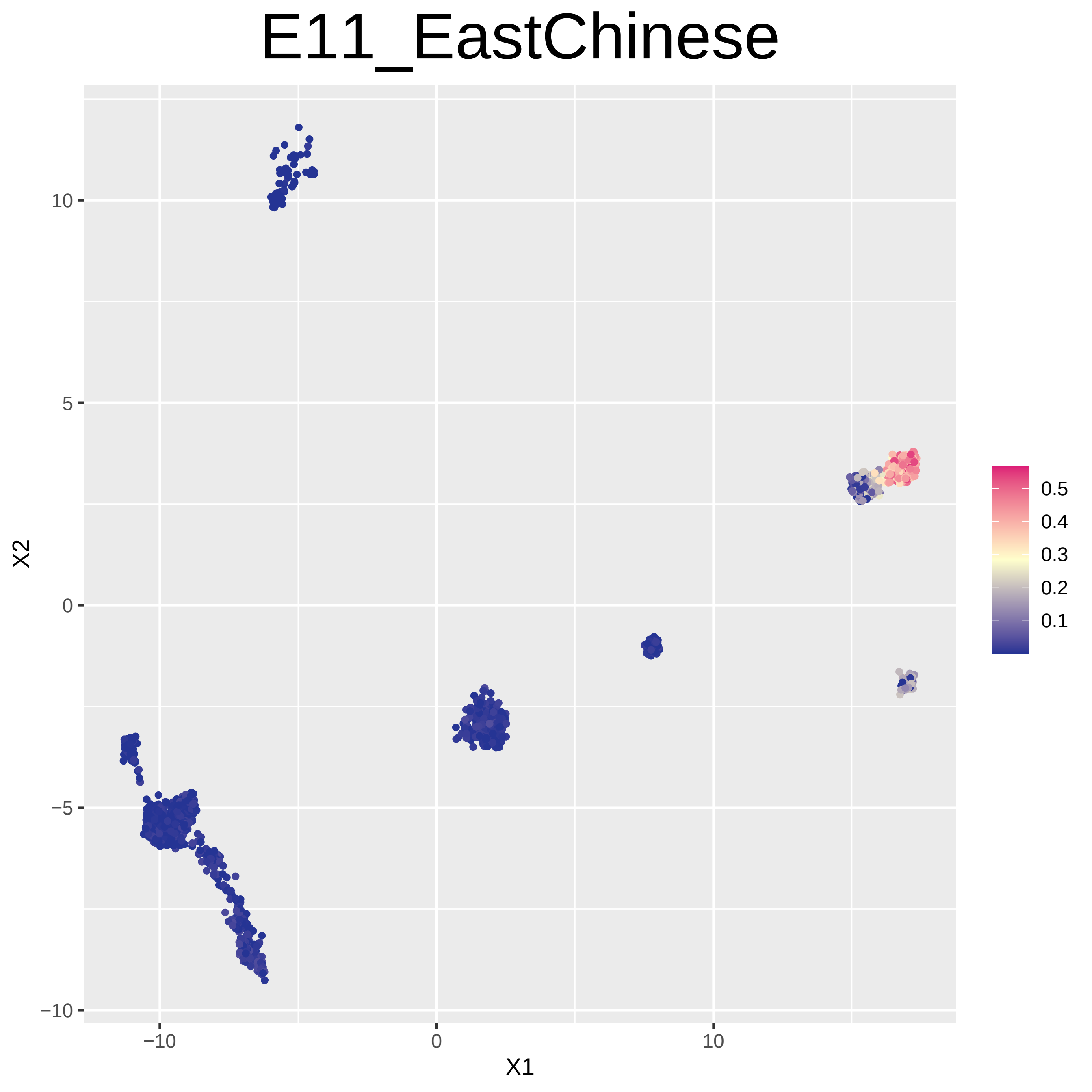

E11_EastChinese
X2
X1
10
5
0
−5
−10
−10
0
10
0.5
0.4
0.3
0.2
0.1

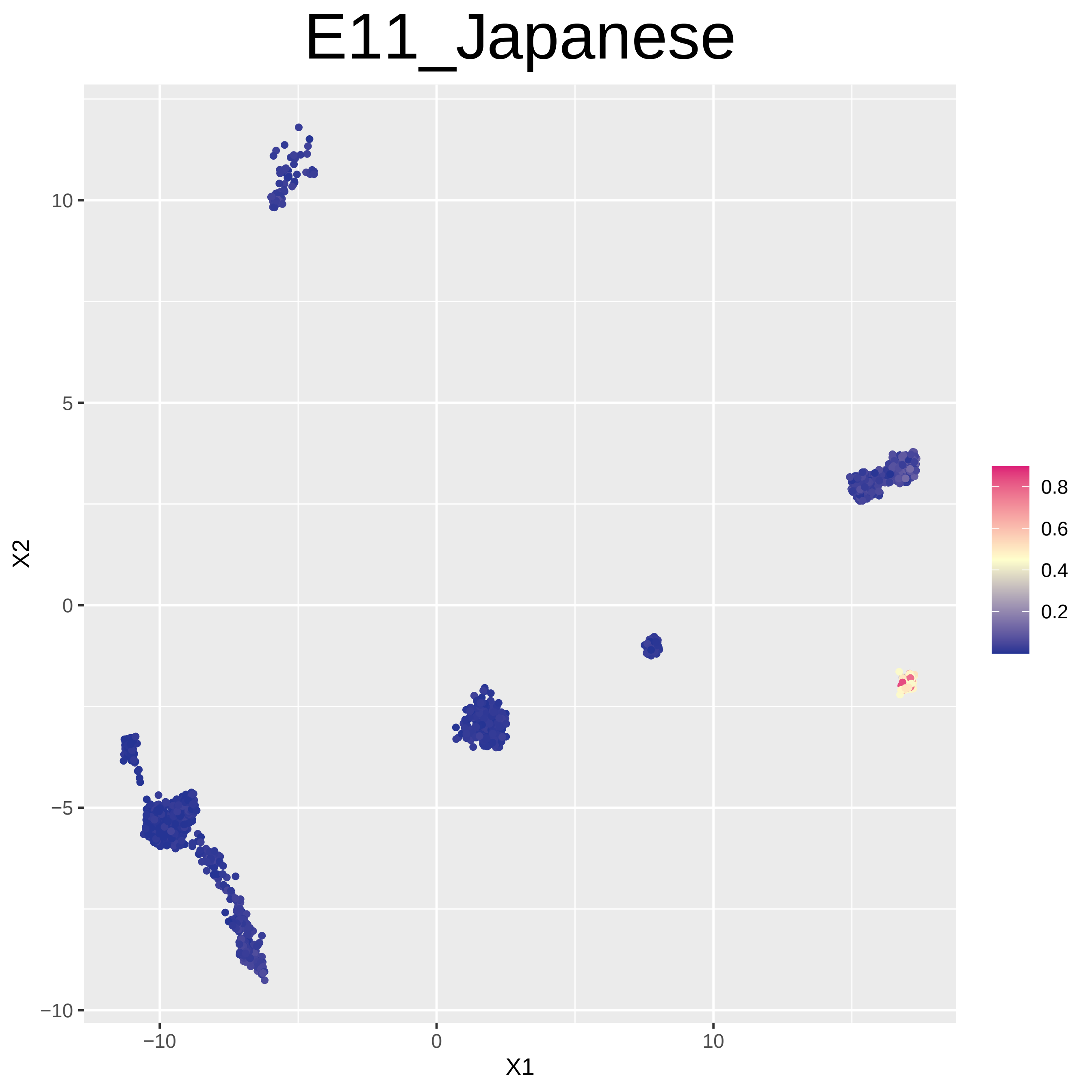
E11_Japanese
X1
X2
−10
0
10
5
−5
0.8
0.6
0.4
0.2

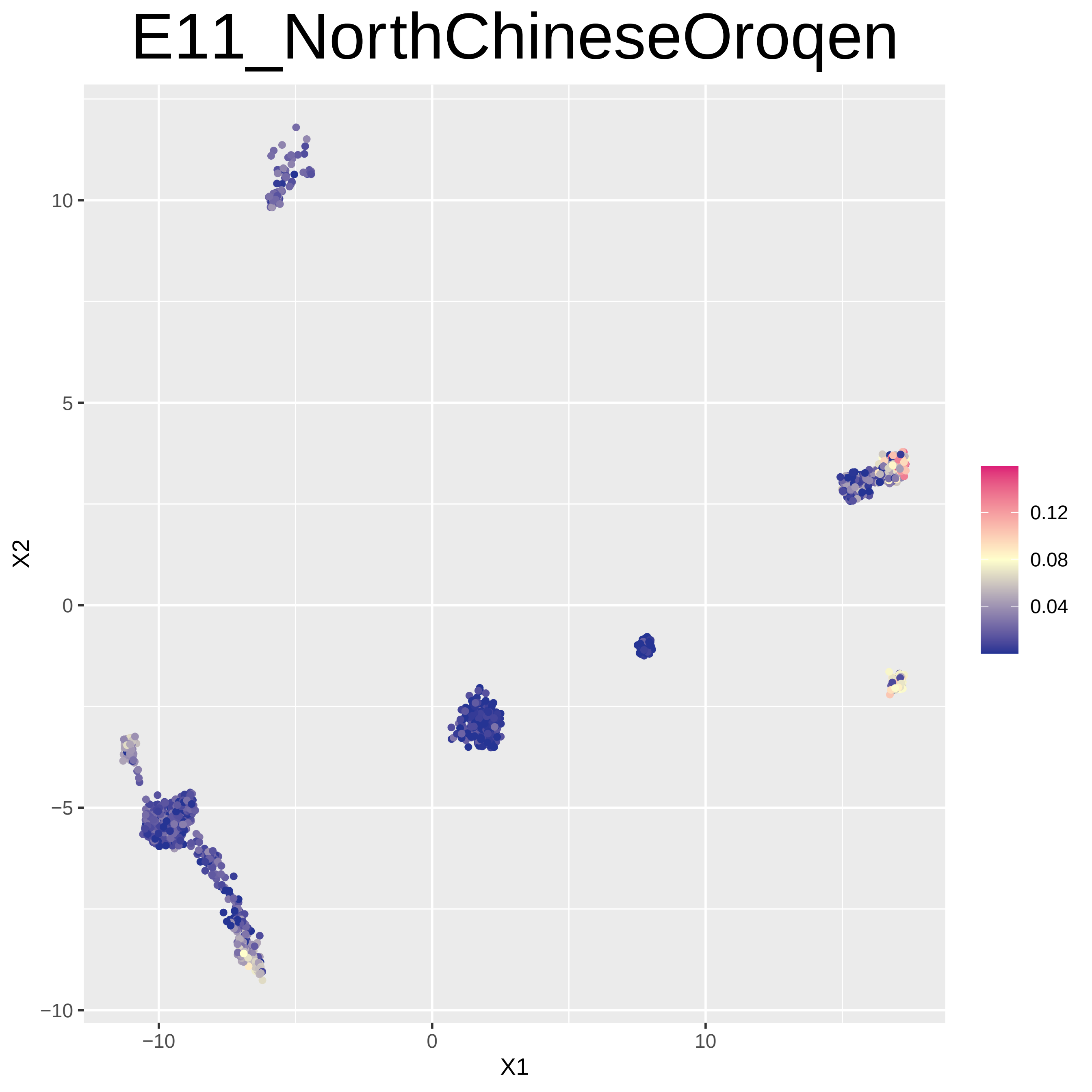

E11_NorthChineseOroqen
X1
X2
−10
0
10
−10
−5
0
5
10
0.12
0.08
0.04

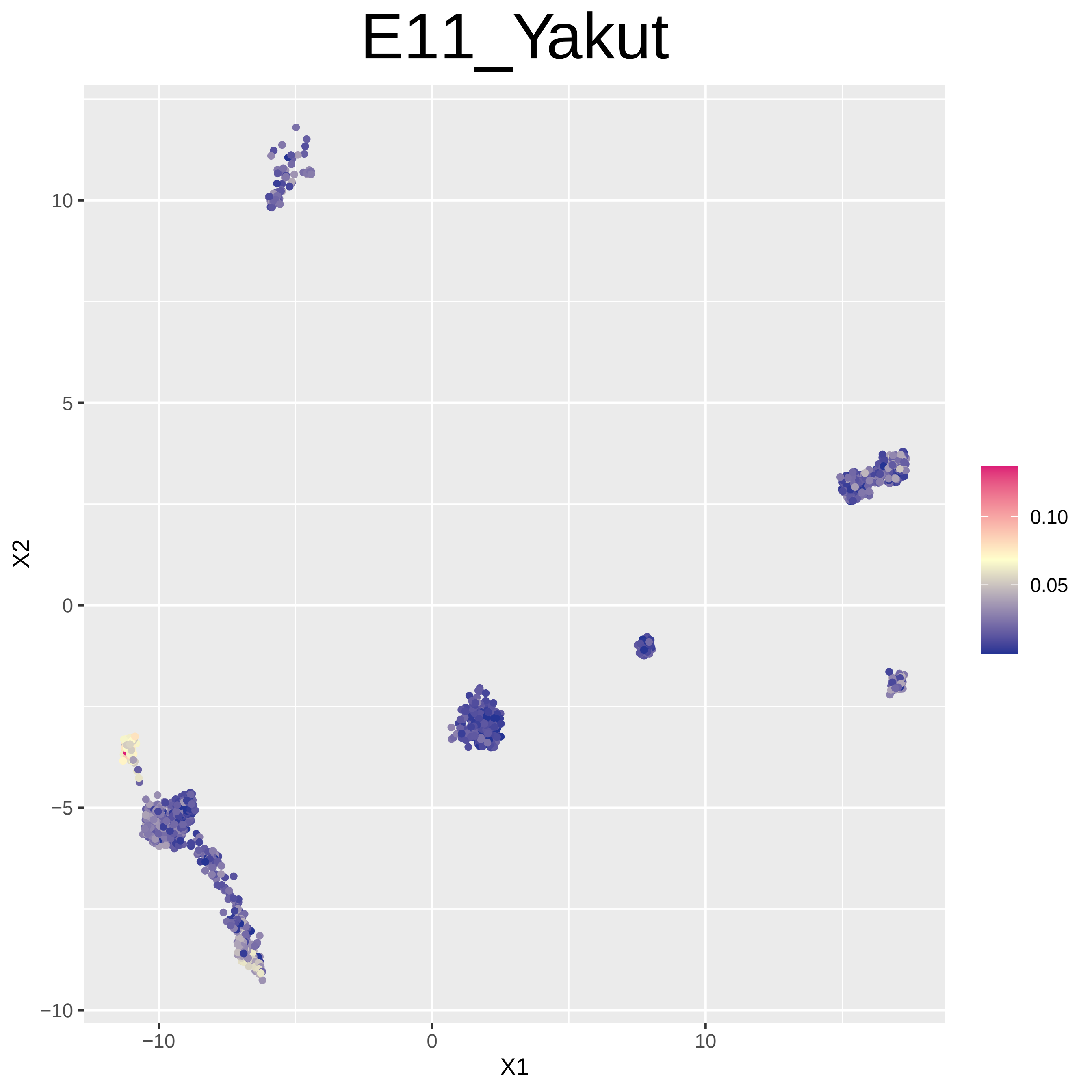
E11_Yakut
X1
X2
−10
0
10
−10
−5
0
5
10
0.10
0.05

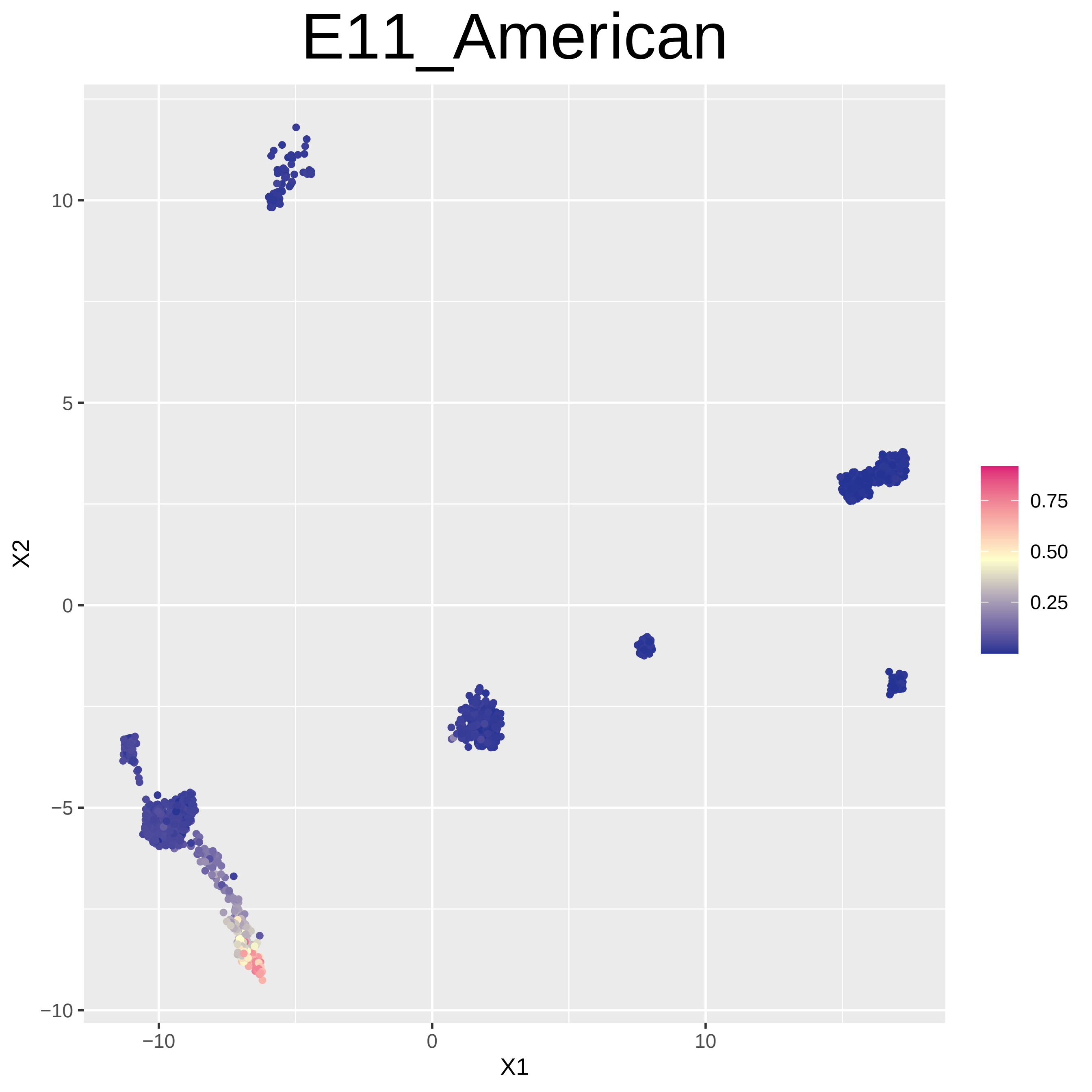
E11_American
X1
X2
−10
0
10
−10
−5
0
5
10
0.75
0.50
0.25

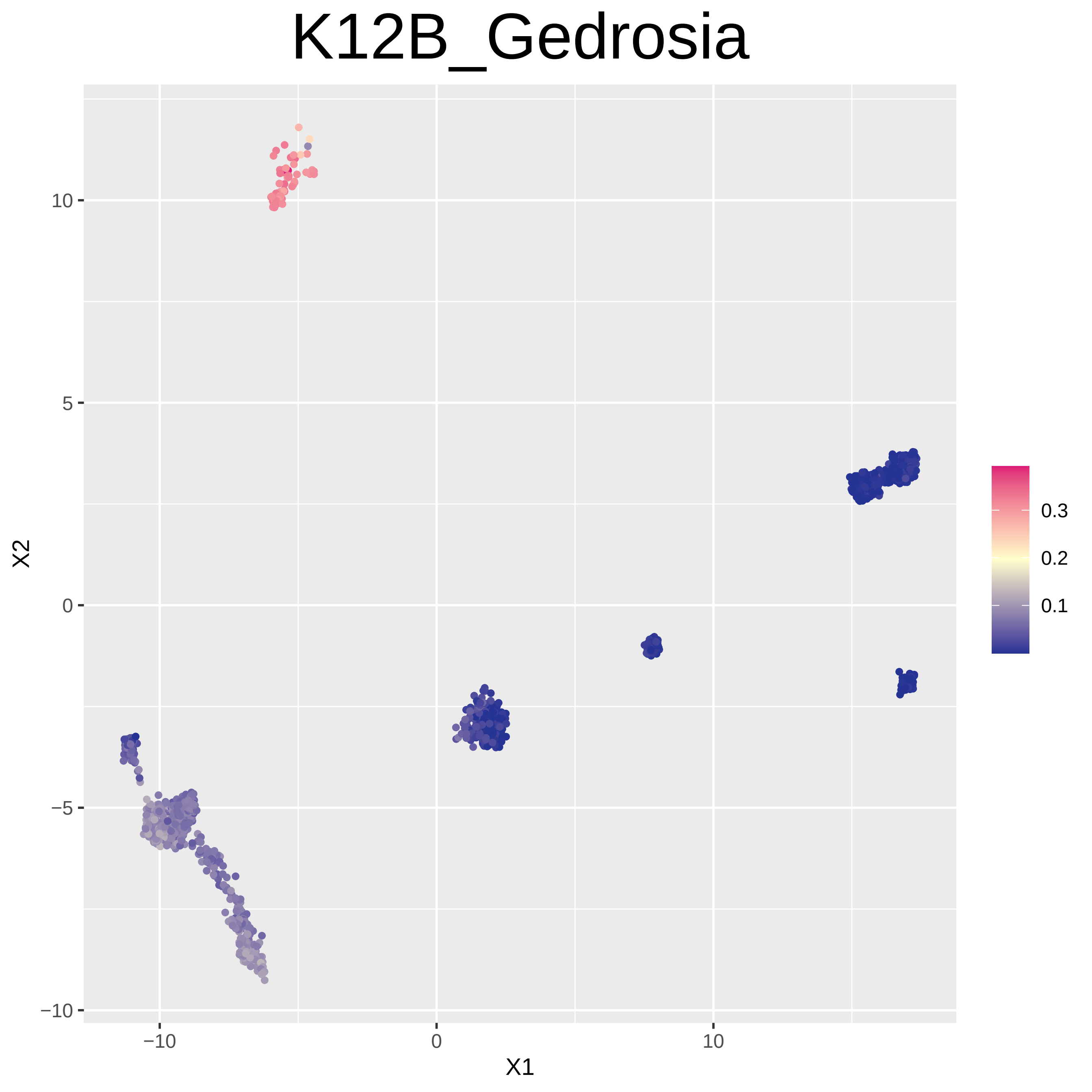
K12B_Gedrosia
X1
X2
−10
0
10
−10
−5
0
5
10
0.3
0.2
0.1

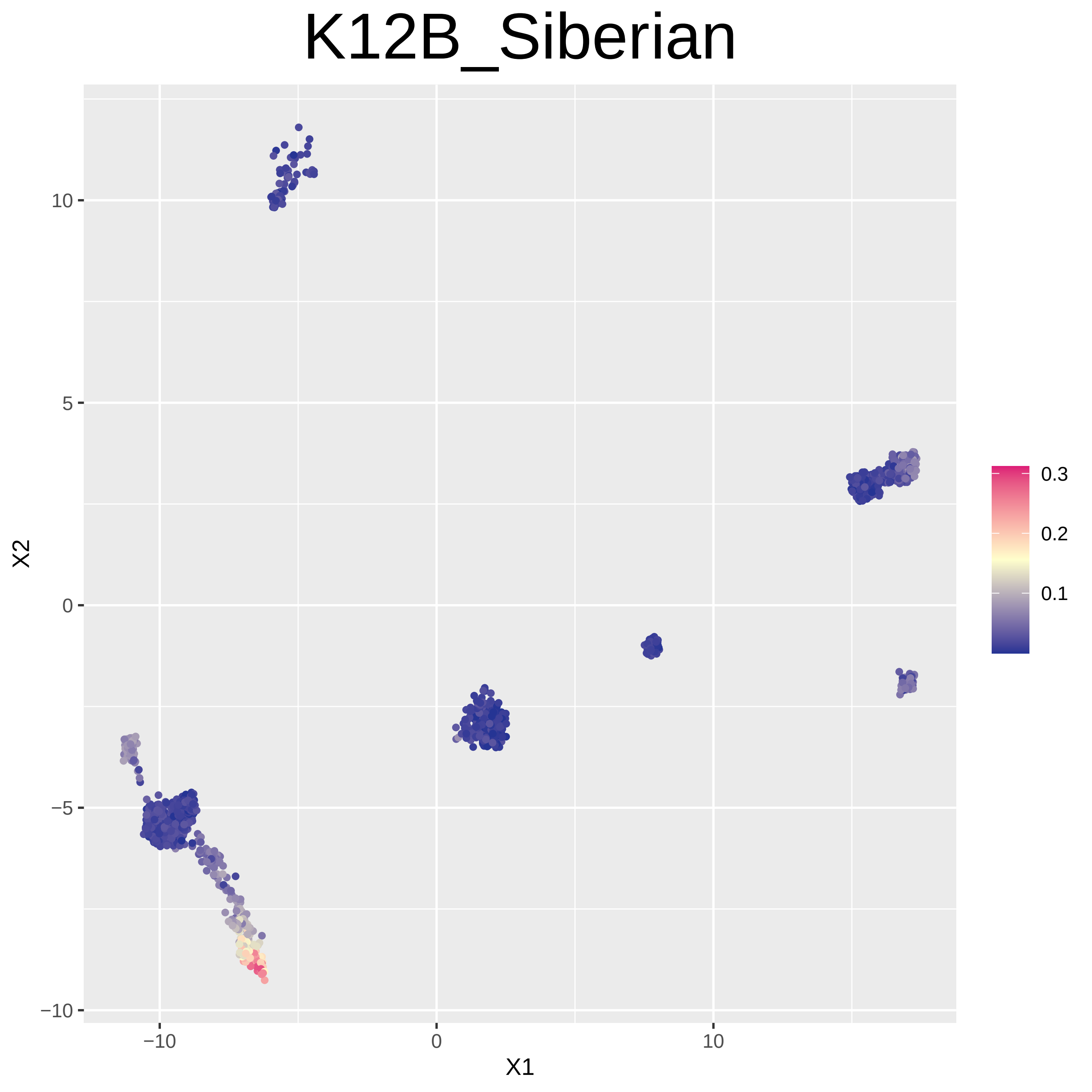

K12B_Siberian
X1
X2
−10
0
10
−10
−5
0
5
10
0.1
0.2
0.3

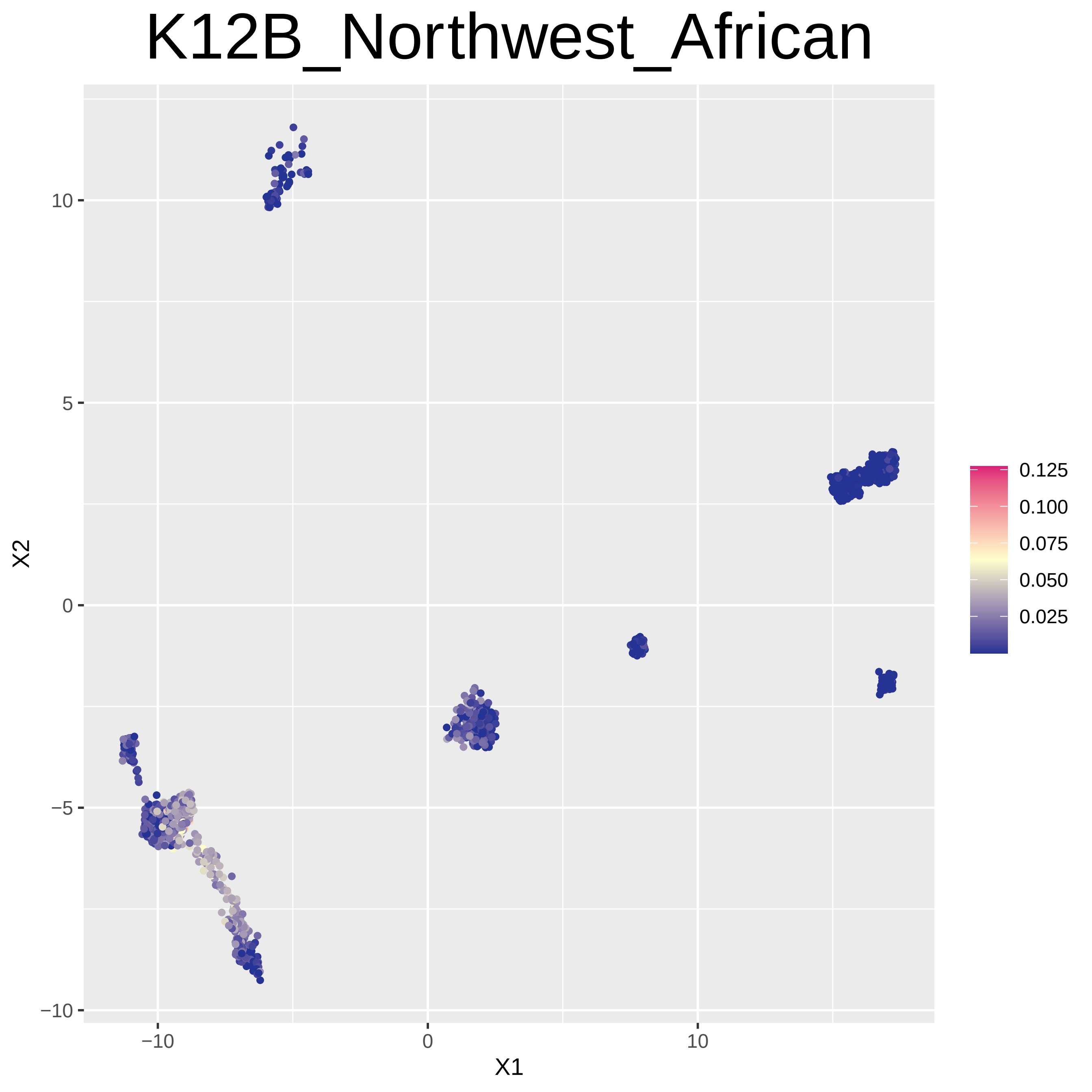
K12B_Northwest_African
X1
X2
−10
0
10
−10
−5
0
5
10
0.125
0.100
0.075
0.050
0.025

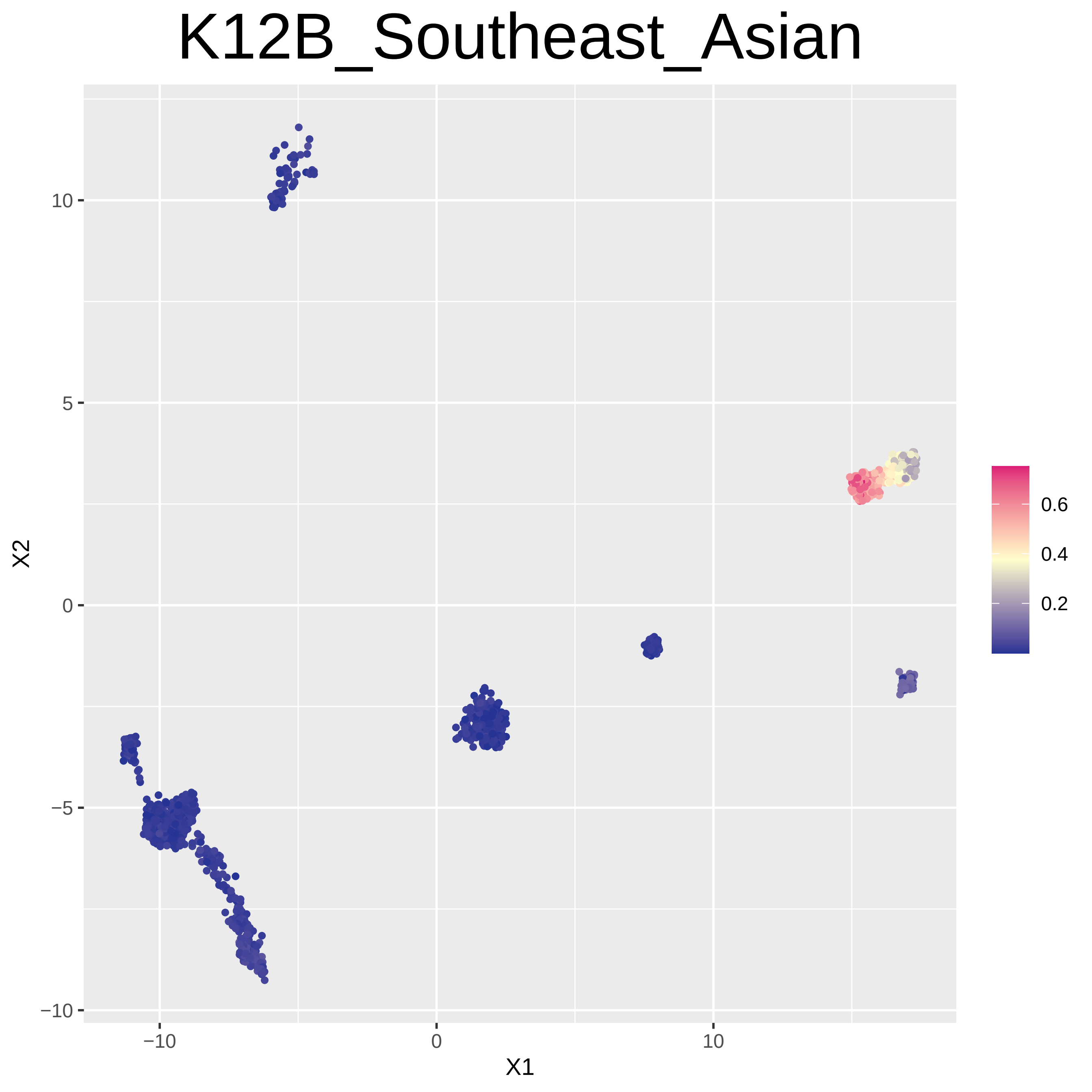
K12B_Southeast_Asian
X1
X2
−10
0
10
5
−5
0.6
0.4
0.2

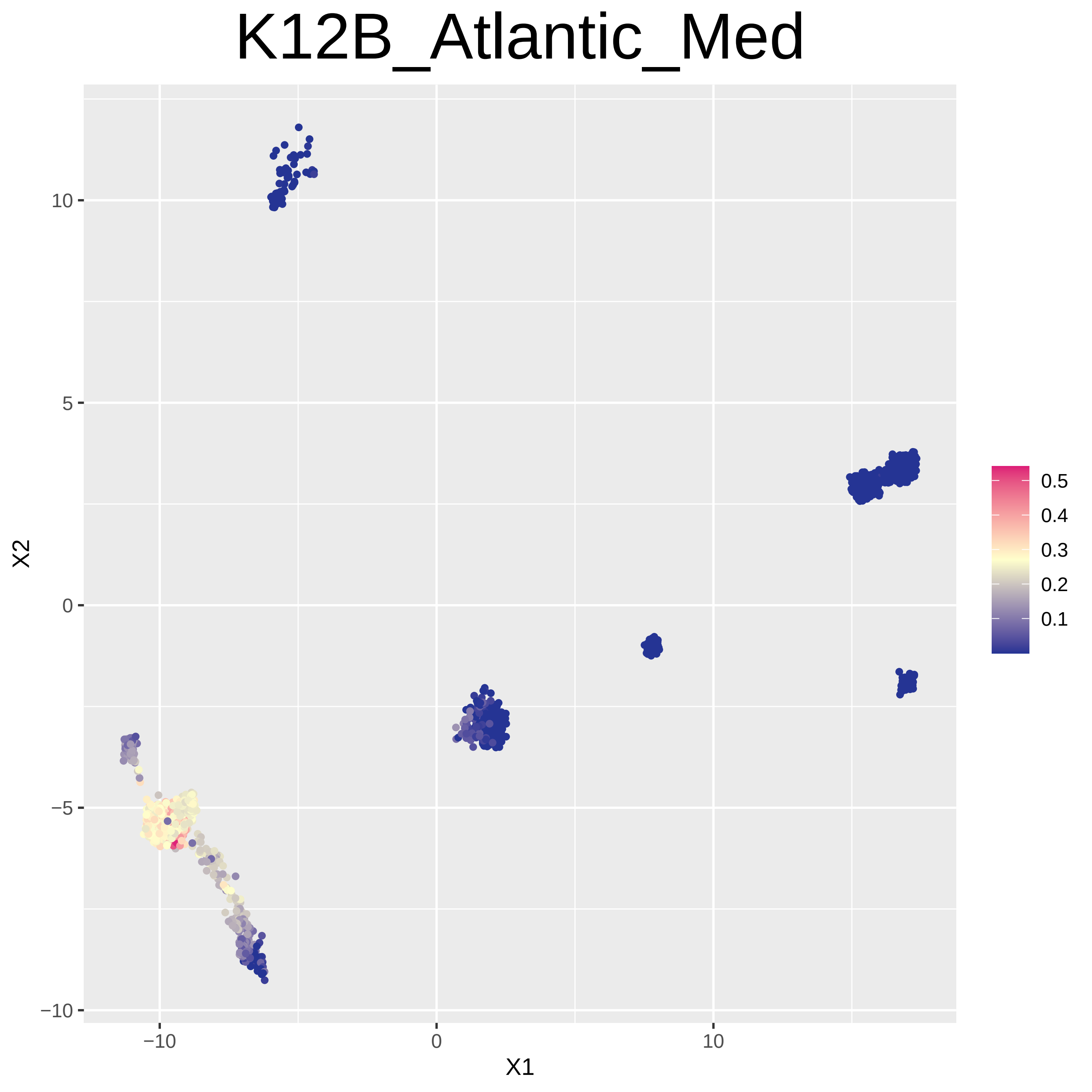

K12B_Atlantic_Med
X2
X1
10
5
0
−5
−10
−10
0
10
0.5
0.4
0.3
0.2
0.1

### K12B_North_European

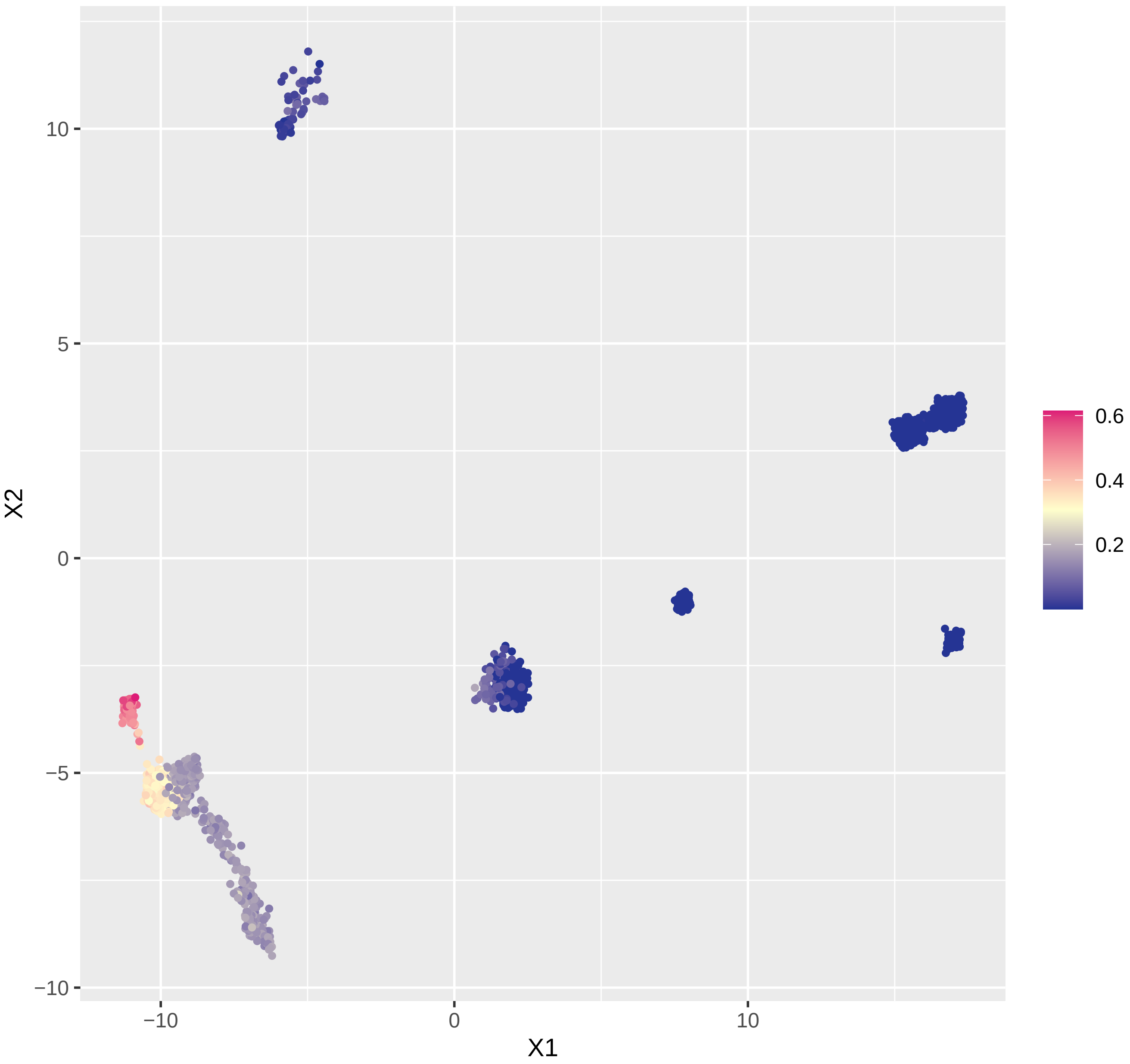

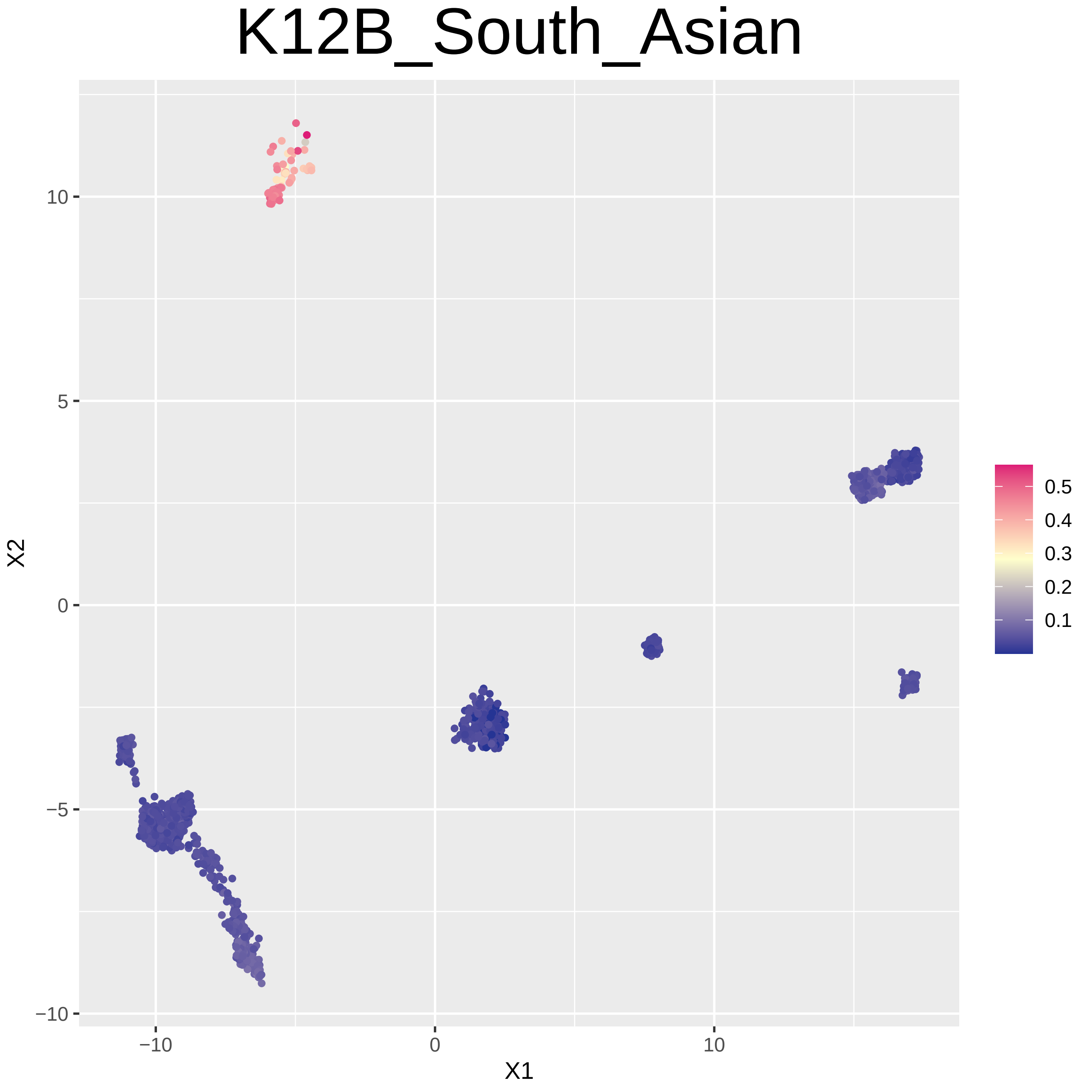

K12B_South_Asian
X2
10
5
0
−5
−10
X1
−10
0
10
0.5
0.4
0.3
0.2
0.1

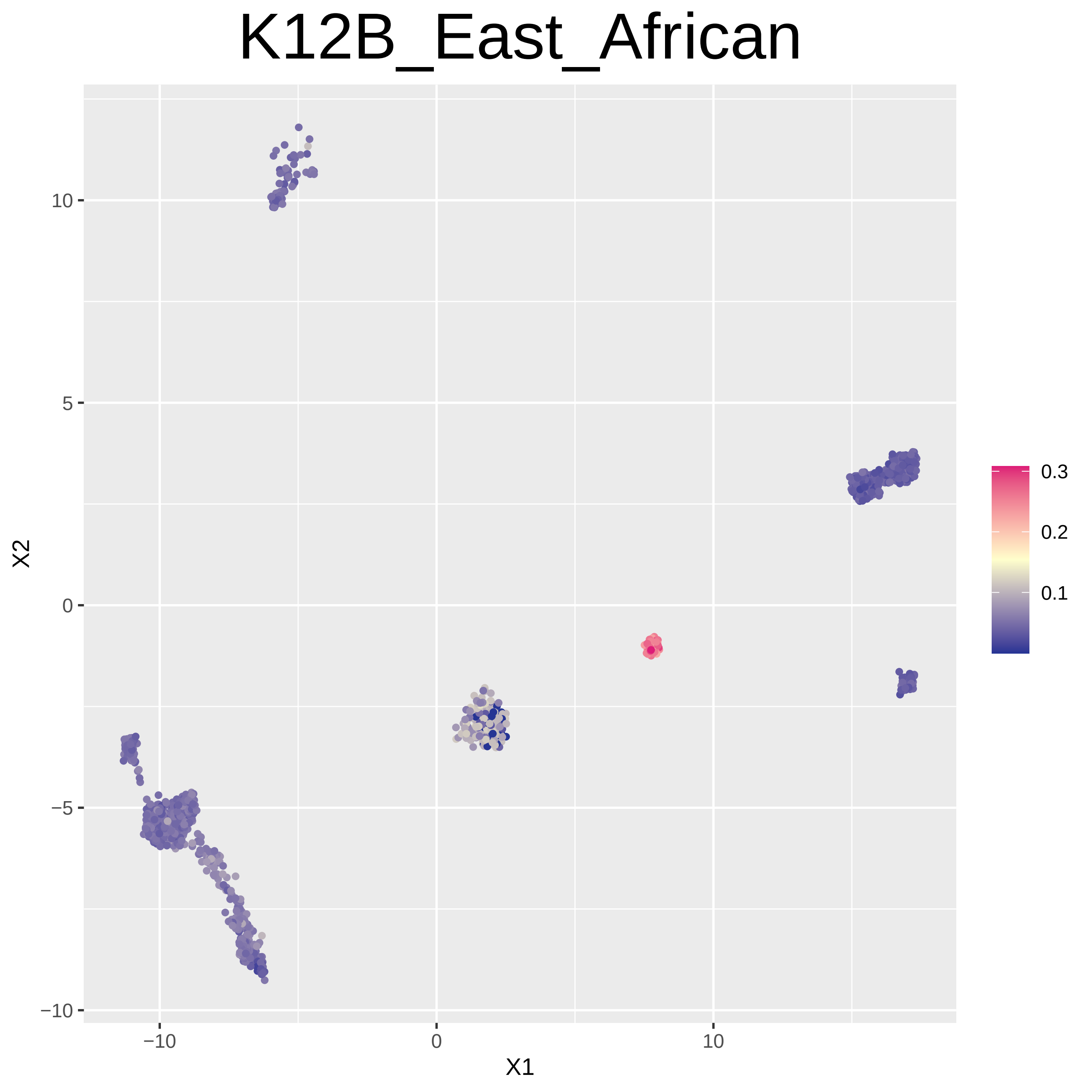
K12B_East_African
X2
X1
10
5
0
−5
−10
−10
0
10
0.3
0.2
0.1

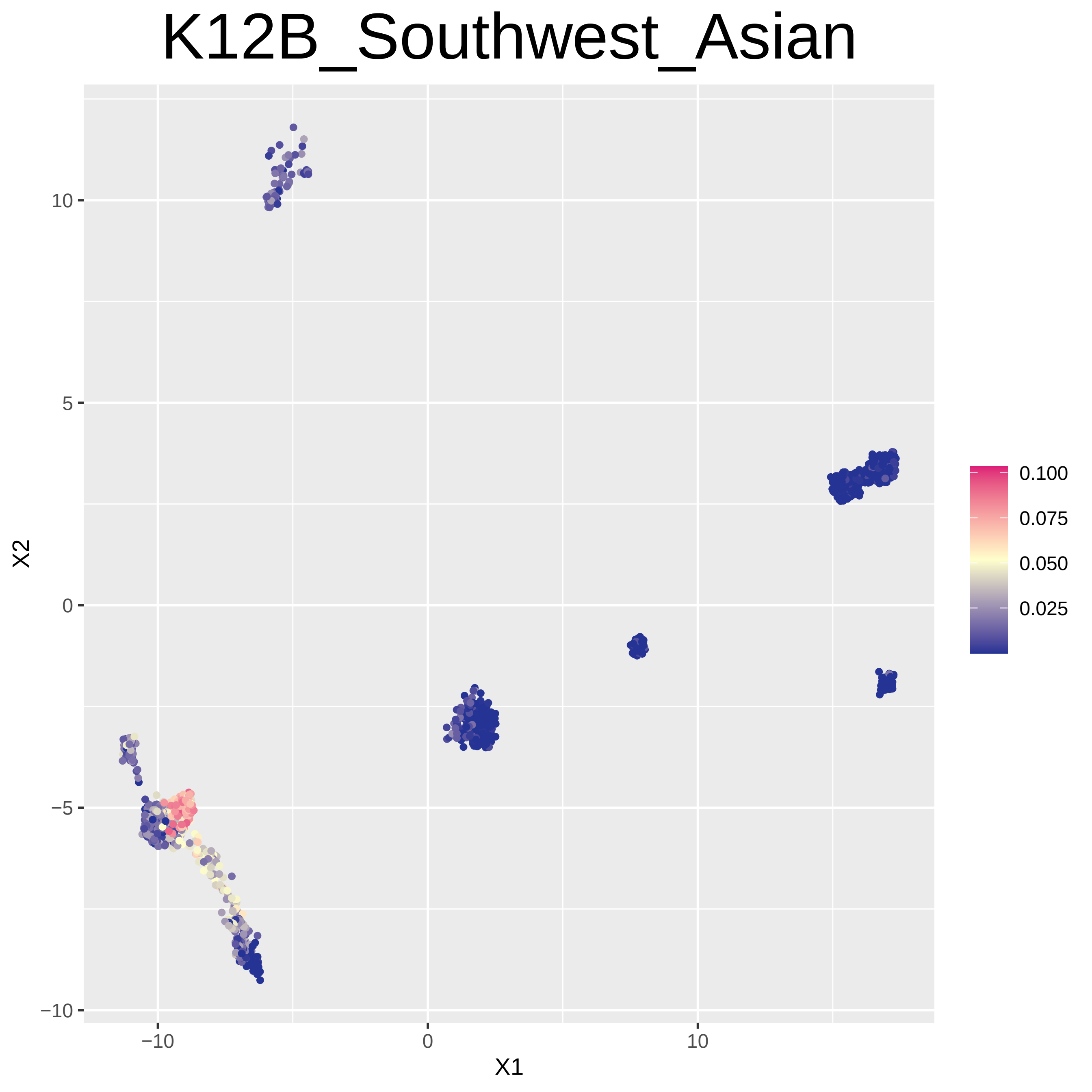
K12B_Southwest_Asian
X2
X1
10
5
0
−5
−10
−10
0
10
0.100
0.075
0.050
0.025

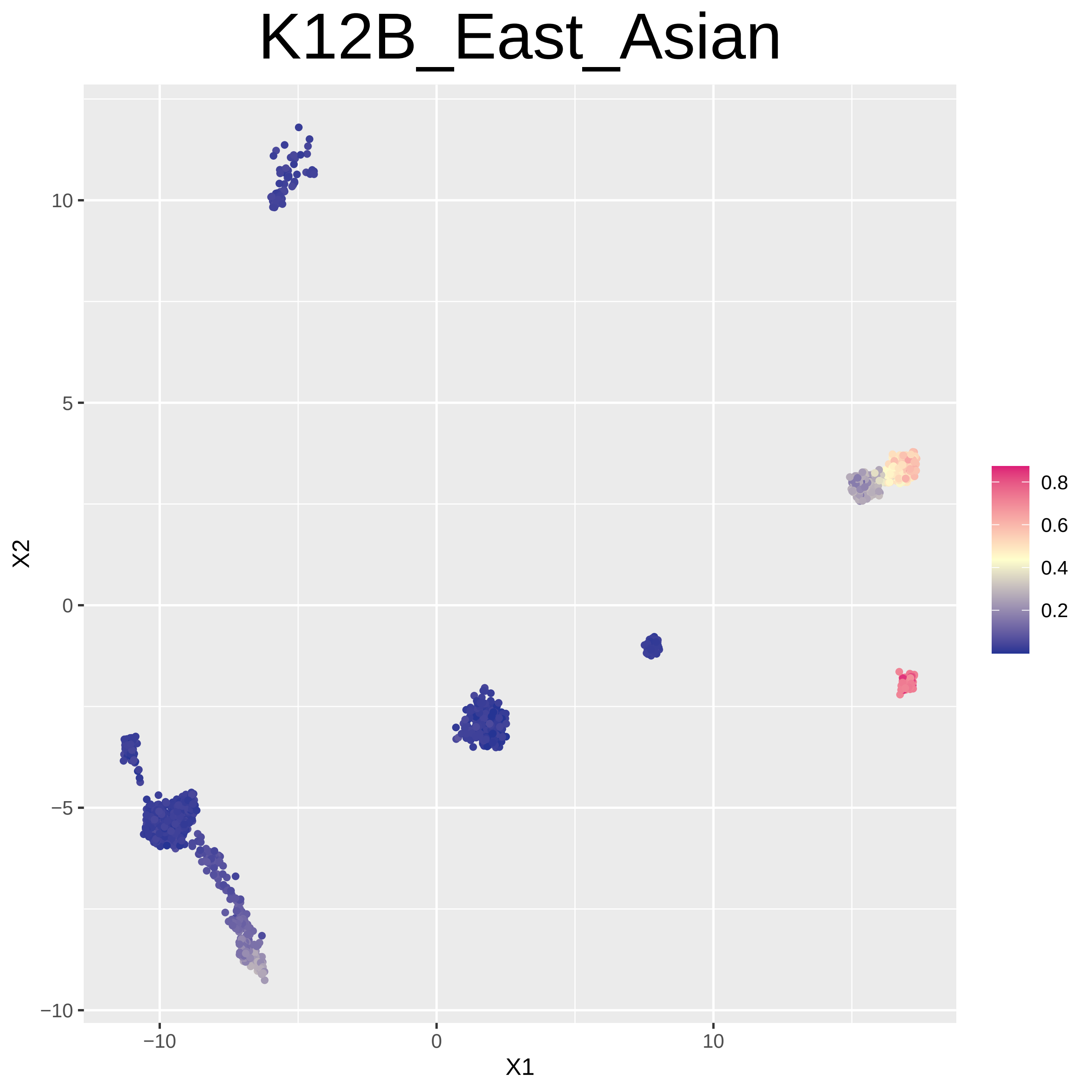
K12B_East_Asian
X1
X2
−10
0
10
5
−5
0.2
0.4
0.6
0.8

### K12B_Caucasus

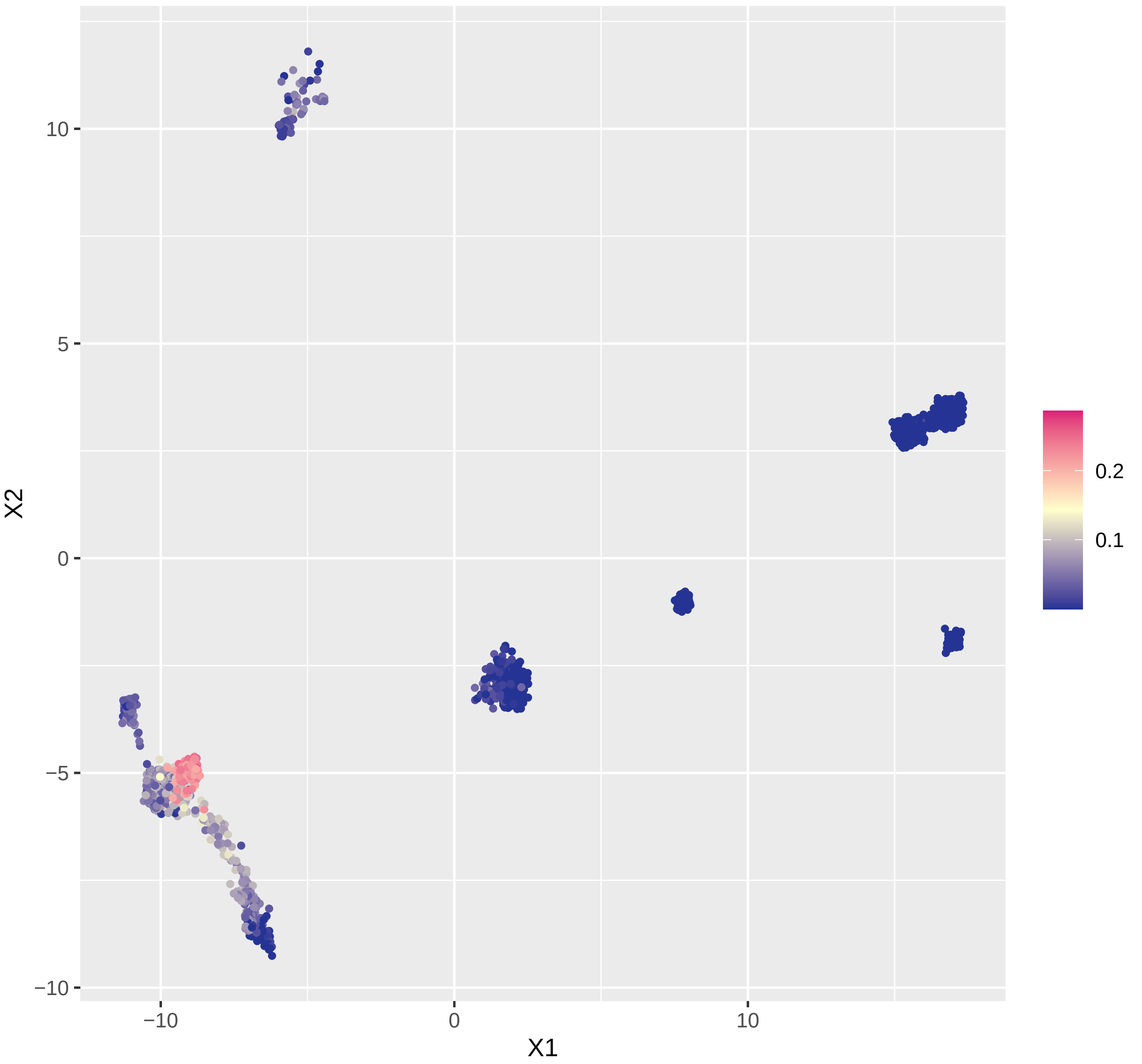

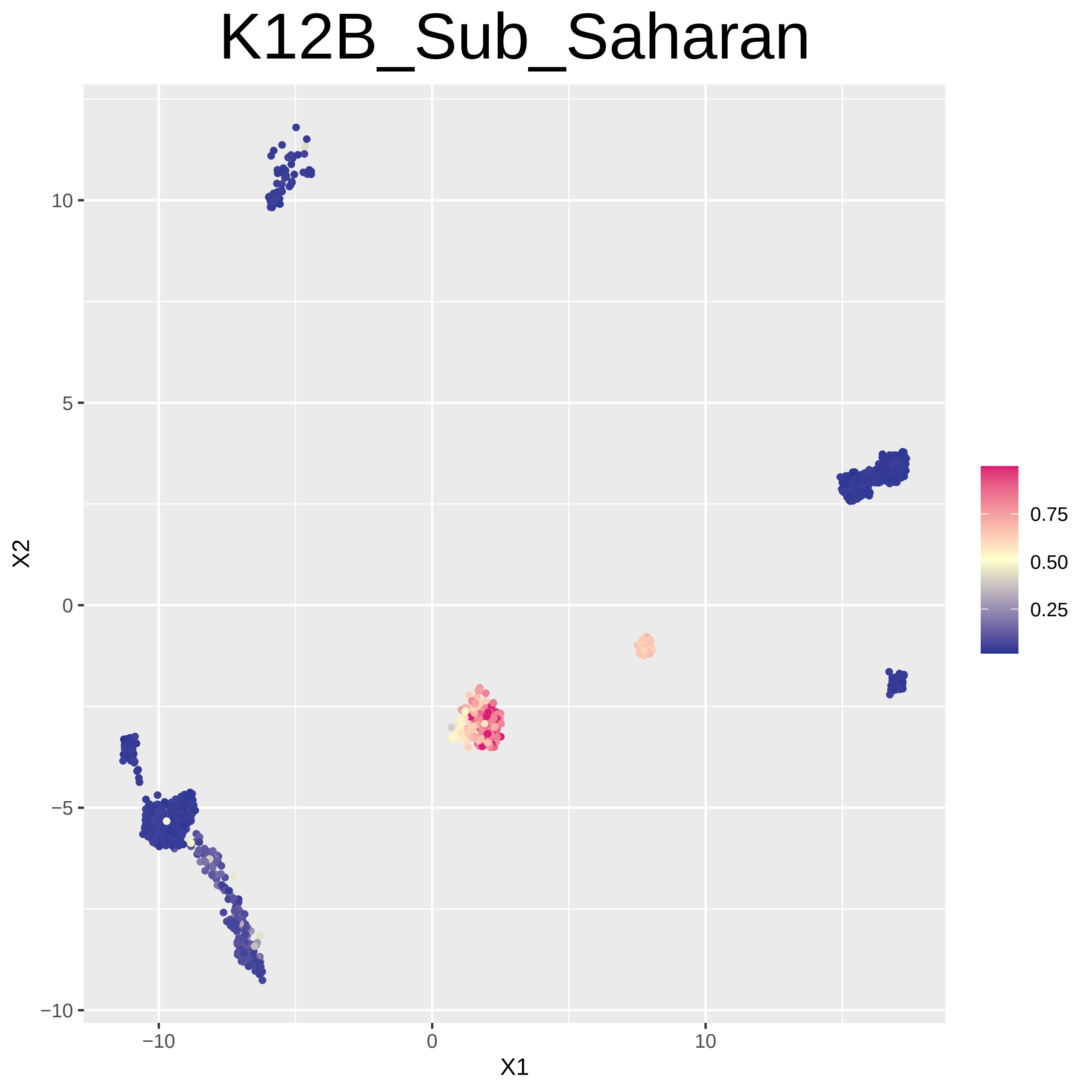
K12B_Sub_Saharan
X1
X2
−10
0
10
−10
−5
0
5
10
0.75
0.50
0.25
